# Inter- and intra-tumor heterogeneous immune presentation in primary uterine carcinosarcoma determined by digital spatial profiling

**DOI:** 10.1101/2024.09.05.611350

**Authors:** Ying-Xuan Li, Wei-Chou Lin, Ko-Chen Chen, Wen-Chun Chang, Chin-Jui Wu, Lin-Hung Wei, Ruby Yun-Ju Huang, Bor-Ching Sheu

**Author notes:** **Corresponding Author:** Bor-Ching Sheu, MD, Ph.D., **E-mail:****, Mailing Address:** Department of Obstetrics and Gynecology, National Taiwan University Hospital and National Taiwan University College of Medicine, 15F, No. 8, Chung-Shan South Road, 10002, Taipei, Taiwan. **Corresponding Author: Ruby Yun-Ju Huang, M.D., Ph.D. E-mail:**, **Mailing Address: School of Medicine, College of Medicine, National Taiwan University, No.1 Ren-Ai Road Section 1, Taipei, Taiwan**.

## Abstract

Uterine carcinosarcoma (UCS) is a rare gynecologic cancer with unfavorable prognosis. Its biphasic histologic components are known to associated with outcomes. Dissecting the tumor heterogeneity within UCS is crucial to discover molecular biomarkers for patient stratification. The aim of this study was to utilize spatial profiling technology to analyze immune presentation in different histological components within UCS tumors. The study included 23 UCS patients whose tumor sections were subjected to nanoString GeoMx Digital Spatial Profiling (DSP) to profile 30 immune-related proteins in 491 segments. We identified distinct cluster-labeled subtypes showing varied immune marker expressions. The immune presentations within the carcinoma and sarcoma components showed significant differences. Markers indicating M2 macrophages (CD45+CD163+) and epithelial-mesenchymal transition (EMT) (SMA, fibronectin) were highly expressed in the sarcoma components. The immune presentation subtypes within each histological component also differed between patients with recurrence in 5 years and without recurrence over 5 years. The cluster-labeled subtype with high Neutrophil/PMN-MDSC marker expression (CD66b) was associated with longer survival, whereas the subtype with high NK cell marker expressions (CD56) was linked to poor prognosis. Our result suggested that immune features within the carcinoma component of UCS tumors would determine patient outcomes and should be considered as a strategy for patient stratification.

## Introduction

Uterine carcinosarcoma (UCS) is a rare endometrial cancer, accounting for about 5% of uterine malignancy, and the incidence has increased gradually. The prognosis of UCS is poor, responsible for 15% of deaths from uterine malignancy^1-3^. UCS exhibits a biphasic histological feature characterized by the presence of both epithelial and mesenchymal elements. The epithelial element is usually high-grade in nature, such as high-grade endometrioid or serous carcinoma, while the mesenchymal element can be either homologous (gynecologic tissue) or heterologous (non-gynecologic tissue) sarcoma^2,3^. At the histology level, the diverse compositions of these elements contribute significantly to the tumor heterogeneity within UCS, which has impacts on its poor prognosis. In terms of outcome, the clinical course of UCS is more like that of its carcinoma counterpart presenting with the same histology rather than that of sarcoma counterpart in terms of risk factor profile, dissemination pattern, and sensitivity to cytotoxic agents^4,5^. Therefore, the understanding of tumor heterogeneity within UCS will be crucial to help identify pathophysiological mechanisms leading to novel therapeutic interventions.

Several mechanisms and hypotheses have been proposed to explain the biphasic histologies of UCS^2-4^. The generation of the dual histological pattern has been supported by the conversion theory with many studies proposing epithelial-mesenchymal transition (EMT) as a potential mechanism for converting carcinoma cells into sarcoma cells^4^. This conversion theory is confirmed by genomic profiling and phylogenetic evolution analysis of the two components of a large UCS cohort to validate the histogenic hypothesis with a clonal origin with retention of driver events^4^. Quantification of EMT scores of the two histological components has further indicated that the sarcoma component showed a higher mesenchymal score compared to the carcinoma component^4^. These changes of EMT expression patterns are also correlated with hypermethylation of the CpG regions of miR-141/200a/200b/200c/429^4^, the known EMT suppressors to antagonize the EMT inducing transcription factor, ZEB1. These findings put the metaplastic differentiation of clonal origins onto the center stage of contention for tumor heterogeneity of UCS and further suggest that cellular plasticity contributes to the complexity of UCS.

In addition to the heterogeneity and complexity of cancer cells in UCS, the tumor microenvironment (TME) associated with these diverse cancer cells also play important roles in shaping the UCS landscape. EMT is known to link to immunosuppression in the TME with the mesenchymal states being associated with lower expression of MHC-1 in the cancer cells, increased infiltration of CD4+Foxp3+ regulatory T cells, M2 macrophages, and exhausted CD8+ T cells, and elevated immune checkpoint molecules, such as PD-L1 and PD-L2 in the cancer cells, and PD-1 and CTLA-4 in tumor infiltrated T cells^6-8^. It is therefore very likely that there also exists distinct TME features within the dual histological components of UCS. Several studies^9-12^ have investigated the immune TME of UCS but with highly variable and even contradictory results. Karpathiou et al. showed that CD3 expression was more frequent in the sarcoma than in the carcinoma component in 37 patients with gynecological carcinosarcoma, including 22 UCS^9^. This seemed to contradict the findings from Da Silva et al. that only 10.5% of patients showed highly positive CD3 expression in sarcoma component^12^. This discrepancy was also evident from studies investigating PD-1 and PD-L1 expressions. Gulec et al. surveyed 59 UCS and found that both PD-1 and PD-L1 were detected in a quarter of cases^10^. In 75 carcinosarcoma cases, including 60 UCS, Hacking et al. reported that 65 (87%) patients demonstrated positive PD-1 expression in tumor cells and 50 (67%) patients had positive PD-L1 expression in immune stromal components^11^. Da Silva et al. showed that PD-1 and PD-L1 were highly expressed in the carcinoma component in about half of patients and in the sarcoma component in about a quarter of patients. However, PD-L1 was negative in all cases studied in Karpathiou et al.^9^. The variability in marker expression could be attributed to the limitation of the method used in these studies which was immunohistochemistry (IHC) staining. Although IHC can detect the presence of specific immune markers, it is difficult to quantify the expressions levels, and the definition of positive expression varies across different studies. Therefore, it is necessary to adopt other methods to assess the heterogeneity within the UCS TME.

Next generation sequencing-based methods such as CIBERSORT, a versatile computational method for quantifying cell fractions from bulk tissue gene expression profiles, have been shown to facilitate the deconvolution of components within TME^13^. By using this method, the immunogenomic landscape of gynecologic carcinosarcoma has been established^14^. CIBERSORT scores from the RNA-sequencing results of carcinoma and sarcoma components separately collected by laser capture microdissection (LCM) revealed a distinct pattern of multiple cell-type infiltration scores between the two components. Plasma cells, M2 macrophages, endothelial cells and fibroblasts were significantly enriched in the sarcoma component, and a high infiltration of T cells and high cytolytic activity scores were also found in the sarcoma component. On the other hand, M0 macrophages and follicular helper T cells were more frequently found in the carcinoma component^14^.

With the recent advancement of spatial profiling technologies, there has been a significant leap of data quality and quantity to enable teasing out the tumor heterogeneity within TME. The technology solution such as digital spatial profiling (DSP), a region of interest (ROI)-based method which can perform quantitative, high-plex analysis of transcripts or proteins simultaneously in spatially defined regions, has been adopted to decipher the TME architecture in various cancer types^15^. The non-destructive nature of spatial profiling has tremendously solved the limitations faced by methods such as LCM that the profiling could be done in tumor regions expressing specific markers or even down to single cell resolutions. In this study, we used DSP to investigate the inter– and intratumor heterogeneity in immune TME in spatially defined lesions of UCS. We identified specific features of infiltrating immune signals within the carcinoma and sarcoma components that were clinically significant in patient’s outcomes.

## Materials and methods

### Case recruitment

Patients diagnosed with uterine carcinosarcoma (UCS) between November 1, 2010, and December 31, 2020, from the Cancer Registry of National Taiwan University Hospital (NTUH) were enrolled for this study. Inclusion criteria comprised: (1) documented pathology reports confirming UCS diagnosis; (2) identifiable discrete biphasic histology phenotypes of carcinoma and sarcoma components via hematoxylin and eosin (H&E) staining (verified by experienced pathologists WC Lin); (3) absence of chemotherapy, immunotherapy, or radiotherapy prior to cancer staging operations; (4) absence of immunocompromised conditions. Surgical / pathologic staging follows definitions from the AJCC Cancer Staging Manual (8th edition) and the FIGO 2018 update. Overall survival (OS) is defined as the interval between staging surgery and death. Progression-free survival (PFS) is defined as the interval between staging surgery and the first documented disease progression. Institutional review board approval was obtained from the ethics committee of National Taiwan University Hospital (NTUH) (202005080RIND). Clinical details are summarized in Table 1.

**Table 1.** Twenty-three patients’ clinical data.

| **Variables (n=23)** |  |
| --- | --- |
| Age (mean +/-SD, years) | 60.7 +/-8.0 |
| BMI (mean +/-SD) | 24.1+/-3.1 |
| Parity (mean +/-SD) | 2.2 +/-1.5 |
| menopause, n (%) | 20 (87) |
| FIGO stage, n (%) |  |
| I | 8 (34.7) |
| IA | 5 (21.7) |
| IB | 3 (13.0) |
| II | 1 (4.3) |
| III | 10 (43.5) |
| IIIA | 2 (8.7) |
| IIIB | 0 (0) |
| IIIC | 8 (34.8) |
| IV | 4 (17.3) |
| Lymphovascular invasion positive, n (%) | 18 (78.3) |
| Ascites cytology positive, n (%) | 6 (26.1) |
| Carcinoma type, n (%) |  |
| endometrioid | 8 (34.8) |
| Serous | 4 (17.4) |
| clear | 1 (4.3) |
| Endometrioid & serous | 2 (8.7) |
| Endometrioid & mucinous | 1 (4.3) |
| High grade | 3 (13.0) |
| Non-specific | 4 (17.4) |
| Sarcoma type, n (%) |  |
| homologous | 1 (4.3) |
| heterologous | 9 (39.1) |
| Non-specific | 13 (56.5) |
| Adjuvant therapy, n (%) |  |
| Chemotherapy + radiotherapy | 6 (26.1) |
| Chemotherapy only | 11 (47.8) |
| Radiotherapy only | 4 (17.4) |
| No adjuvant therapy | 2 (8.7) |
| Outcomes |  |
| Progression free median survival (PFS), month (range) | 17.1 (4.8-120.6) |
| Overall median survival (OS), month (range) | 29.5 (4.8-120.6) |
| 5-year PFS, n (%) | 7 (30.4) |
| 5-year OS, n (%) | 9 (39.1) |

### Sample processing

After verifying the slides, we retrieved the formalin-fixed paraffin-embedded (FFPE) sections (5μm) from the Department of Pathology of NTUH. FFPE tissue sections were baked at 60°C for 30 minutes. Deparaffinization was performed using CitriSolv, followed by sequential rehydrated in 100% and 95% ethanol and a wash in ddH_2_O. Antigen retrieval was achieved by boiling at 121°C in pH 6 Citrate buffer solution for 15 minutes in a pressure cooker. Tissue sections were then blocked in a blocking buffer for 1 hour at room temperature before being incubated overnight at 4°C in the dark with a panel of antibodies conjugated with UV photocleavable oligonucleotide barcodes for nanoString GeoMx DSP comprising of Immune Cell Profiling, Immune Activation Status, Immune Cell Typing, PanCK-Cy3 (1:40) and CD45-Texas Red (1:40). Sections were stained with SYTO13 (1:10) for 15 minutes the following day after being post-fixed in 4% paraformaldehyde (PFA).

### ROI selection and collection

Regions of interest (ROIs) were identified based on the morphology of H&E staining to distinguish the carcinoma, sarcoma, and stroma components of each sample. Areas of illumination (AOIs) were delineated separately for the carcinoma, sarcoma, and stroma components. Carcinoma and sarcoma components were further defined and gated based on their tumor periphery or tumor center locations to collect geometric segments. The “central area” refers to the region distant from the boundary of other histology parts, whereas the “peripheral area” denotes the region near other histology parts. Subsequently, signal intensity from these segments (central carcinoma, peripheral carcinoma, central sarcoma, peripheral sarcoma, and stroma) was gated to select three AOIs with each segment. Additionally, three extra ROIs were selected in each component to collect AOIs based on the presence or absence of CD45 signals within the segments (Figure 1A)^16^. The UV-cleaved oligonucleotides were collected and dispensed into a microtiter plate. Afterward, the tags were hybridized, pooled and quantified using the nCounter MAX/FLEX system.

**Figure 1.** (A) Workflow of the identification and selection of ROIs and AOIs. (B) Distribution chart showing numbers of selected ROIs in each component. (C) The clustering heatmap of AOI. ROI: region of interest; AOI: area of illumination

Twenty-three FFPE slides of primary UCS from 23 patients were enrolled in this study. Ultimately, 491 AOIs from different segments at different histological components (carcinoma or sarcoma) of primary UCS were included in the subsequent analysis. Among these, 257 AOIs were from geometric segments: 63 AOIs at central sites of carcinoma components, 63 AOIs at peripheral sites of carcinoma components, 3 AOIs at carcinoma components where the center or periphery could not be clarified, 58 AOIs at central sites of sarcoma components, 57 AOIs at peripheral sites of sarcoma components, 6 AOIs at sarcoma components where the center or periphery could not be clarified, and 25 AOIs at stroma components. Additionally, 106 AOIs were from CD45-positive segments with 41 AOIs at carcinoma components, 36 AOIs at sarcoma components and 29 AOIs at stroma components. 110 AOIs were from CD45-negative segments with 42 AOIs at carcinoma components, 39 AOIs at sarcoma components, and 29 AOIs at stroma components (Figure 1B).

### Data analysis

The readouts from nCounter (version 4.0.0.3) were transferred to GeoMx, and quality control and normalization were performed using the built-in data analysis software (version 2.1.0.33). Initially, AOIs were flagged if they had a positive normalization number >3, a minimum nuclei count < 20, or a minimum surface area < 1600μm^3^. AOIs were excluded if the positive normalization number exceeded 20. Then, the data were then normalized using housekeeper normalization. Following normalization, the data were log-transformed, mean-centered, and subjected to unsupervised hierarchical clustering using a similarity metric of Pearson correlation and a tree-construction method of centroid linkage.

The normalized data were extracted and utilized to create volcano plots, box plots and violin plots. Statistical significance of association was assessed using the Chi-square test, while median differences were evaluated using the Mann-Whitney test. Differences in PFS were assessed using Kaplan-Meier analysis.

## Results

### Clinical characteristics of the UCS cohort

As shown in Table 1, the mean age at UCS diagnosis of this cohort was 60.7 years old. 39.1% of cases were early-stage disease, and 60.9% were late-stage disease. Lymphovascular space invasion (LVSI) was seen in most cases (78.3%). The rate of malignant peritoneal cytology was 26.1% which is higher than the previous study^2,17,18^. The median PFS time was 17.1 months, and the 5-year PFS rate was 29.5%. The median OS time was 29.5 months, and the 5-year OS rate was 39.1%. The clinical demographics of this present cohort was comparable with a previous Japanese study which included 483 UCS patients^19^.

### Unsupervised hierarchical clustering of areas of interest (AOIs) showing heterogeneity of immune marker expression

A total of 518 AOIs were collected, with twenty-seven outliers excluded after quality check, resulting in 491 AOIs kept for the subsequent analysis. Unsupervised hierarchical clustering based on the correlation of the composition of quantified immune-related cell markers showed that the AOIs were classified into five groups (Figure 1C). The C1 cluster exhibited a hot immune feature with several immune-related cell markers, such as PD-L1, PD-L2, CTLA4, CD80, CD4, CD25, FOXP3, CD163, ICOS, CD8, and beta-2-microglobulin (B2M), were relatively high (Figure 2A-K). CD56 expression was relatively low in C1 (Figure 2L). The C2 cluster also exhibited hot immune features as well as the immune-related cell markers observed in C1 and also showed high expression of smooth muscle action (SMA), fibronectin (FN), and CD56 (Figure 2L-N). The C3 cluster showed a generally cold immune feature. The C4 cluster was subdivided into two subgroups C4a and C4b, with the C4a subgroup showing a relatively high expression of CD66b (Figure 2O), while the C4b subgroup showed relatively high expression of CD56 and Ki-67 (Figure 2L, Figure 2P). These clusters indicated the diverse features within UCS tumors which could inform the landscape of immune heterogeneity.

**Figure 2.** (A) A violin plot showing the level of PD-L1 at each cluster. (B) A violin plot for showing the level of PD-L2 at each cluster. (C) A violin plot showing the level of CTLA4 at each cluster. (D) A violin plot showing the level of CD80 at each cluster. (E) A violin plot showing the level of CD4 at each cluster. (F) A violin plot showing the level of CD25 at each cluster. (G) A violin plot showing the level of FOXP3 at each cluster. (H) A violin plot showing the level of CD163 at each cluster. (I) A violin plot showing the level of ICOS at each cluster. (J) A violin plot showing the level of CD8 at each cluster. (K) A violin plot showing the level of beta2-microglobulin at each cluster. (L) violin plot showing the level of CD56 at each cluster. (M) A violin plot showing the level of fibronectin at each cluster. (N) A violin plot showing the level of SMA at each cluster. (O) A violin plot showing the level of CD66b at each cluster. (P) A violin plot showing the level of Ki-67 at each cluster. (Q) A stacked plot of the proportion of immune expression of sarcoma, stratified by C4a carcinoma. (R) A stacked plot of the proportion of immune expression of sarcoma, stratified by C4b carcinoma. **: p-value =0.001∼0.01; ***: p-value =0.0001∼0.001; ****: p-value <0.0001

### Correlation of immune cluster labeling between carcinoma and sarcoma

Distributions of these clusters were further correlated between carcinoma and sarcoma components. According to the conversion theory, the sarcoma components were believed to originate from the carcinoma components. We thus compared whether the AOIs from the carcinoma and sarcoma components within the same patient sample would exhibit concordant or disparate cluster labeling. The predominant cluster labeling of the carcinoma component within the same patient sample was used as the reference. The labeling of the sarcoma component was thus annotated as being the same or different cluster. AOIs within carcinoma components exhibited cluster labeled as C4a (45.1%), followed by C4b (43.1%). C3 (7.8%), C2 (2.9%), and C1 (1.0%) (Supplementary Figure 1). Carcinoma AOIs with the cluster label of C4a or C4b were used in the stratification for analysis. Among patient samples with AOIs predominantly annotated as C4a within the carcinoma component, 85.7% exhibited different cluster labeling in the sarcoma counterpart. This frequency was significantly higher (p= 0.025, Chi-square test) compared to those annotated as non-C4a (Figure 2Q). However, this trend was not observed in patient samples with carcinoma AOIs predominantly annotated as C4b, that approximately 61.9% and 74.1% of C4b and non-C4b annotated carcinoma AOIs showed different cluster labeling in the sarcoma counterpart (Figure 2R). Nevertheless, the majority of the AOIs showed discordant cluster labeling between the carcinoma and sarcoma components in UCS. These cluster labeling results suggested that the immune features of the sarcoma components of UCS might further undergo remodeling with the sarcoma counterpart of the carcinoma C4a cluster being the least concordant. Since the AOIs subjected to clustering consisted of non-segmented geometric and segmented AOIs (Figure 1B), we continued to stratify the analysis based on the nature of these AOIs to further explore the heterogeneity.

### Intra-tumor immune heterogeneity analysis within geometric AOIs

Geometric AOIs could be seen as the mini-bulk consisting of both the tumor cell and the tumor microenvironment. Therefore, the assessment of the immune cluster labeling of the geometric AOIs could provide comprehensive characterization of the neighborhood at different histologic components. As shown in Figure 3A, the immune clusters were diverse in both histologic components. Within the carcinoma component, the majority of AOIs were classified as C4b (44.2%), followed by C4a (28.7%) and C3 (20.9%), with fewer AOIs classified as C2 (3.1%) or C1 (3.1%). In contrast, the sarcoma component was primarily composed of C4b (48.8%), followed by C2 (28.1%), with smaller proportions classified as C4a (12.4%) or C3 (10.7%) without any presence of C1. This indicated a distinct distribution pattern of the two histologic components within each cluster (Figure 3B), that with cluster labeling of C3 and C4a, the proportion of carcinoma was higher than that of sarcoma (p=0.028 and p=0.002, respectively), whereas the proportion of sarcoma was higher within C2 (p<0.0001). Notably, only carcinoma geometric AOIs of carcinoma exhibited C1. The sarcoma geometric AOIs showed significantly higher expression of almost all immune markers compared to the carcinoma AOIs (Figure 3C). The expressions of SMA and Fibronectin were significantly higher in sarcoma (SMA: p <0.05, FC=-2.44; Fibronectin: p <0.05, FC=-2.11), indicating the mesenchymal nature. There was no significant difference of the immune cluster labeling frequency between geometric AOIs located at the central or the peripheral parts of the carcinoma (Figure 3D) or the sarcoma (Figure 3E) components.

**Figure 3.** Geometric AOIs analysis (A) A stacked bar plot of the immune composition within each histology component. (B) A stacked bar plot of the proportion of histology within each cluster. (C) A volcano plot showing the difference in immune-related marker levels between the sarcoma and the carcinoma components. (D) A stacked plot of immune composition at the central and peripheral sites of the carcinoma component. (E) A stacked plot of immune composition at the central and peripheral sites of the sarcoma component. *: p-value=0.01∼0.05; **: p-value=0.001∼0.01; ****: p-value<0.0001

The expression of CTLA4 in the carcinoma component geometric AOIs was significantly higher at the central site than at the peripheral site (p=0.02) with a fold change of 1.45 (Supplementary Figure 2A). Conversely, the expression level of all immune markers in the sarcoma geometric AOIs showed no significant difference at both sites (Supplementary Figure 2B).

### Intra-tumor immune heterogeneity analysis within CD45+ AOIs

To further tease out the intra-tumor heterogeneity of tumor-infiltrated lymphocytes composition, CD45+ AOIs were subjected to the cluster distribution analysis, revealing heterogeneous presentations within and between histological components. In the carcinoma component, most of the CD45+ AOIs were classified as C4a (63.4%), followed by C4b (26.8%), with a few C2 (9.8%). Conversely, the sarcoma component was primarily annotated as C2 (41.7%) and C4a (50.0%), with only a small number of AOIs classified as C1 (5.5%) or C4b (2.6%) (Figure 4A). Distinct patterns of distribution of the two histologic components within each cluster were also observed. C1 CD45+ AOIs were purely located at the sarcoma component. The majority of C2 CD45+ AOIs (78.9%) were preferentially located at the sarcoma component (p=0.001). On the contrary, the majority of C4b CD45+ AOIs (91.7%) were located at the carcinoma component rather than at the sarcoma component (p=0.004) (Figure 4B). The distribution of C4a CD45+ AOIs at carcinoma and sarcoma components was approximately equal. This indicated that the immune infiltration within the carcinoma and sarcoma components of UCS were distinctly different. To further characterize the identify of these immune infiltrates, differential expression of the immune markers between the two histology components was analyzed. Similar to the results found in the geometric AOIs (Figure 3C), most of the immune markers in the sarcoma component exhibited significantly higher levels than in the carcinoma component (Figure 4C). The most significantly expressed immune markers included CD163, with a fold change (FC) of about 2.88 (p <0.0001), PD-L2, with a fold change of about 2.3 (p=0.0006), SMA and fibronectin with fold changes of 2.06 (p=0.009) and 1.61 (p=0.01), respectively. The data suggested that the immune infiltrates inside the sarcoma components of UCS were potentially enriched for CD163+ macrophages with high expression of myofibroblast markers.

**Figure 4.** CD45 AOIs analysis (A) A stacked bar plot of the CD45+ immune composition within each histology component. (B) A stacked bar plot of the proportion of histology component within each cluster in CD45+ AOI analysis. (C) A volcano plot about the differences in immune-related cell marker levels between the sarcoma and the carcinoma components in the CD45+ AOI analysis. (D) A stacked bar plot of the CD45-immune composition within each histology component. (E) A stacked bar plot of the proportion of histology components within each cluster in the CD45-AOI analysis. (F) A volcano plot showing the differences in immune-related marker levels between the sarcoma and carcinoma components in the CD45-AOI analysis. **: p-value=0.001∼0.01; ****: p-value<0.0001

### Intra-tumor immune heterogeneity analysis within CD45-AOIs

To assess the tumor microenvironment components other than immune cells, CD45-AOIs were subjected to the cluster distribution analysis. Diverse presentations of the clusters in each histological component was observed (Figure 4D). In the carcinoma component, the majority of AOIs were classified as C4a (52.4%), followed by C4b (45.2%), with a small proportion classified as C2 (2.4%). Conversely, most of the AOIs in the sarcoma component were classified as C2 (46.2%), followed by C4b (41.0%) and C4a (10.2%), with only a few AOIs classified as C1 (2.6%). Distinct patterns of distribution of the two histologic components within each cluster were observed again. C1 CD45-AOIs were exclusively located in the sarcoma component. However, the frequency of C4a CD45-AOIs was higher in the carcinoma than in the sarcoma components (p<0.0001), while the frequency of C2 CD45-AOIs was higher in the sarcoma than in the carcinoma components (p<0.0001) (Figure 4E). The frequencies of C4b CD45-AOIs in carcinoma and sarcoma components were approximately equal. This indicated that the microenvironment within the carcinoma and sarcoma components of UCS were also distinctly different. Differentially expressed markers revealed that SMA (p<0.05, FC=-4.2), fibronectin (p<0.05, FC=-2.86), CD34 (p<0.05, FC=-2.49) and CD56 (p<0.05, FC=-2.46) were significantly up-regulated in the sarcoma component (Figure 4F). CD66b seemed to be more highly expressed in the carcinoma component though without reaching statistical significance. The data suggested that the microenvironment within the sarcoma component of UCS were enriched for a pro-fibrotic or pro-angiogenic milieu.

### Inter– and intra-tumor immune heterogeneity associated with UCS patient outcomes

There was a tremendous degree of inter– and intra-tumor heterogeneity in terms of cluster labeling of the UCS patient cohort (Supplementary Figure 2C, Supplementary Figure 2D, Supplementary Figure 2E). Among the patients studied, seven experienced no disease recurrence or progression over a period of 5 years, while the remaining sixteen patients experienced recurrent disease within 5 years. We next analyzed whether there would be distribution patterns of these cluster labels associated with UCS patient outcomes based on the stratification of geometric, CD45+, CD45-AOIs.

In the analysis using geometric AOIs, a notable difference was observed in the cluster labeling of the carcinoma component between patients who remained free of disease progression for over 5 years and those with recurrent disease. Specifically, the frequency of C4a was prominent in the carcinoma component of patients who remained free of disease progression over 5 years (53.8%), compared to those with recurrent disease (17.8%). Conversely, the cluster labeling of the carcinoma component of patients with recurrent disease was mainly composed of C4b (54.4%), whereas the proportion of C4b in the carcinoma component of patients without disease progression over 5 years was only 20.5% (Figure 5A). The progress-free survival (PFS) was poorer in patients with tumors exhibiting non-C4a in the carcinoma component (p<0.0001, Figure 5B) and tumors displaying C4b cluster labeling in the carcinoma component (p<0.0001, Figure 5C). A z-score was employed to investigate whether tumors from patients with long PFS were more likely to exhibit the representative immune cell markers of C4a at the carcinoma component. The expression level of CD66b was significantly higher at the carcinoma component of tumors from patients with long PFS compared to those with short PFS (p<0.0001) (Figure 5D). Conversely, the levels of CD56 and Ki-67 were significantly higher at the carcinoma component of tumors from patients with short PFS compared to those with long PFS (p<0.0001, p<0.0001) (Figure 5E-F).

**Figure 5.** (A) A stacked plot of immune composition within different histology components and different disease statuses within geometric AOIs analysis. (B) The Kaplan-Meier survival curve of tumors with carcinoma showing C4a immune expression versus the rest in geometric AOIs analysis. (C) The Kaplan-Meier survival curve of tumors with carcinoma showing C4b immune expression versus the rest in geometric AOIs analysis. (D) Box plot showing the z-score distribution of CD66b in carcinoma component with disease with and without recurrence over 5 years in geometric AOIs analysis. (E) Box plot showing the z-score distribution of CD56 in carcinoma component with the disease with and without recurrence over 5 years in geometric AOIs analysis. (F) Box plot showing the z-score distribution of Ki-67 in carcinoma component with disease with and without recurrence over 5 years in geometric AOIs analysis. (G) A stacked plot of immune composition within different histology components and different disease status in CD45+ AOIs analysis. (H) The Kaplan-Meier survival curve of tumors with carcinoma showing C4a immune expression versus the rest in CD45+ AOIs analysis. (I) The Kaplan-Meier survival curve of tumors with sarcoma showing C2 immune expression versus the rest in CD45+ AOIs analysis. (J) The Kaplan-Meier survival curve of tumors with sarcoma showing C4a immune expression versus the rest in CD45+ AOIs analysis. (K) Box plot showing the z-score distribution of CD66b in carcinoma component with disease with and without recurrence over 5 years in CD45+ AOIs analysis. (L) Box plot showing the z-score distribution of SMA in sarcoma component with disease with and without recurrence over 5 years in CD45+ AOIs analysis. (M) Box plot showing the z-score distribution of fibronectin in sarcoma component with disease with and without recurrence over 5 years in CD45+ AOIs analysis. (N) A stacked plot of immune composition in different histology components and different disease status in CD45-AOIs analysis. (O) The Kaplan-Meier survival curve of tumors with carcinoma showing C4a immune expression versus the rest in CD45-AOIs analysis. (P) The Kaplan-Meier survival curve of tumors with carcinoma showing C4b immune expression versus the rest in CD45-AOIs analysis. (Q) Box plot showing the z-score distribution of CD66b in carcinoma component with disease with and without recurrence over 5 years in CD45-AOIs analysis. (R) Box plot showing the z-score distribution of CD56 in carcinoma component with disease with and without recurrence over 5 years in CD45-AOIs analysis.

Furthermore, CD45+ AOIs analysis was utilized to assess tumor-infiltrated immune cells in different tumors with varying prognoses. It was observed that all CD45+AOIs in the carcinoma of tumors that remained free of recurrence over 5 years were C4a (Figure 5G). The PFS was significantly longer in tumors where CD45+ AOIs in the carcinoma were C4a compared to those with non-C4a (p=0.05, Figure 5H). This finding corroborated with the result obtained from the analysis of geometric AOIs. Similarly, the majority of the CD45+ AOIs at the sarcoma of tumors in patients that remained free of recurrence over 5 year were C2 (75%), and these C2 CD45+ AOIs in the sarcoma component were associated with disease outcomes (Figure 5I). Although the association did not reach statistical significance (p=0.054), there was a trend suggesting that patients with tumors having C2 CD45+ AOIs in the sarcoma component tended to have longer PFS compared to others with non-C2 CD45+ AOIs (Figure 5I). 57.1% of the CD45+ AOIs at the sarcoma component of tumors from patients with short PFS were C4a, whereas only 25.0% of the CD45+ AOIs at the sarcoma component of tumors from patients with long PFS were C4a (Figure 5G). However, the difference in PFS did not reach statistical significance (p=0.21) (Figure 5J). A z-score was again employed to investigate whether tumors from patients with long PFS were more likely to present representative immune markers of C4a at the carcinoma component (Figure 5K). The level of CD66b was found to be significantly higher at the carcinoma component of tumors from patients with long PFS compared to those with short PFS (p=0.0012, Figure 5K). Similarly, the same score was utilized to determine whether tumors from patients with long PFS were more likely to present representative immune markers of C2 at the sarcoma component. However, the levels of SMA and fibronectin did not show significant differences between tumors with and without recurrence. (Figure 5L-M)

The CD45-AOIs analysis was conducted to assess the tumor immune microenvironment components other than lymphocytes in different tumors with varying prognoses. It was found that all the CD45-AOIs at the carcinoma of tumors from patients that remained free of recurrence over 5 years were C4a (Figure 5N). The survival was significantly longer in tumor where CD45-AOIs in the carcinoma were C4a compared to those with non-C4a (p<0.0001, Figure 5O). Conversely, a majority of the AOIs at the carcinoma of tumors from patients with short PFS were C4b (63.3%, Figure 5N), and the survival was significantly shorted when the CD45-AOIs at the sarcoma were C4b compared to those with non-C4b (p<0.0001, Figure 5P). In tumors from patients with long PFS, 60% of AOIs at the sarcoma component were C2 and 40% were C4b (Figure 5N). For tumors from patients with short PFS, 41.4% of CD45-AOIs at the sarcoma component were C2, and likewise, 41.4% of CD45-AOIs were C4b (Figure 5N). Additionally, 13.8% of CD45-AOIs at the sarcoma component of tumors from patients with short PFS were C4a, and 3.4% were C1 (Figure 5N). The CD45-AOIs composition at the sarcoma component was similar between tumors from patients with long and short PFS. A z-score analysis was employed to investigate whether a tumor from patients with long PFS would be more likely to present the representative immune markers of C4a at the carcinoma component. The level of CD66b was significantly higher at the carcinoma component of tumors from patients with long PFS compared to those with short PFS (p=0.005, Figure 5Q). Similarly, the z-score was also utilized to determine whether a tumor from patients with short PFS would be more likely to present the representative immune markers of C4b at the carcinoma component. The level of CD56 was significantly higher at the carcinoma component of tumors from patients with short PFS (p=0.009, Figure 5R).

## Discussion

There has been a paradigm shift of the standard of care in endometrial cancer with immune-oncology (IO) agents being proven as an effective agent to improve treatment outcomes. The elucidation of the complexity of the uterine microenvironment is crucial to further understand why patients with certain tumor characteristics fail to respond.

Generally, innate and adaptive immune cells with PD-1 ad PD-L1 expression can be found in the endometrial cancer microenvironment^20,21^. The presence or absence of these infiltrated immune cells within the uterine microenvironment could further characterize each uterine cancer into heterogeneous immune subgroups. For instance, despite that uterine leiomyosarcoma is an immune cold tumor with only a few tumor infiltrating lymphocytes (TILs)-identified in H&E sections, a small subset of leiomyosarcoma with immune hot characteristics could be identified by RNA sequencing^22,23^. Technologies enabling spatial profiling could indeed advance the identification of many more heterogeneous immune subtypes of endometrial cancer.

With the help of Digital Spatial Profiling (DSP) technology, we are able to indicate that the immune presentations with different histology components of uterine carcinosarcoma (UCS) are different. In previous UCS studies^12,14^, da Silva et al. reported that T-cell markers, such as CD3, CD4, CD8, FOXP3, PD-L1 and PD-L2, were highly expressed in most of the carcinoma components, but the frequencies of highly positive expression were lower in the sarcoma components^12^. Gotoh et al. analyzed seven UCS cases and identified a distinct pattern of infiltrated immune cell types between the two components^14^. In our cohort, the sarcoma component had high infiltration of hematologic-origin CD45+CD163+ macrophages, expressed various immune-related cell markers with EMT-related markers fibronectin and SMA being especially enriched. On the contrary, the carcinoma component had a relatively cold immune presentation but showed a relatively higher level of CD66b+ cells. The evidence clearly suggests that the carcinoma and the sarcoma components within UCS tumors would provide diversified microenvironment milieus that could potentially dictate disease outcomes.

Matsuo et al. found that the sarcoma component of UCS was more likely to spread locally^24^. This could be explained by our findings that the sarcoma component exhibited high levels of EMT-related markers fibronectin and SMA. Fibronectin is one of the extracellular matrix (ECM) proteins and interacts with many other ECM proteins to provide signals to induce specific cell behaviors such as differentiation and EMT^25^. SMA contributes to maintaining cell shape and is important for tumor cell migration and invasion, and its upregulated expression is associated with EMT^26^. In addition, the increased presence of CD45+CD163+ cells within the sarcoma component indicated the infiltration of M2 macrophage. The specialized tumor-associated macrophages (TAMs) play a significant immunosuppressive role and are related to tumor progression and metastasis via promoting angiogenesis, immunosuppression, and activation of the tumor cells^27,28^. These TAMs together with various of immune-suppressive cells have also been found to induce EMT through TGFβ signaling. These tissue-resident immune cells with hematopoietic origin contribute significantly to the shaping of the sarcoma TME within UCS.

Intriguingly, this CD45+CD163+ M2 macrophage predominant sarcoma TME (cluster label C2) did not seem to determine the UCS clinical outcomes in our cohort. Our data showed no strong significance in immune presentations within the sarcoma components between the tumors with recurrence within 5 years and those without recurrence for over 5 years. This is in accordance with the known fact that the clinical prognosis of UCS is mainly determined by the carcinoma component^2^.

The high-grade nature of the carcinoma component acts as a driving factor for disease progression and metastasis^2^. Indeed, our data showed significantly different immune presentations in the carcinoma components between the tumors with recurrent within 5 years and those without recurrence for over 5 years. The immune presentations in the carcinoma components of tumors without recurrence showed a higher tendency to express CD66b than those with recurrence. The significance of the impact to UCS outcome was demonstrated in all survival analysis from the geometric segments, CD45+ and CD45-AOIs that had cluster label of C4a with high CD66b expression.

The biomarker CD66b can be found on tumor-associated neutrophils (TAN) and polymorphonuclear myeloid-derived suppressor cells (PMN-MDSC), and the biomarker is related to prognosis and is a predictive marker for treatment ^29^. CD66b expression has been shown to have divergent impacts on cancer prognosis. In early-stage lung^30,31^ and oral cancer^32^, CD66b expression was associated with good prognosis. In advanced-stage lung and oral cancers and gastric cancer^33^, CD66b was found to be associated with short PFS and OS and was related to the resistance to immune checkpoint inhibitors in lung cancer. Our data indicated that CD66b expression in the carcinoma component is associated with the favorable prognosis of UCS. This suggests that the TME shaped by CD66b expressing cells in different contexts is much more complex than dichotomy.

Since the biomarkers expressed by PMN-MDSCs mostly overlap with those that were used to identify neutrophils, and PMN-MDSCs often coexisted with neutrophils, it is difficult to precisely pinpoint whether it was PMN-MDSC or TAN playing a role in UCS. Nevertheless, both PMN-MDSC and TAN are known to have effects in immune suppression within the TME. Neutrophils can form extracellular traps, but these traps may trap circulating tumor cells and prevent interaction between cytotoxic immune cells and tumor cells^29,34^. MDSCs inhibit the activities of activated immune cells and suppress the T-cell-mediated antitumor immune response. They restrain T-cell proliferation, recruitment, and induce T-cell apoptosis^35^. PMN-MDSCs induce immunosuppression and promote cell motility and proliferation of dormant tumor cells. They drive the differentiation of CD4+ T cells into regulatory T cells, inhibits T cell proliferation and responsiveness to antigen-specific stimulation, and prevents the infiltration of T cells into the central tumor^29,34^. Thus, why carcinoma components expressing high CD66b would associate with favorable prognosis in UCS is yet to be studied.

## Conclusions

Our data reveal the presence of both intra– and inter-tumor heterogeneous immune presentations in UCS. The levels of immune cell markers were generally higher in sarcoma components than in carcinoma components, with EMT-related markers being significantly higher. Additionally, the sarcoma component exhibited a higher level of CD45+CD163+ immune-related cell marker than the carcinoma component. Conversely, the carcinoma component displayed a relative cold immune presentation but showed a relatively higher level of CD66b+ TME. Furthermore, these heterogeneous immune presentations were associated with different clinical outcomes. UCS with a higher CD66b level in carcinoma components was related to better clinical outcomes than those with lower CD66b level. The underlying mechanisms by which carcinoma components with high CD66b expression correlate with favorable prognosis in UCS remain to be explored.

## Limitations

Our study is limited by the small sample size due to the nature of a rare cancer. The spatial profiling technology used was not at the single cell resolution so that each “mini bulk” of the segments could further consist of more heterogeneity. The CD45+ segments could be further analyzed by single cell spatial transcriptomics to further distinguish the immune cell types within the carcinoma and sarcoma components. The expression of certain immune-related markers did not indicate activity and functional engagement. Therefore, further studies are needed to clarify the definitive roles of CD66b expressing cells.

---

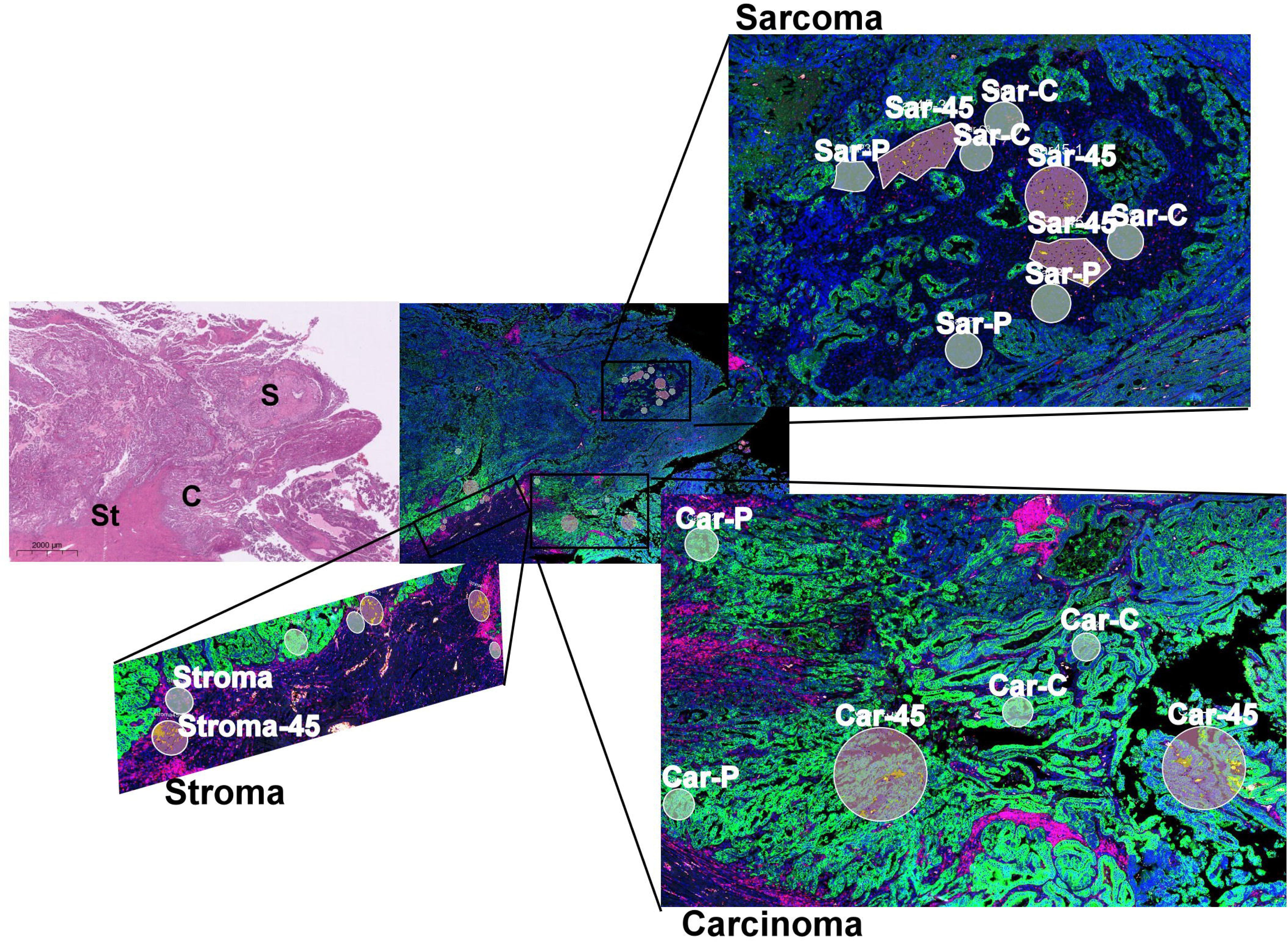

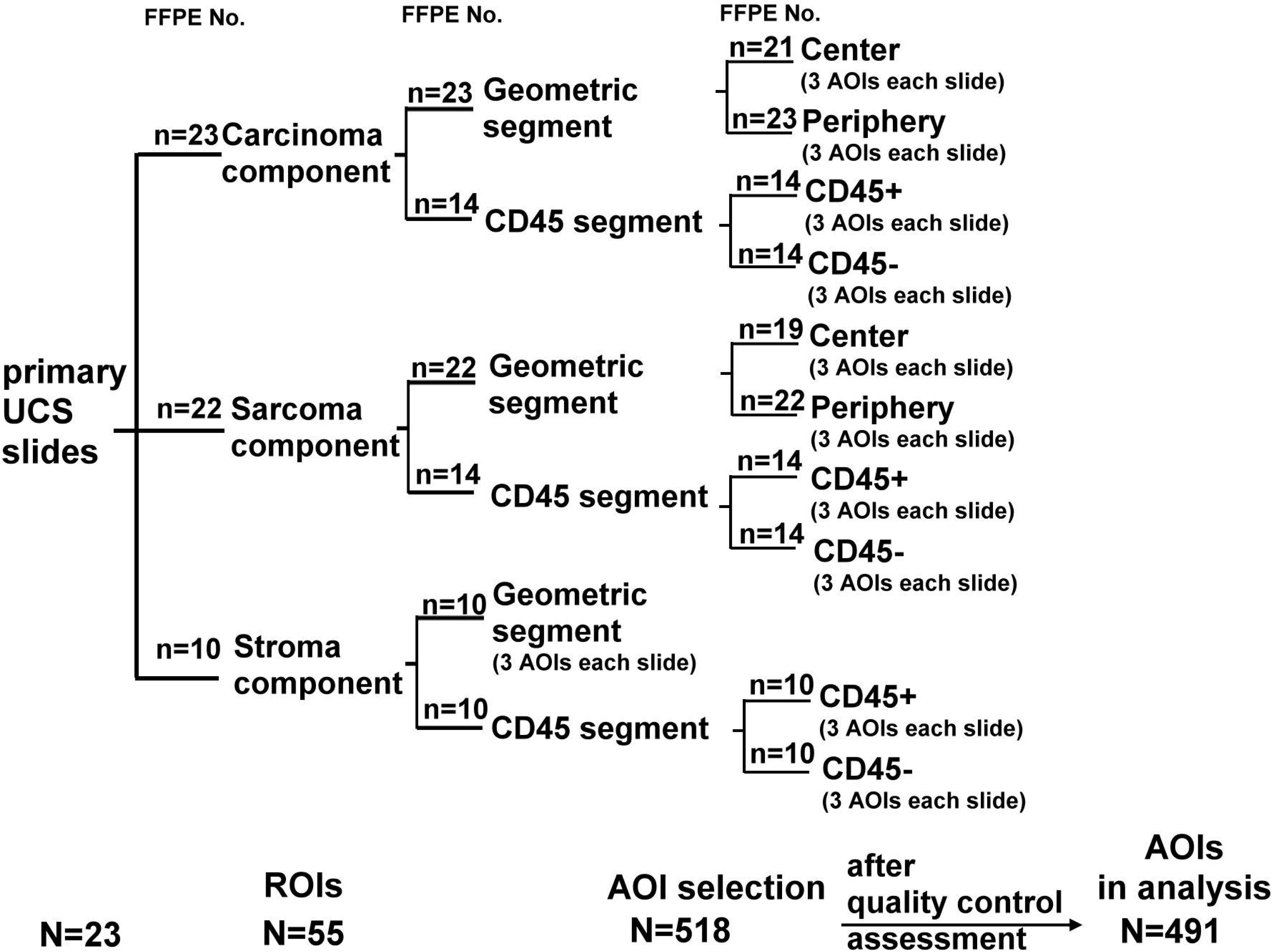

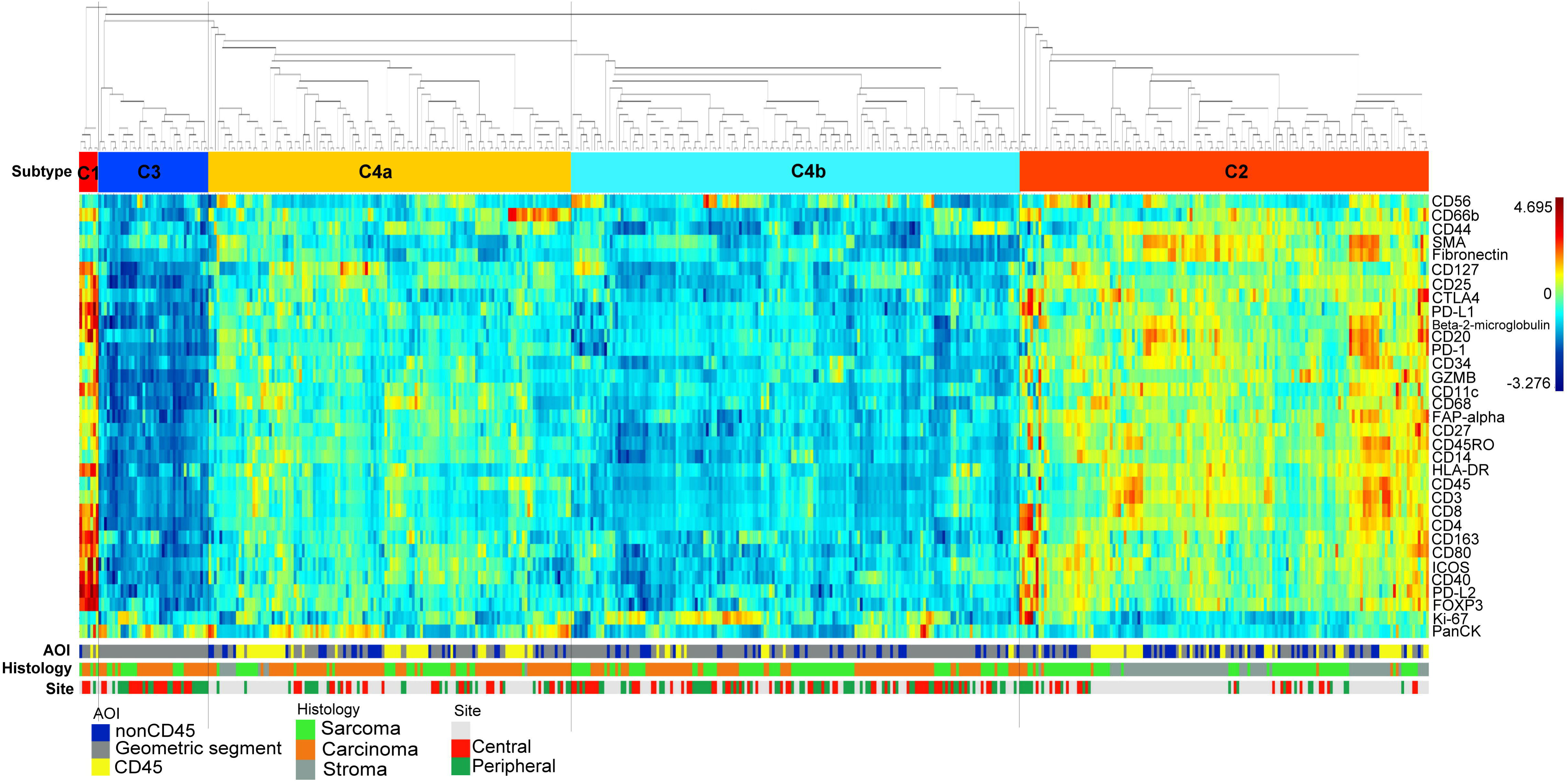

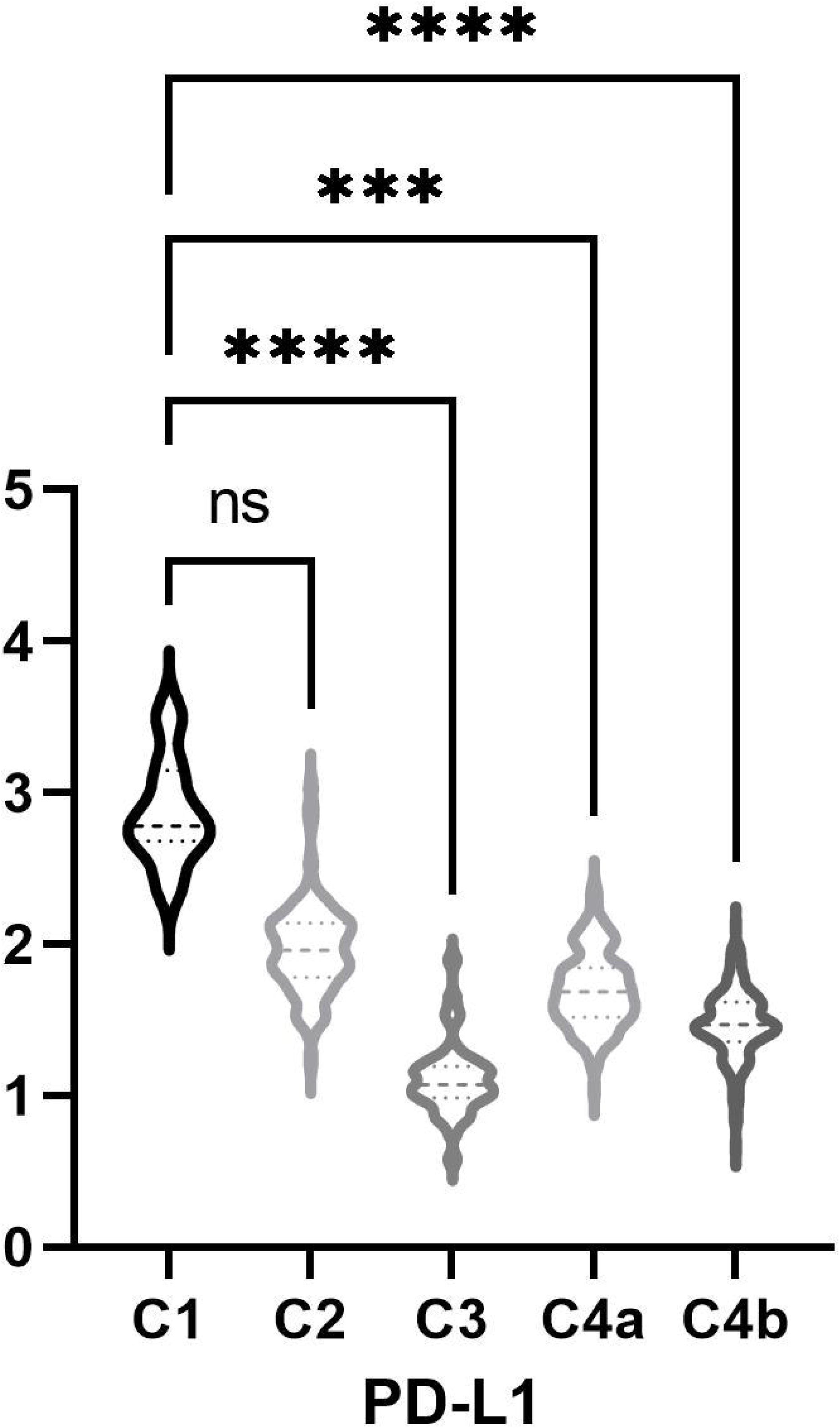

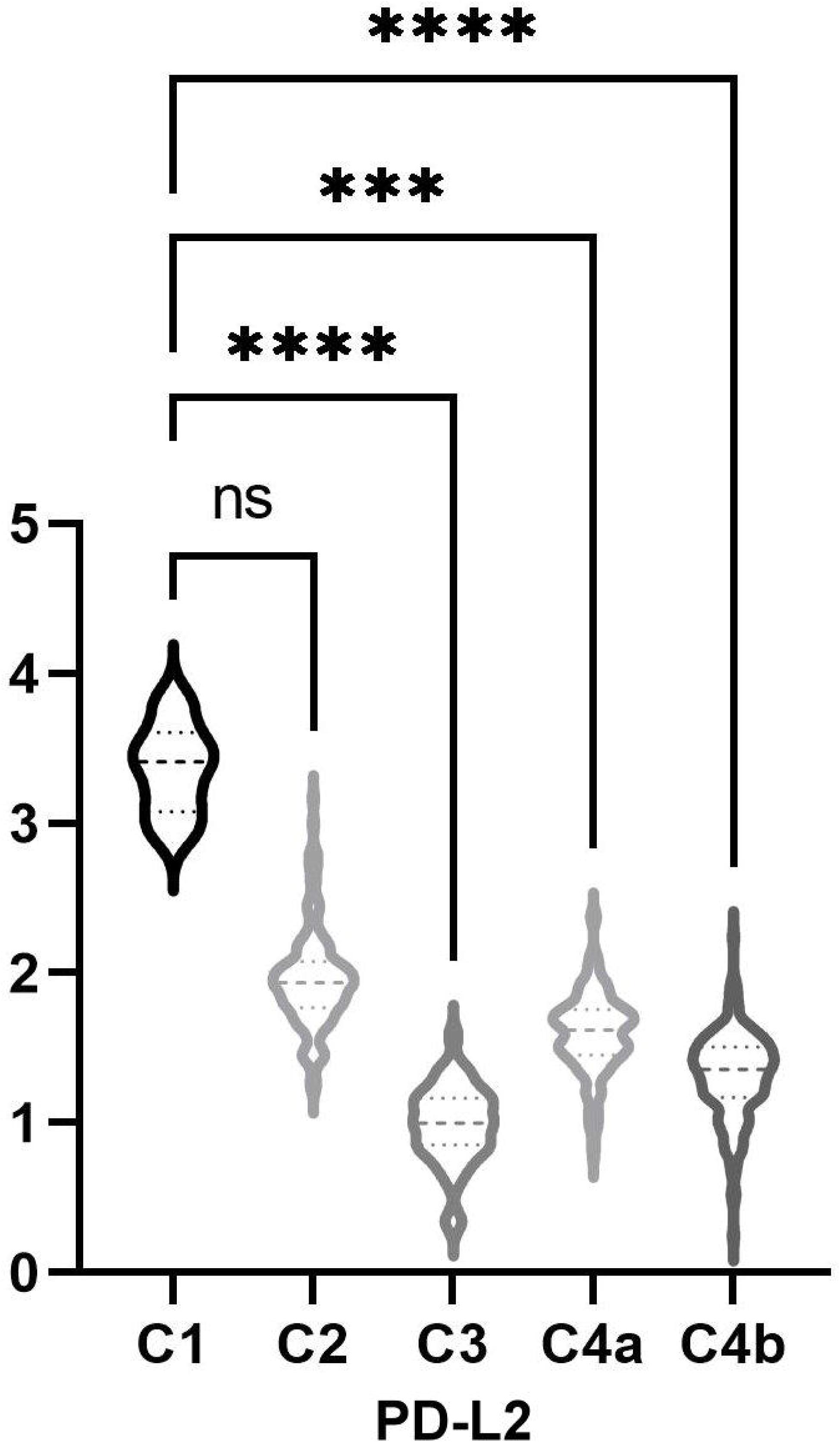

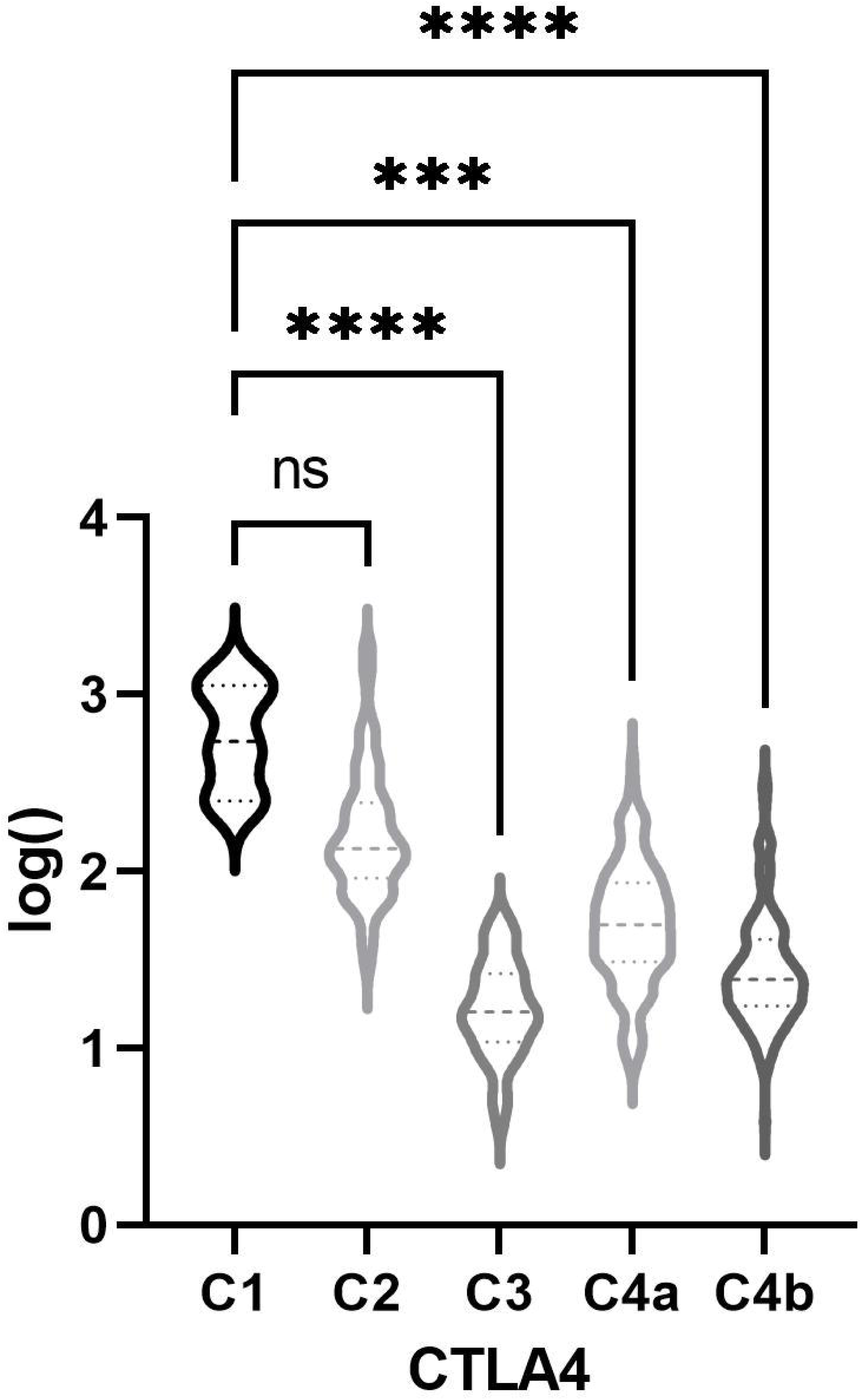

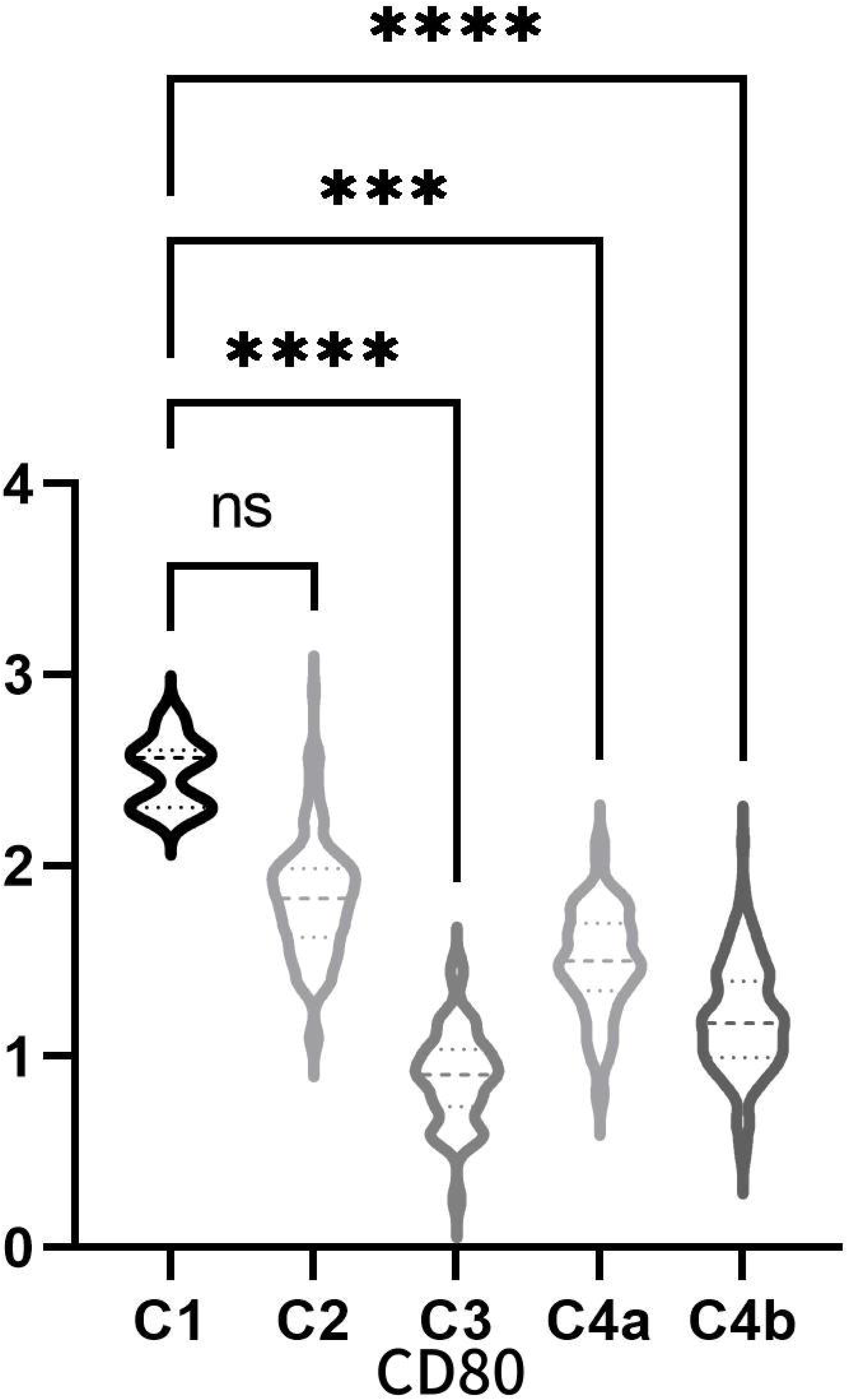

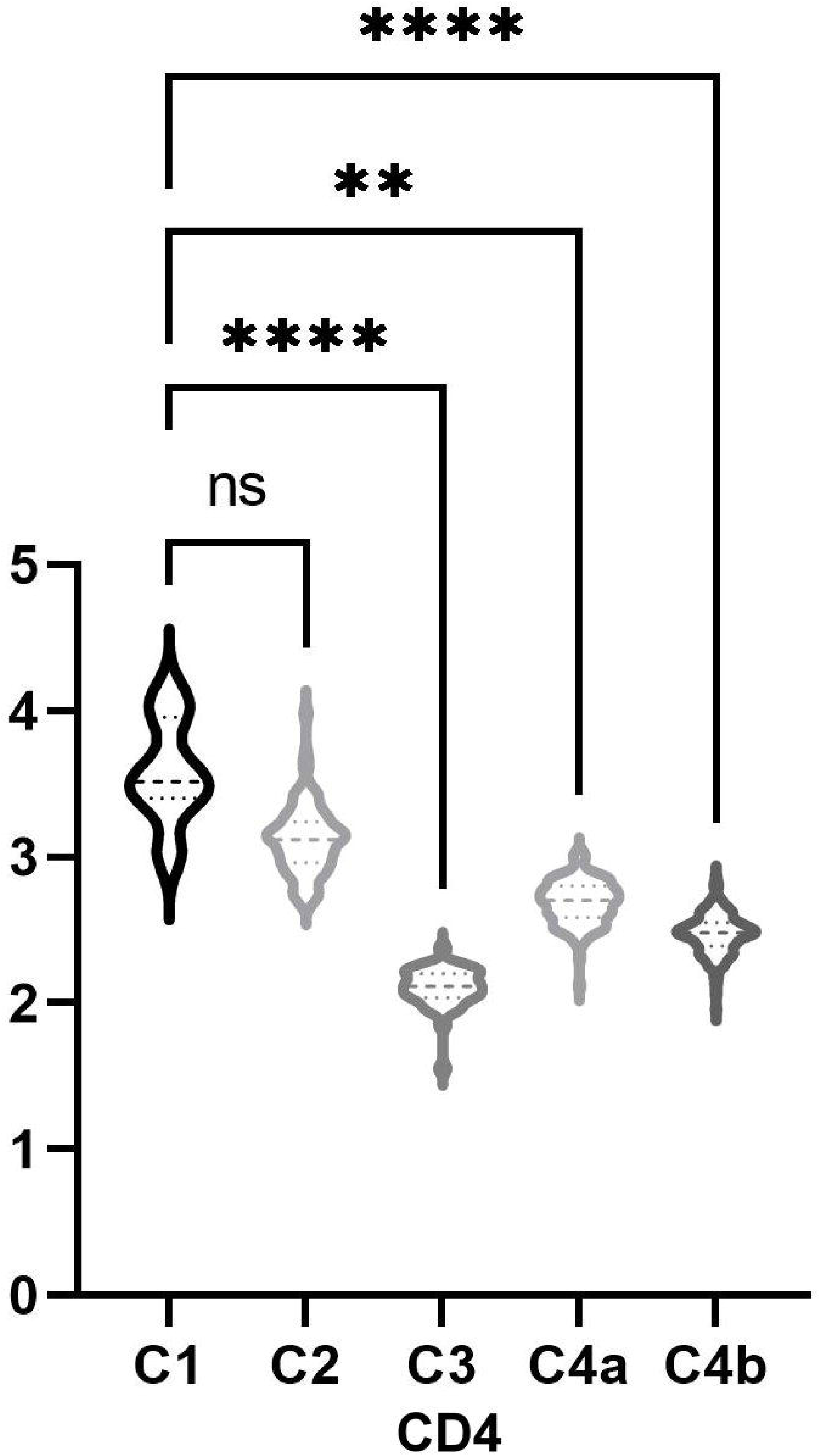

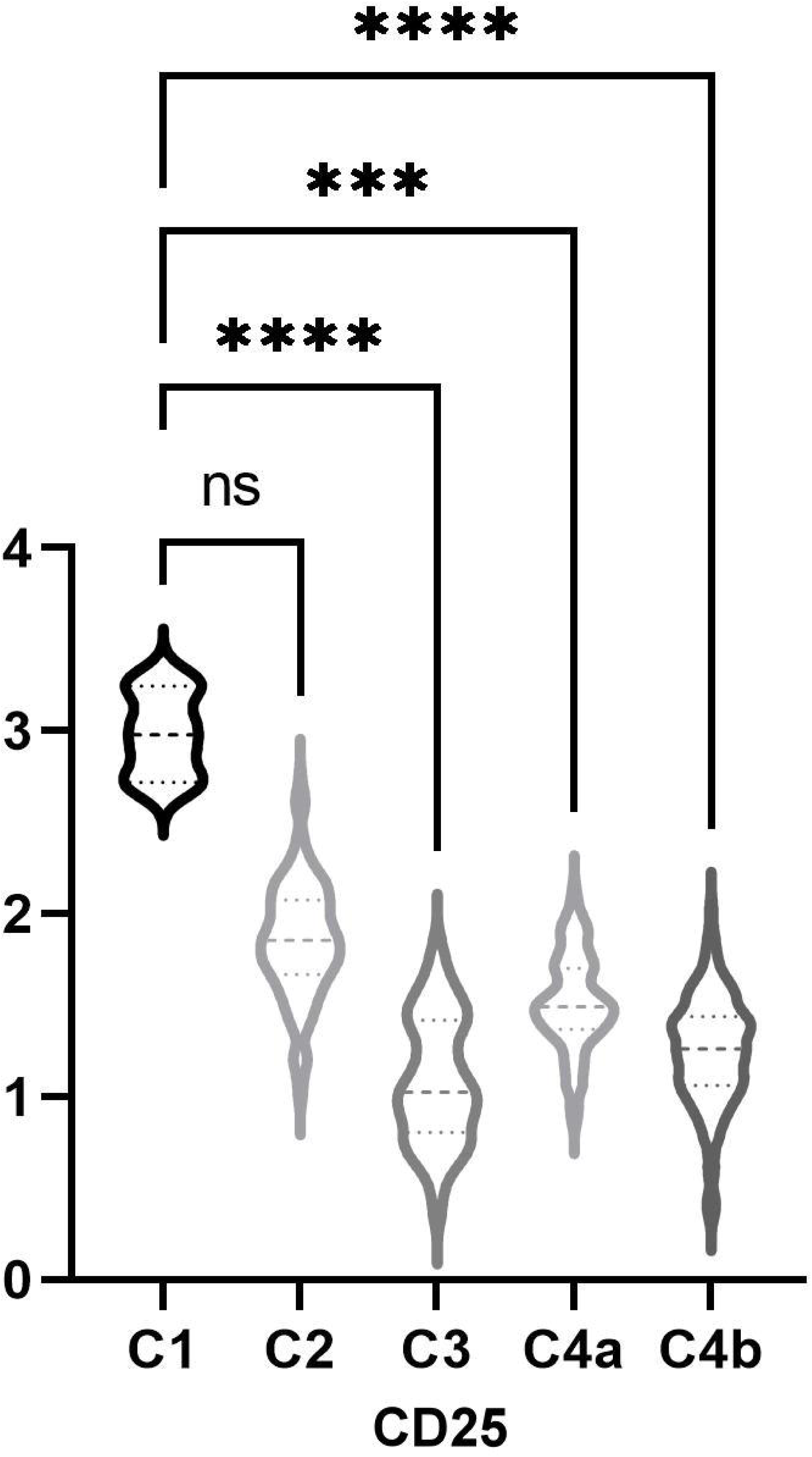

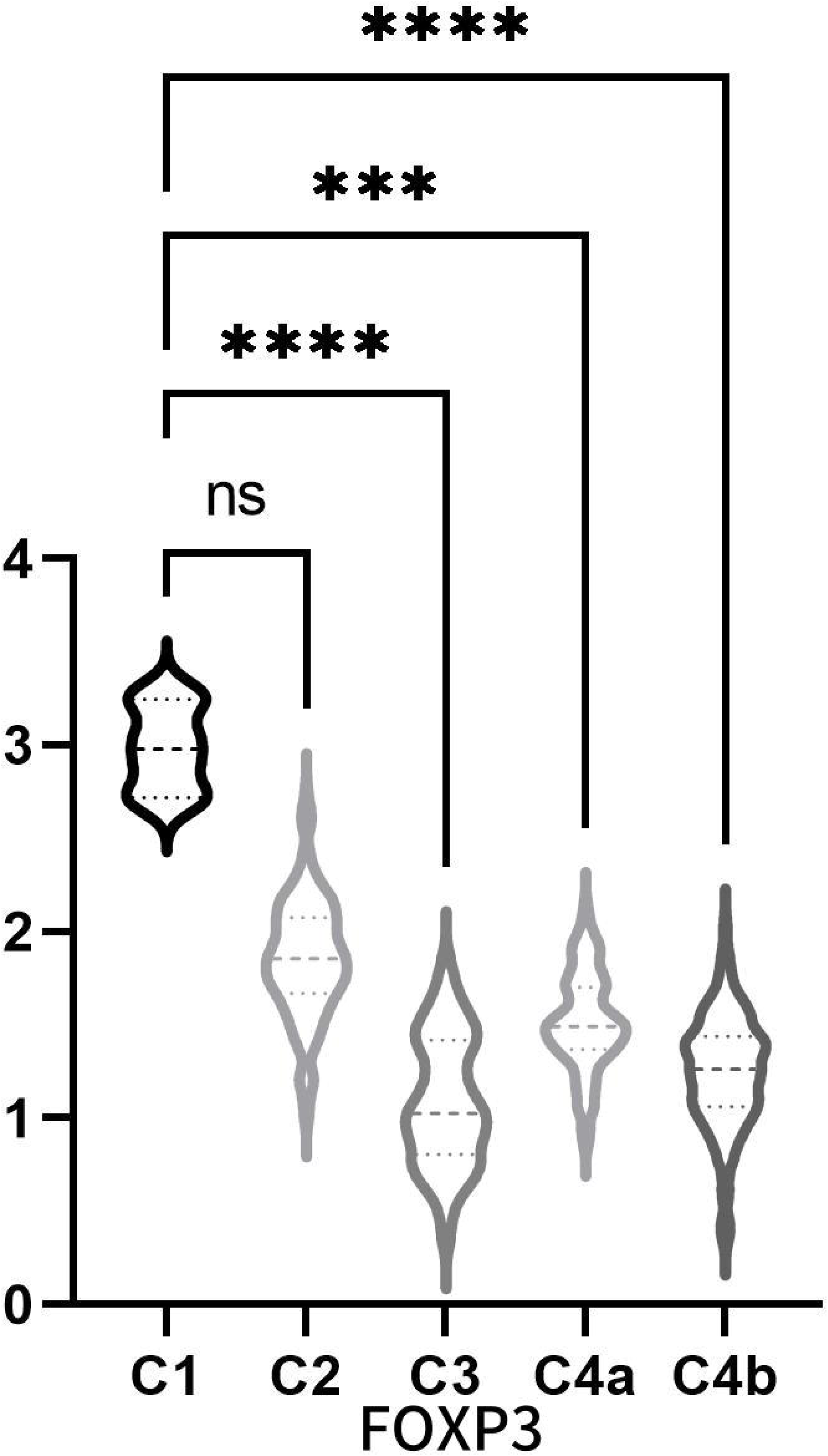

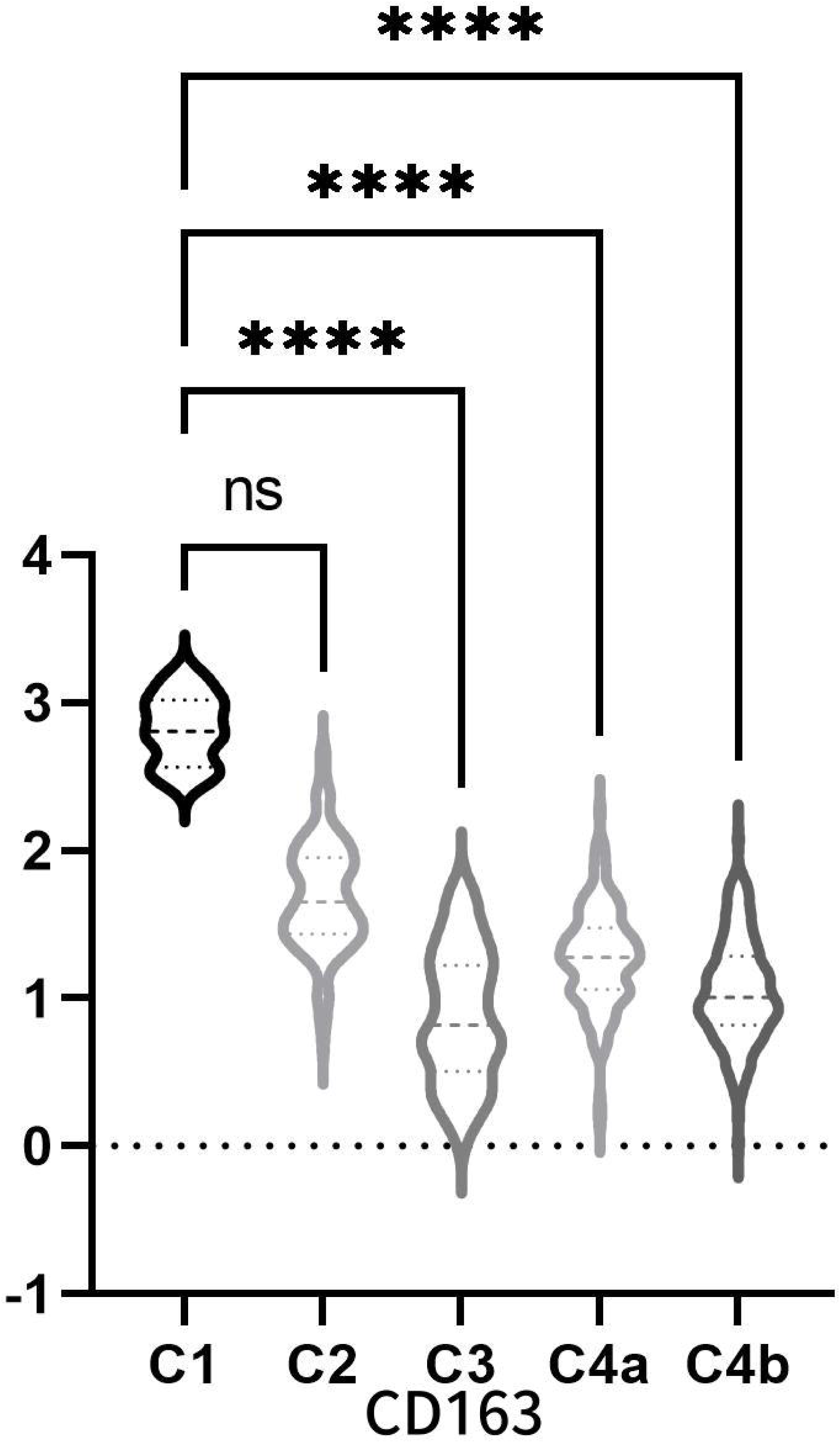

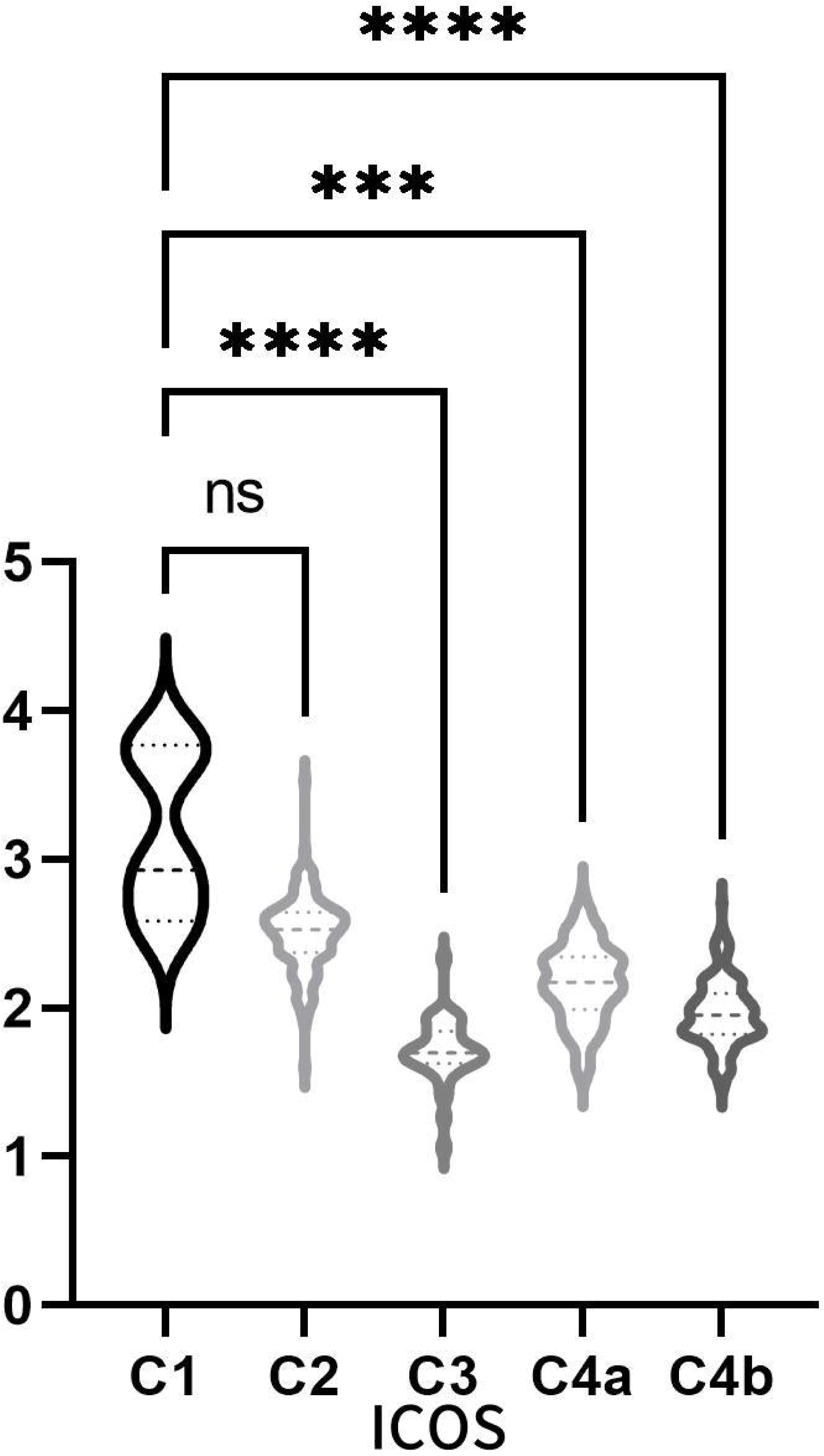

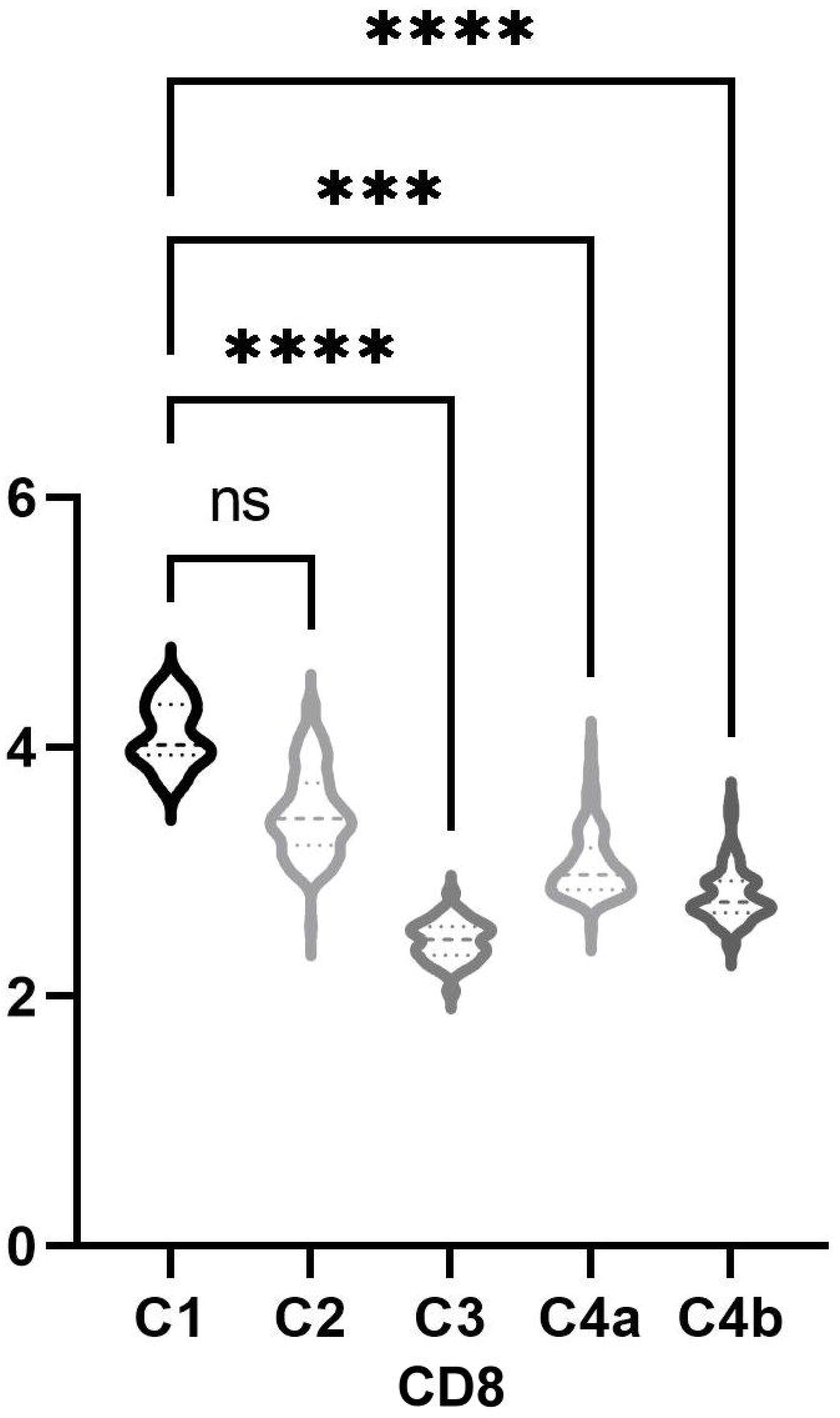

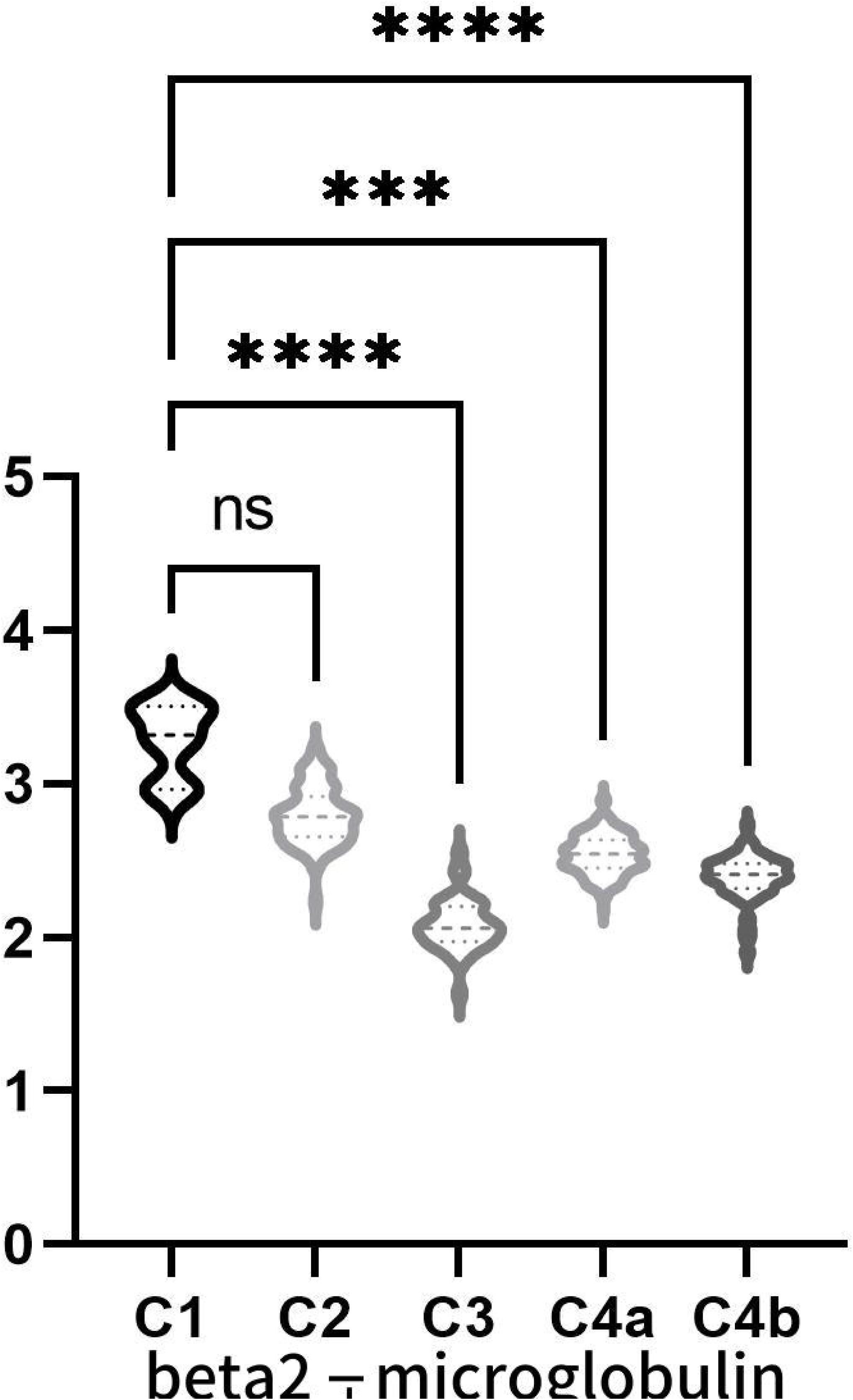

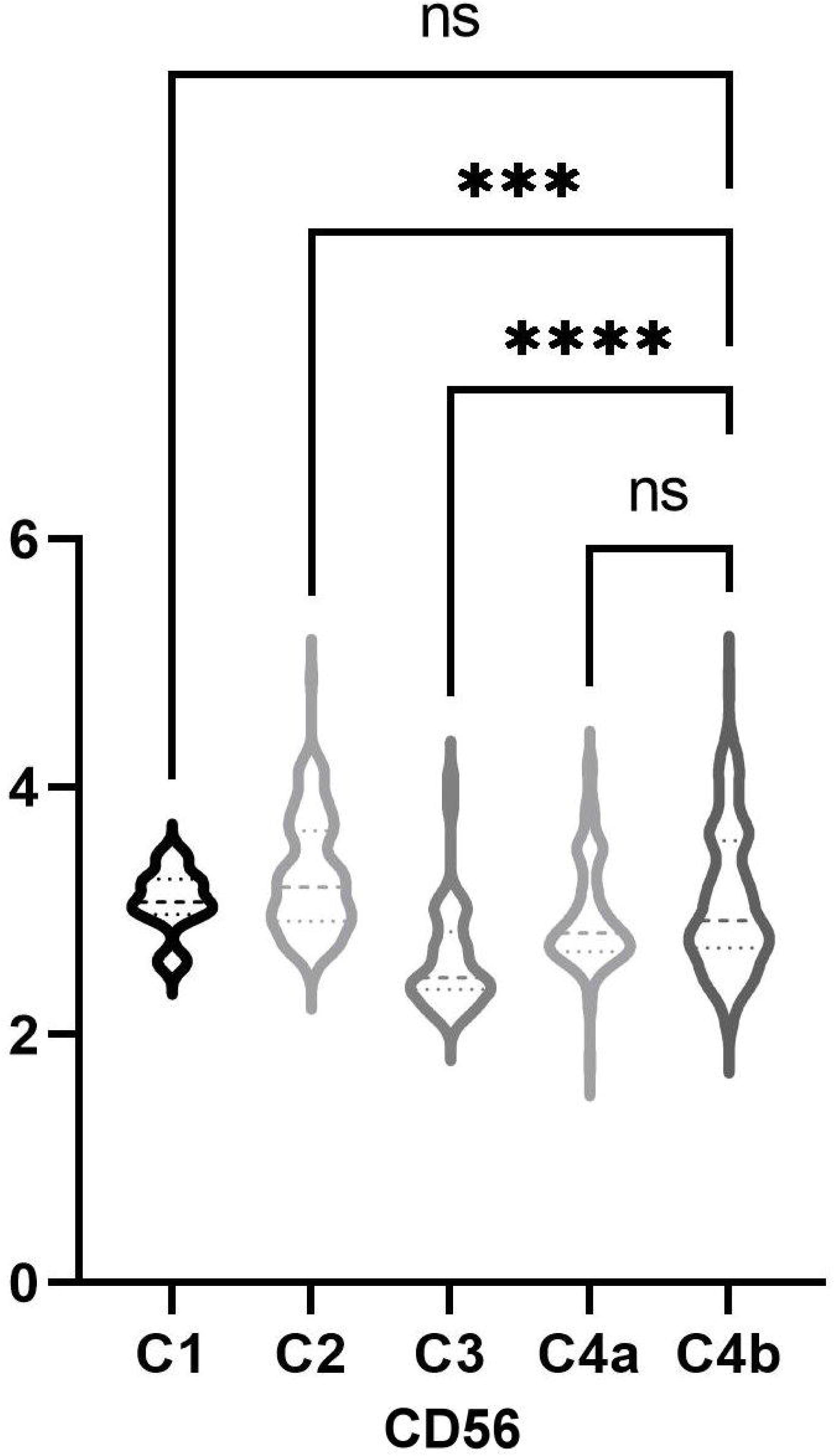

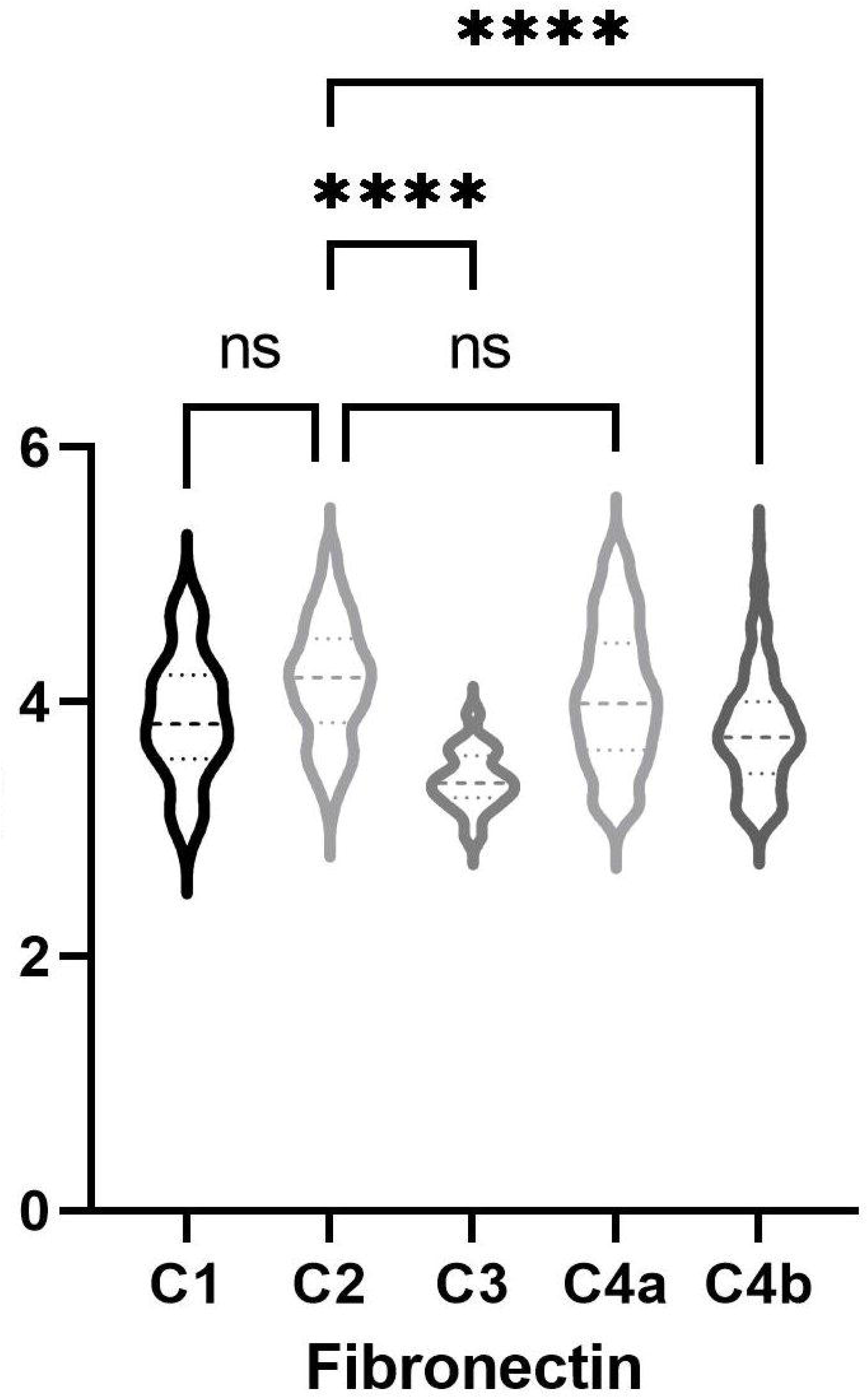

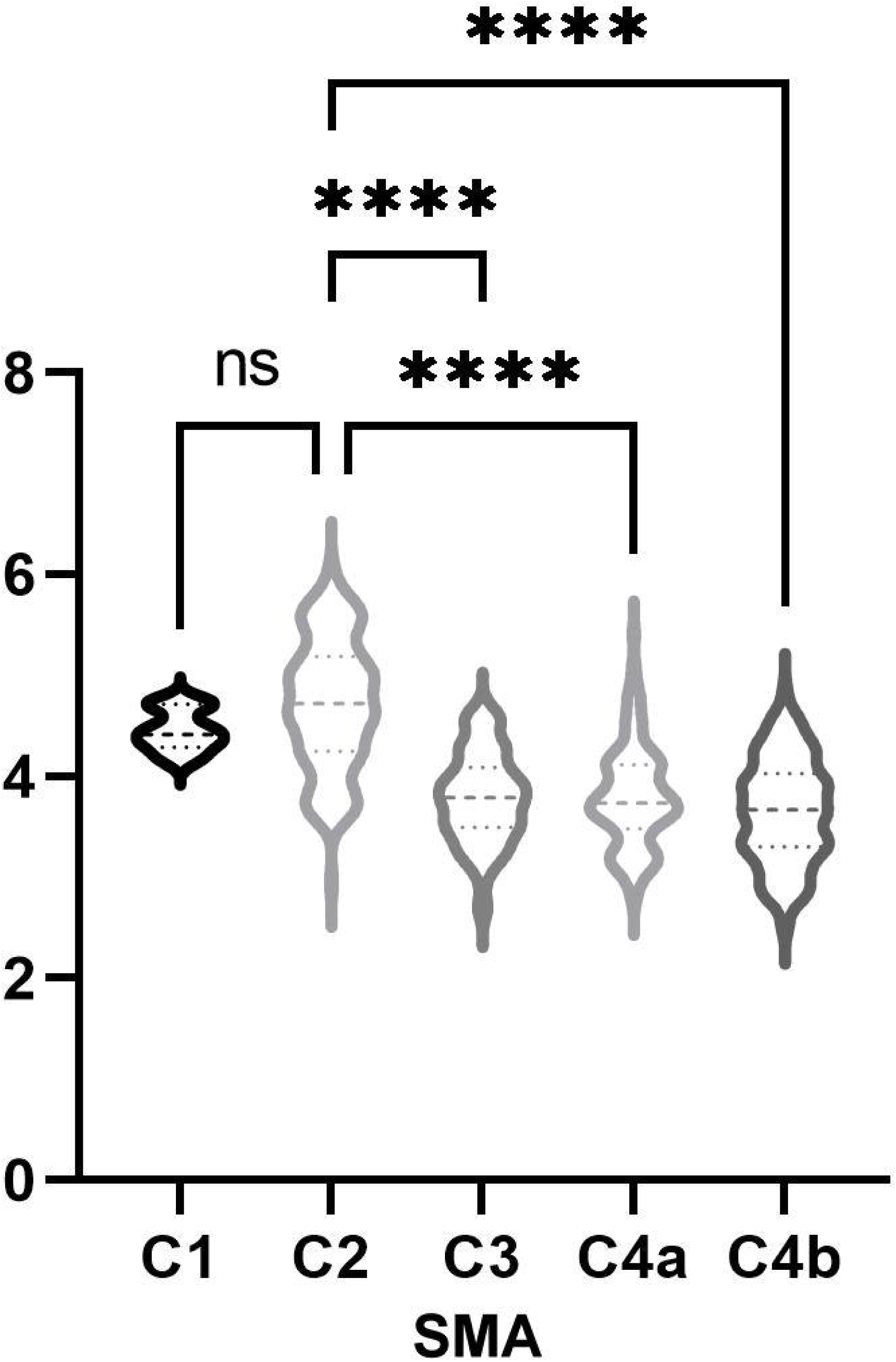

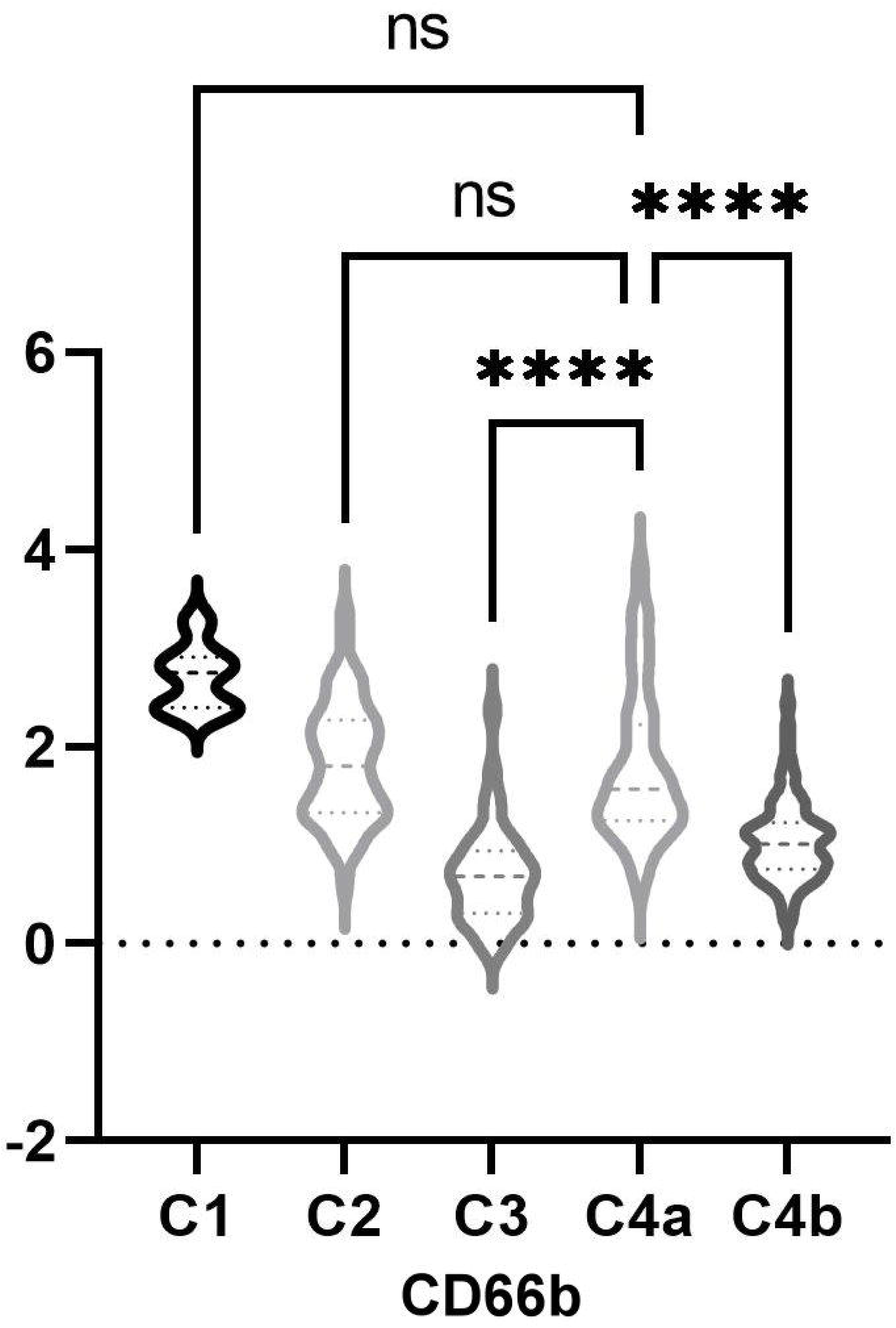

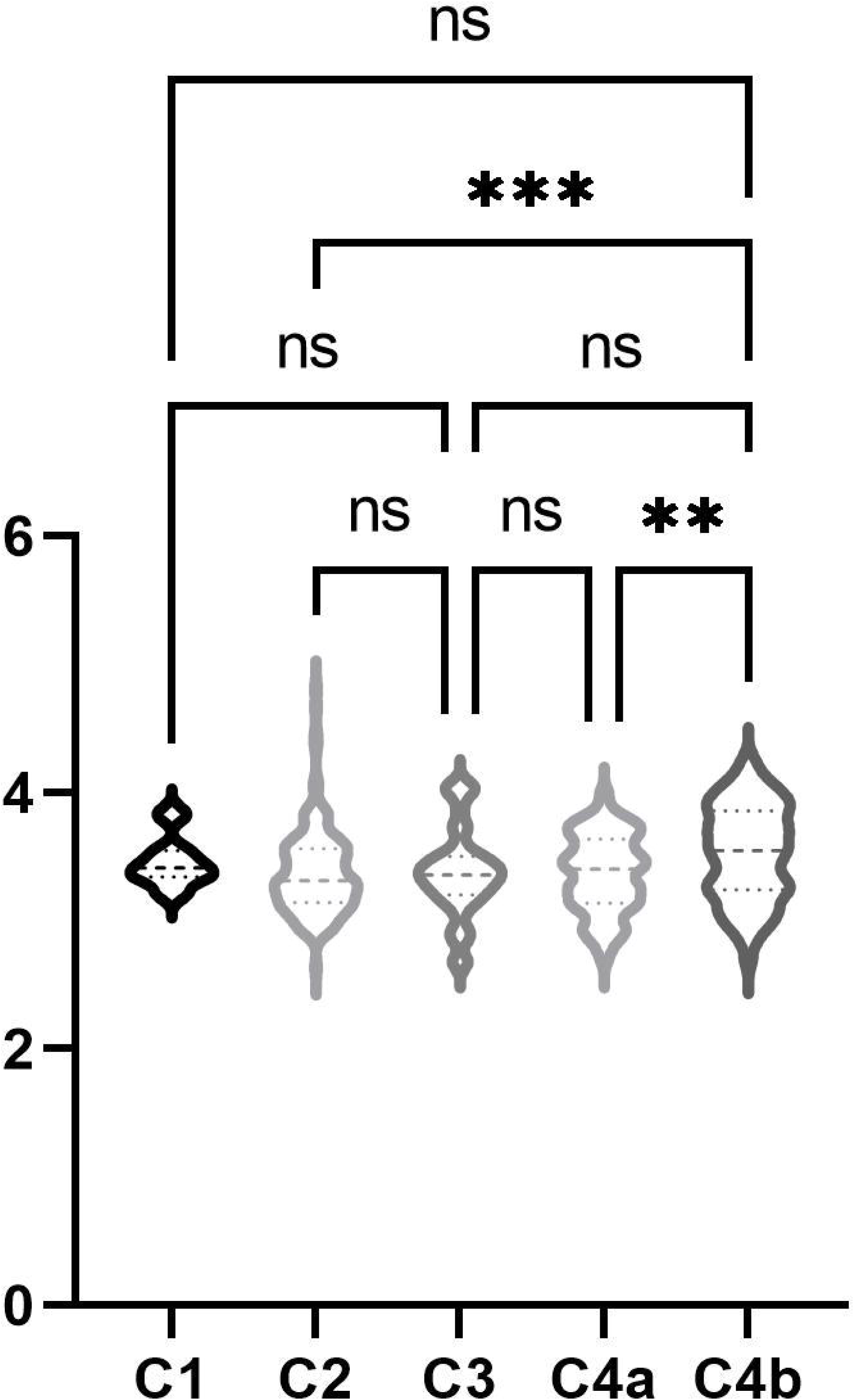

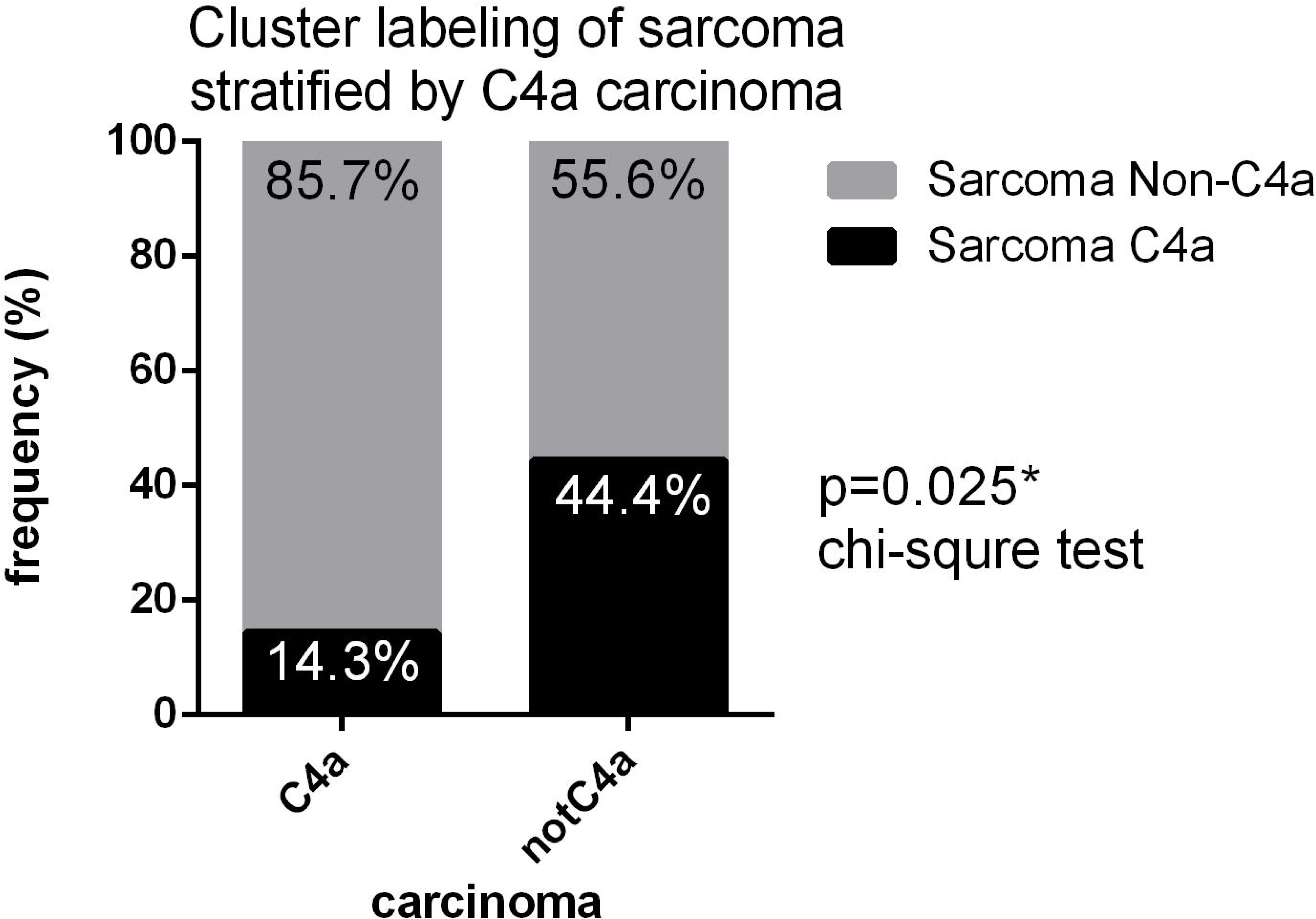

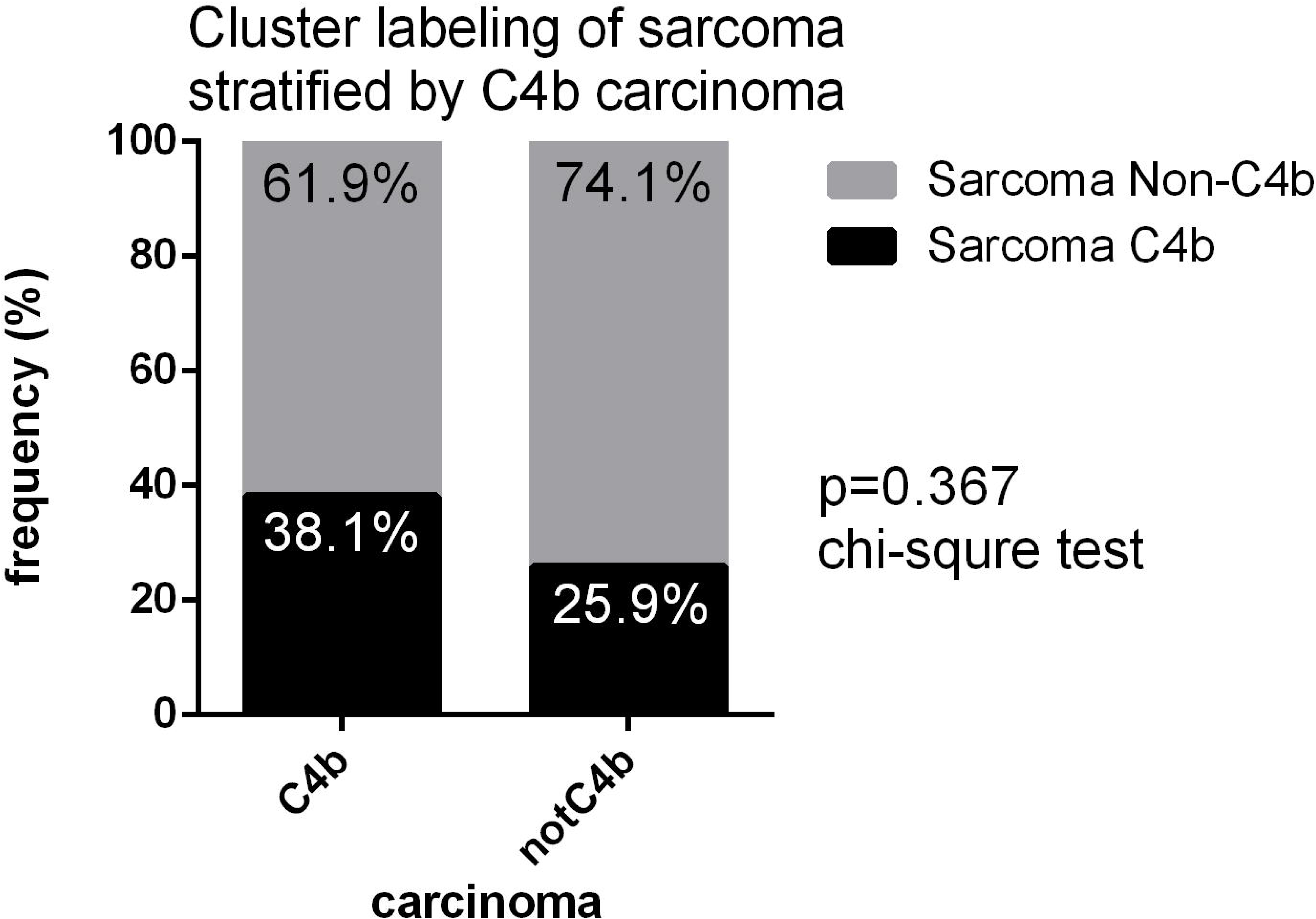

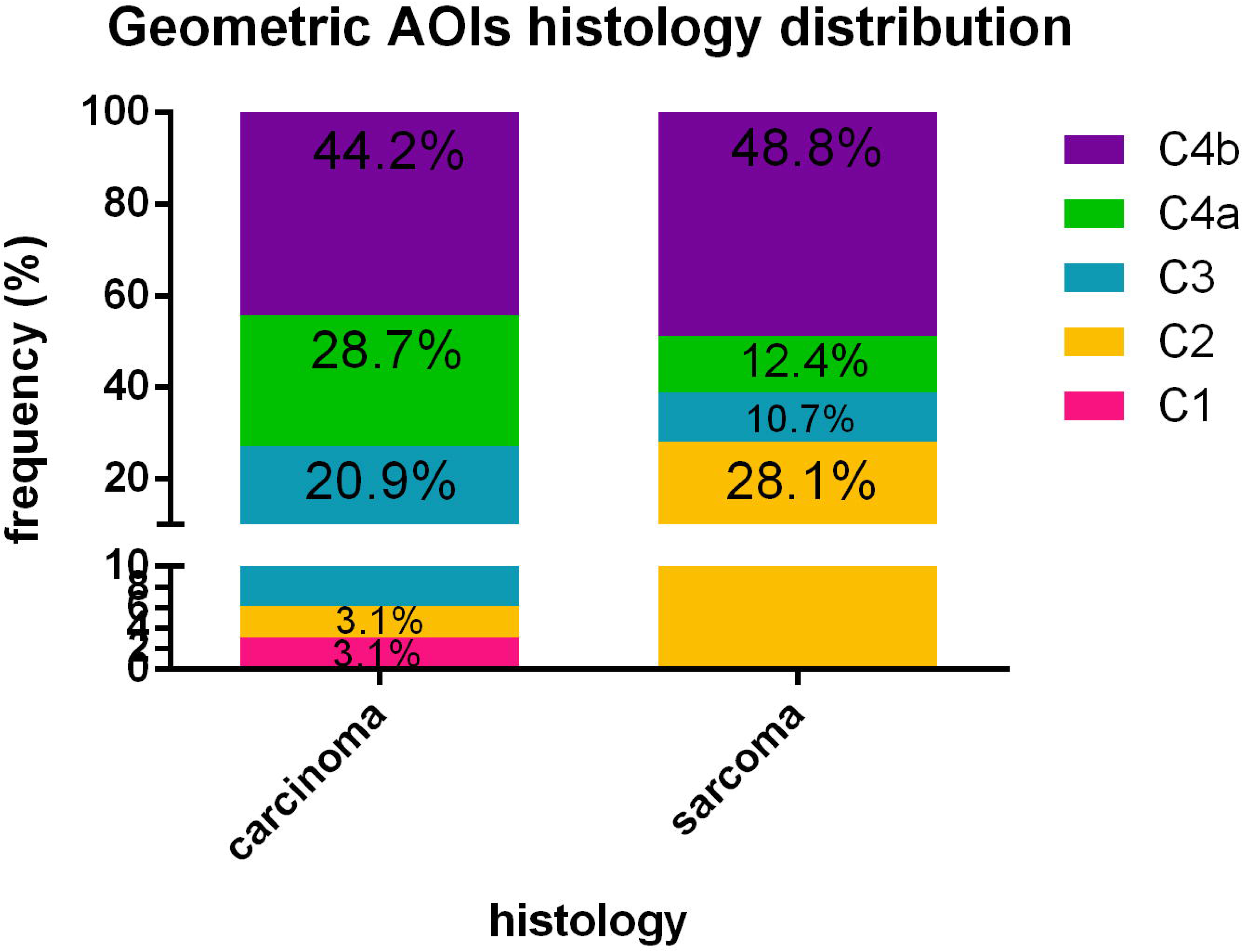

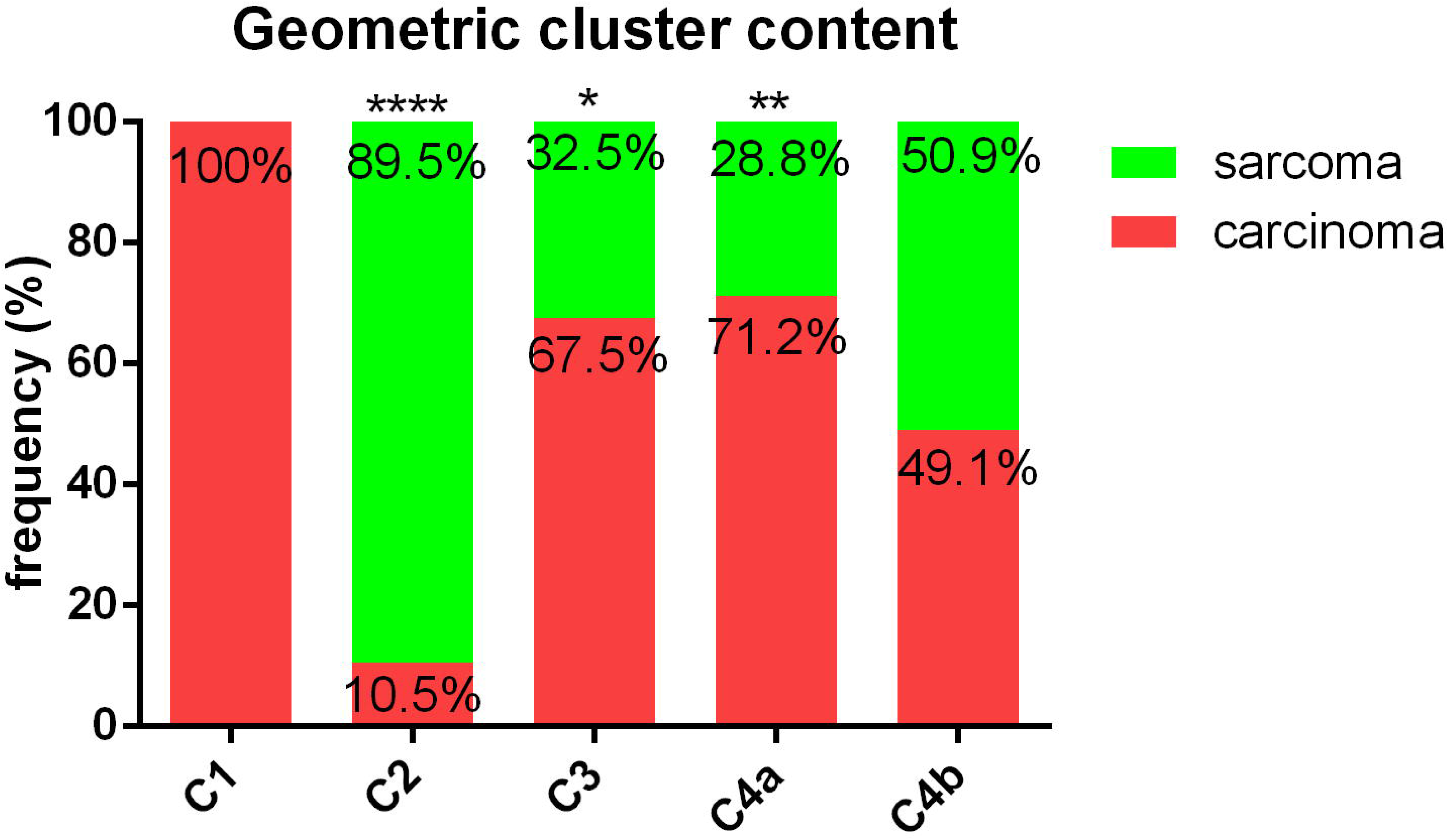

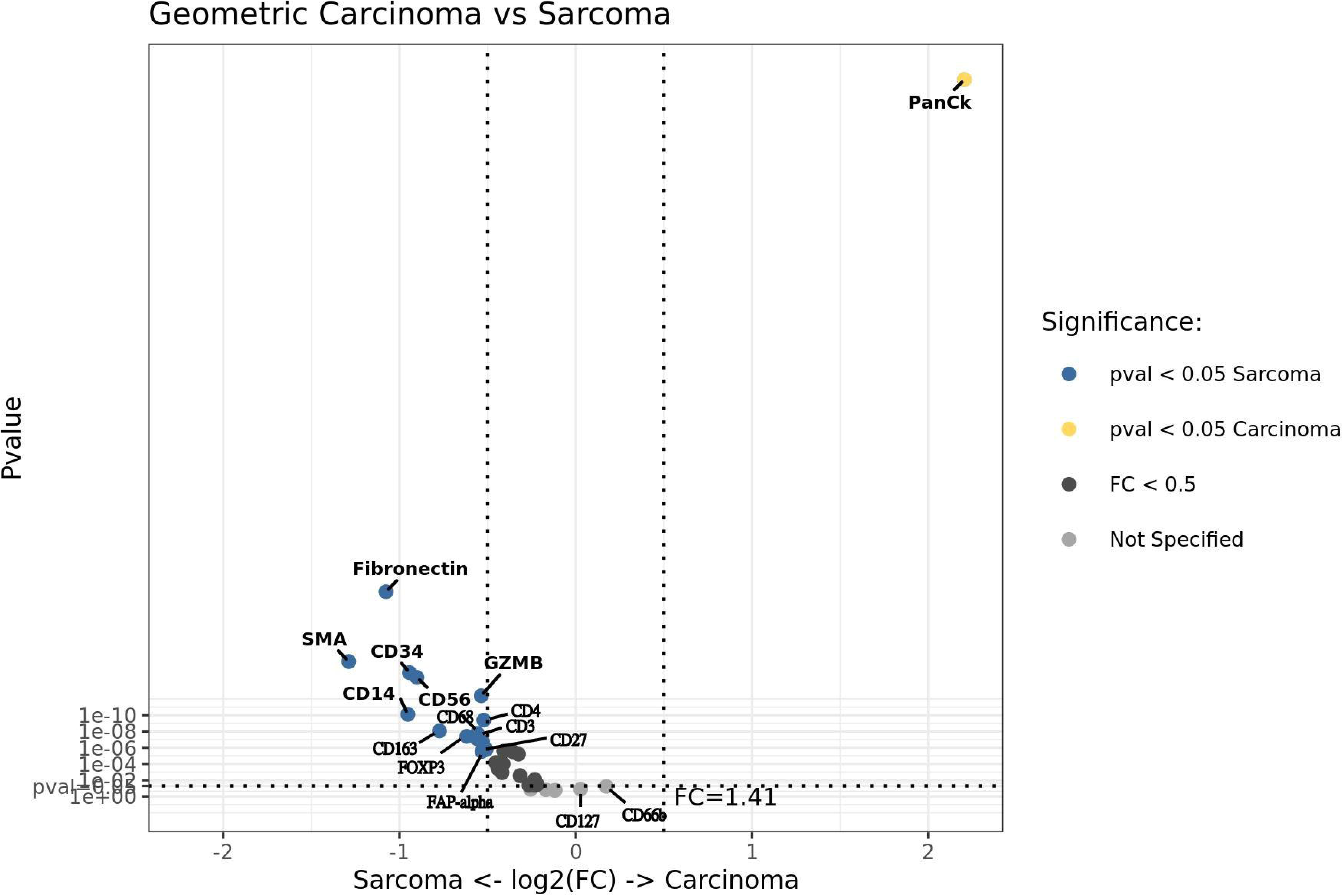

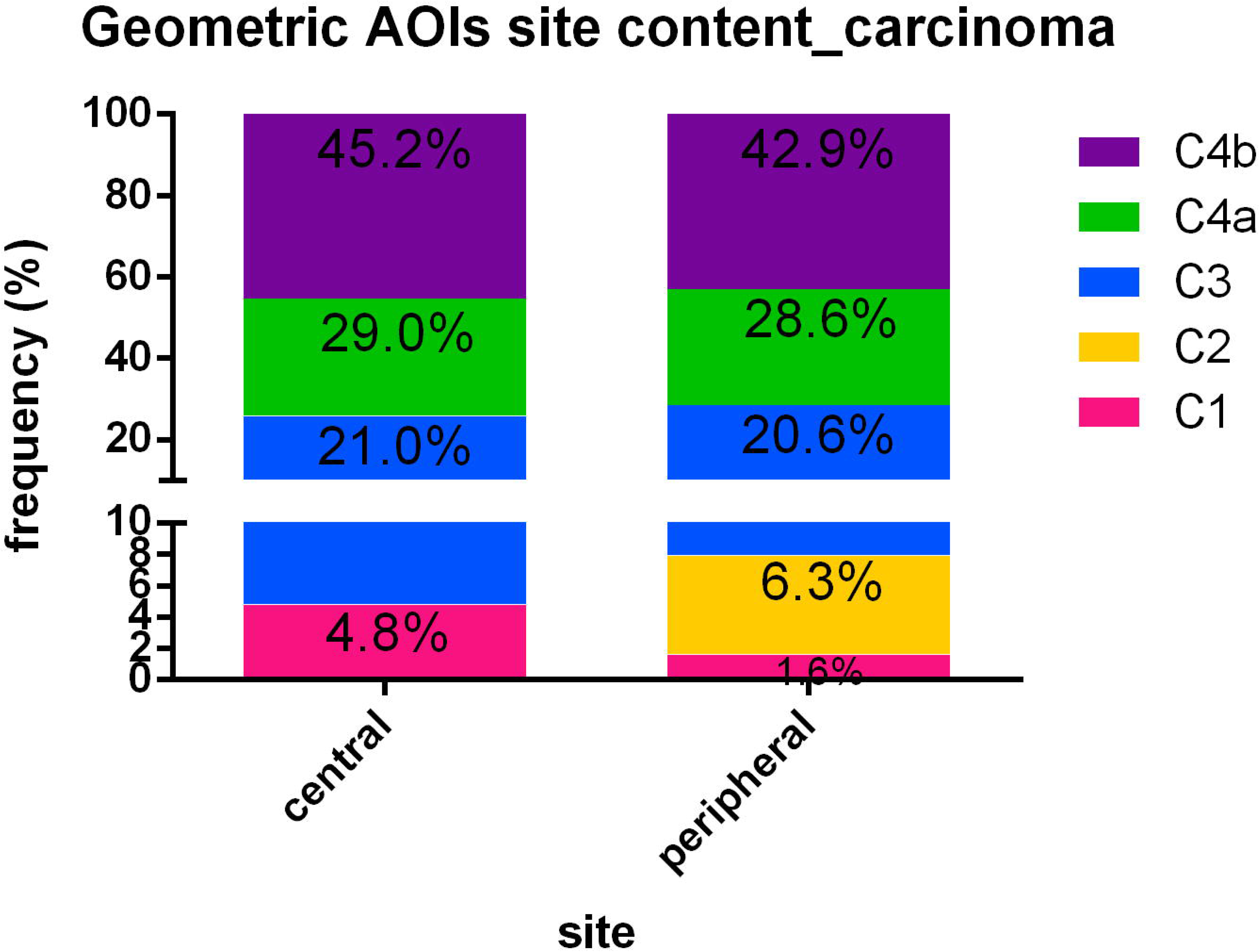

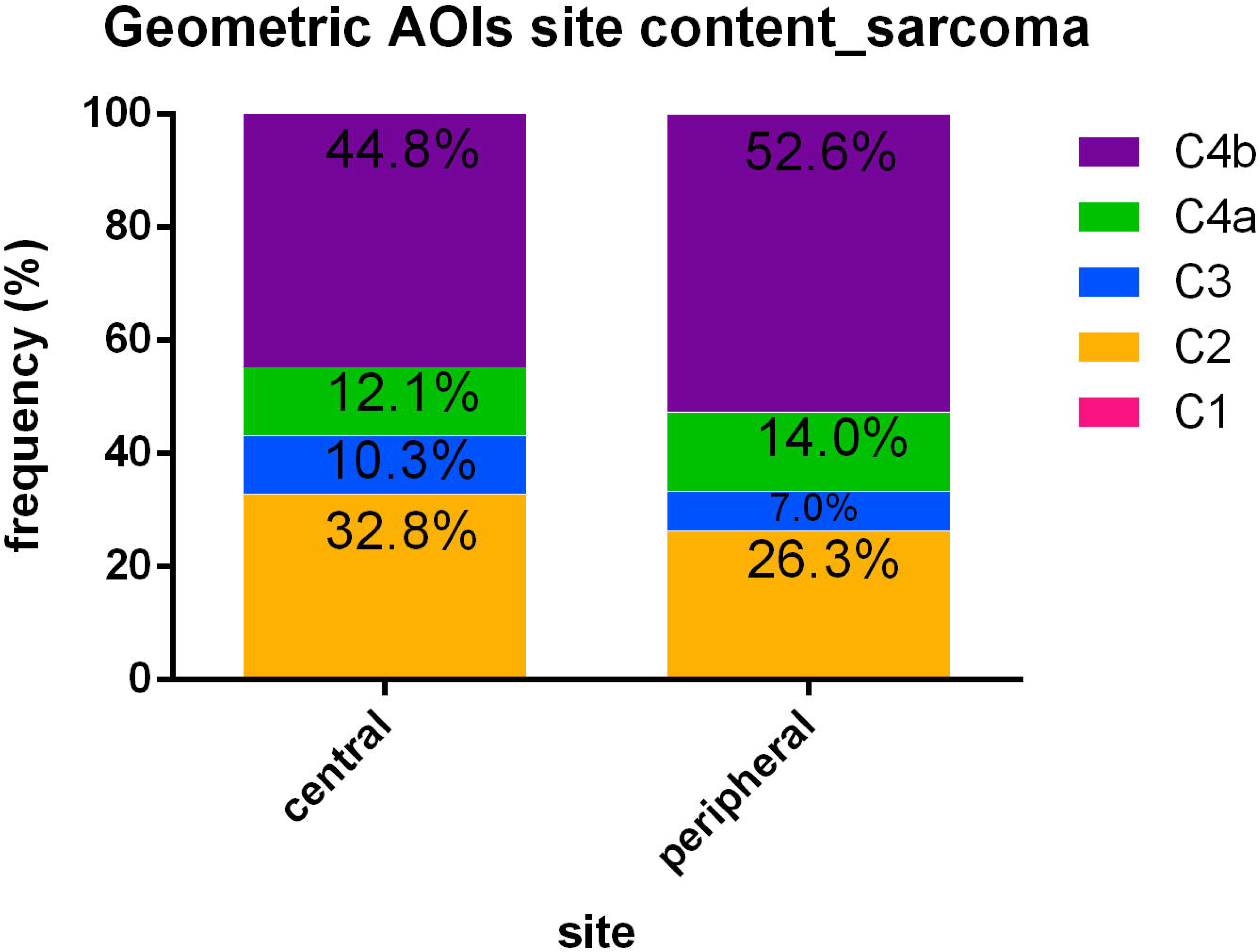

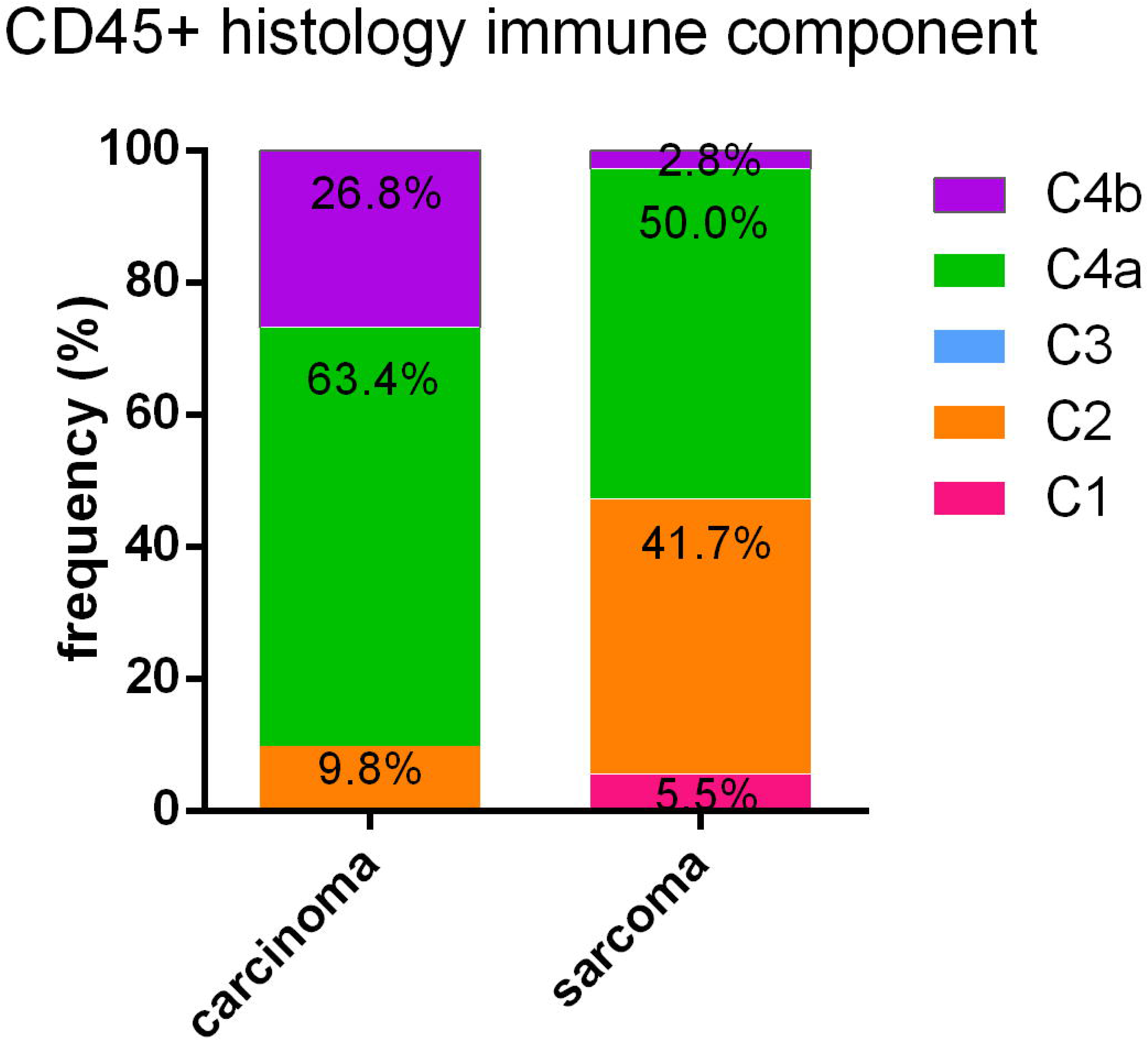

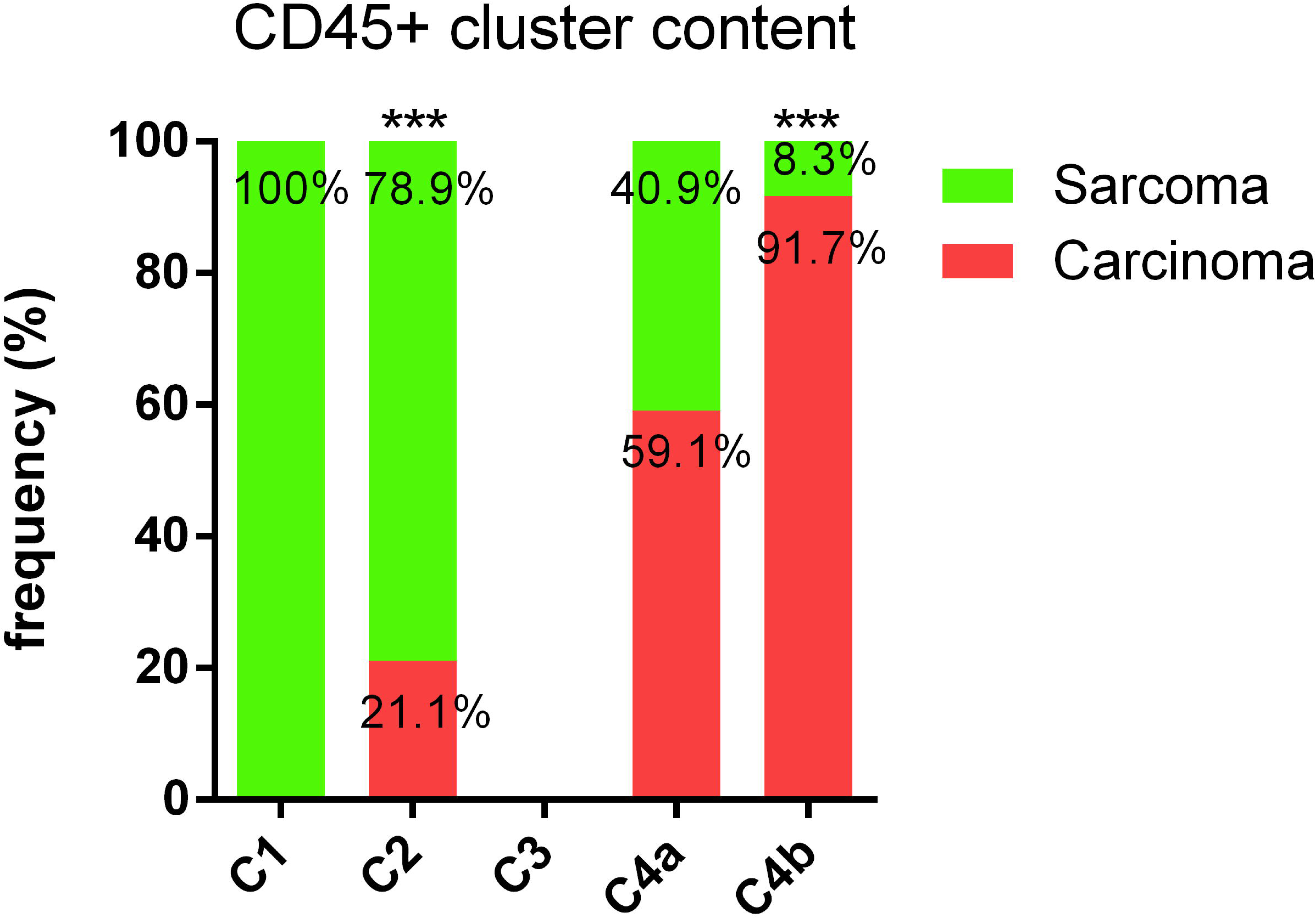

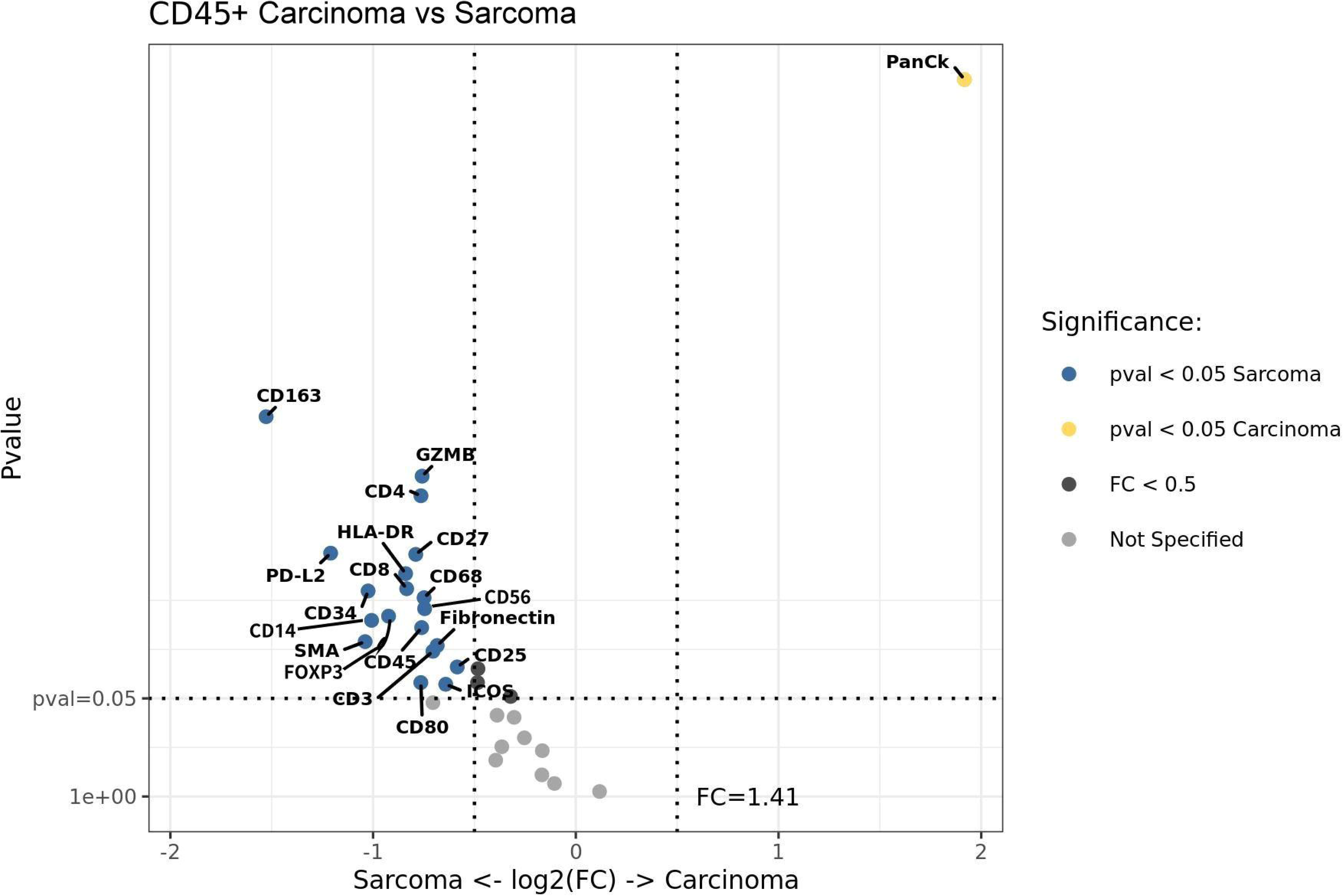

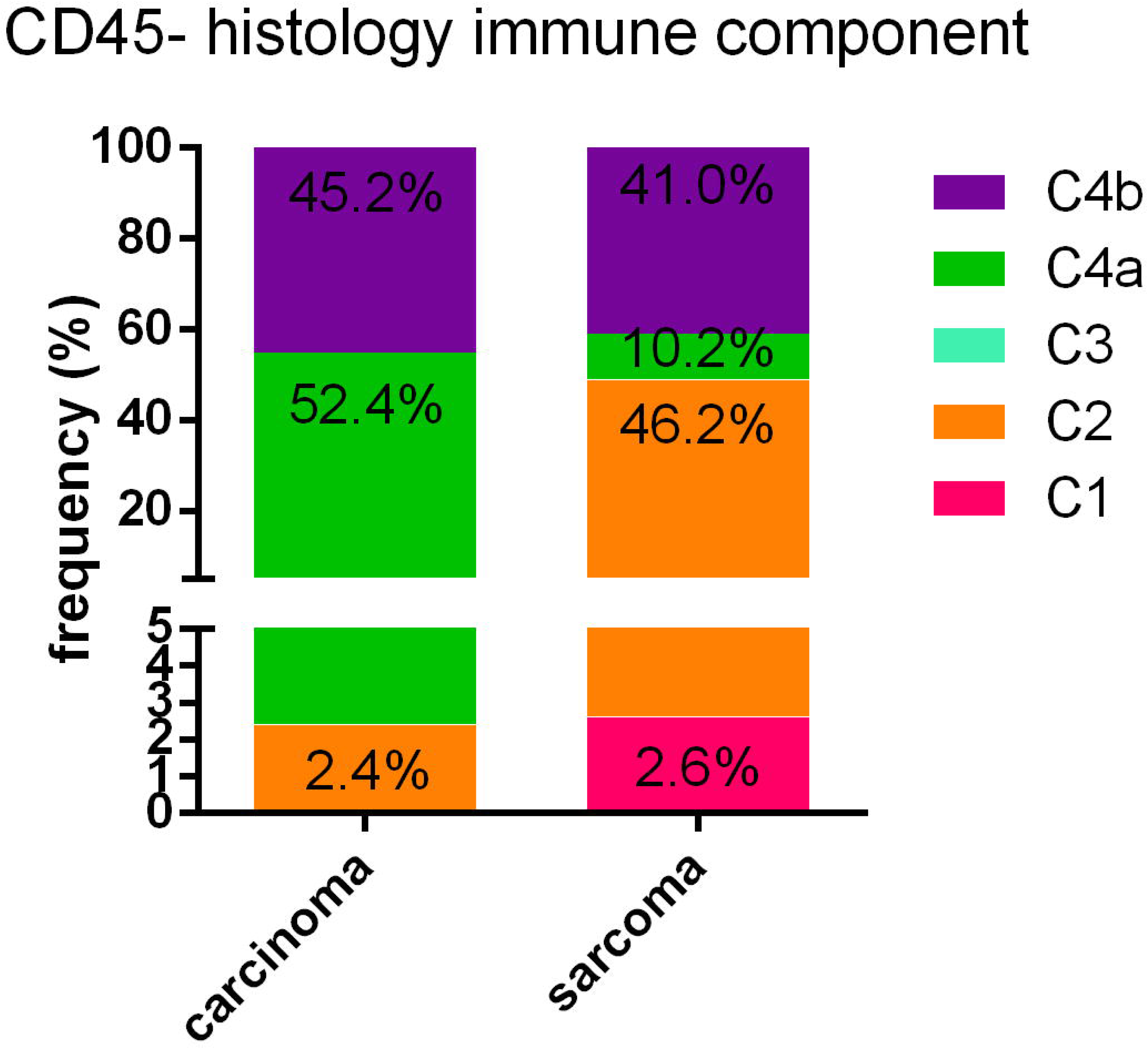

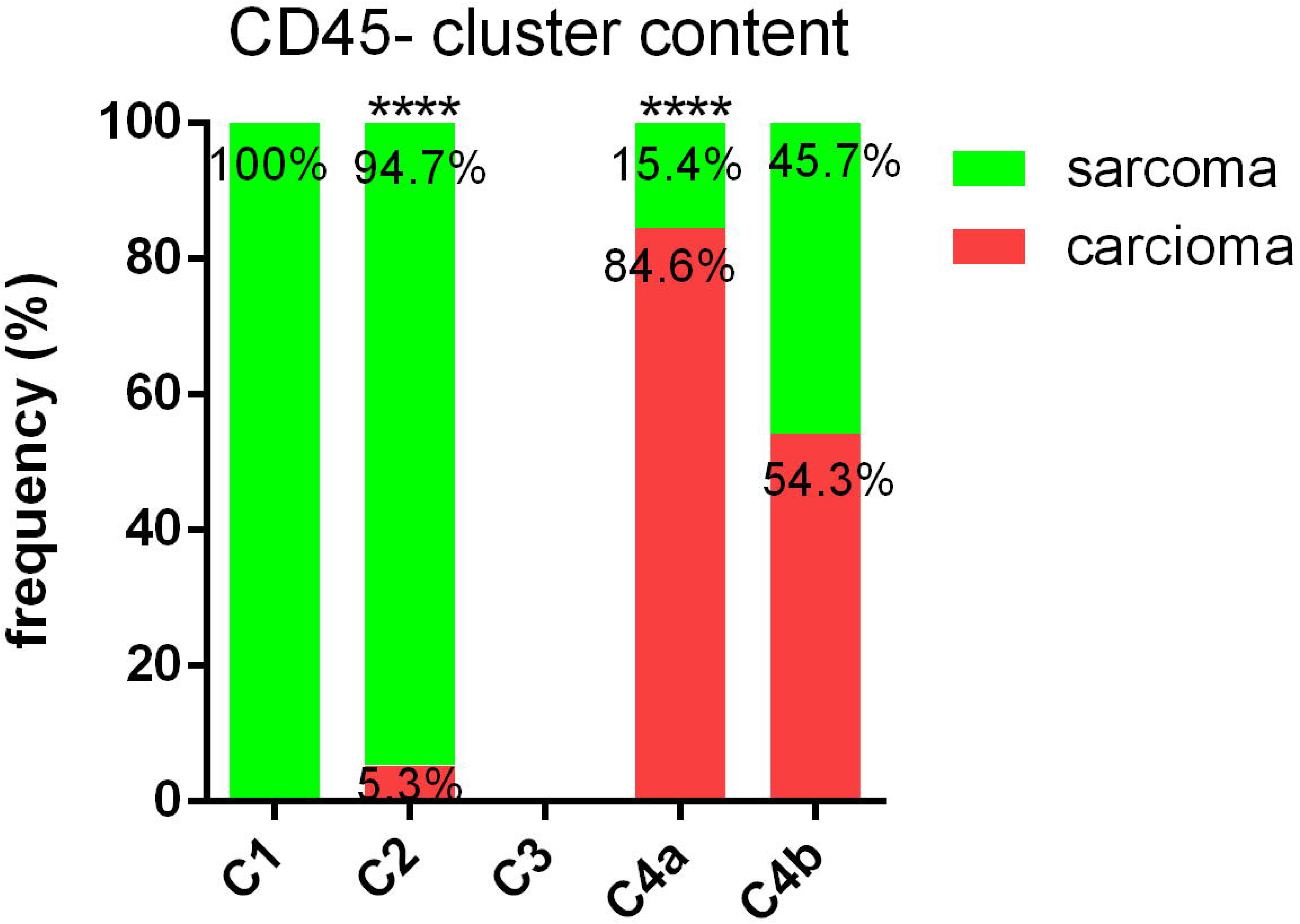

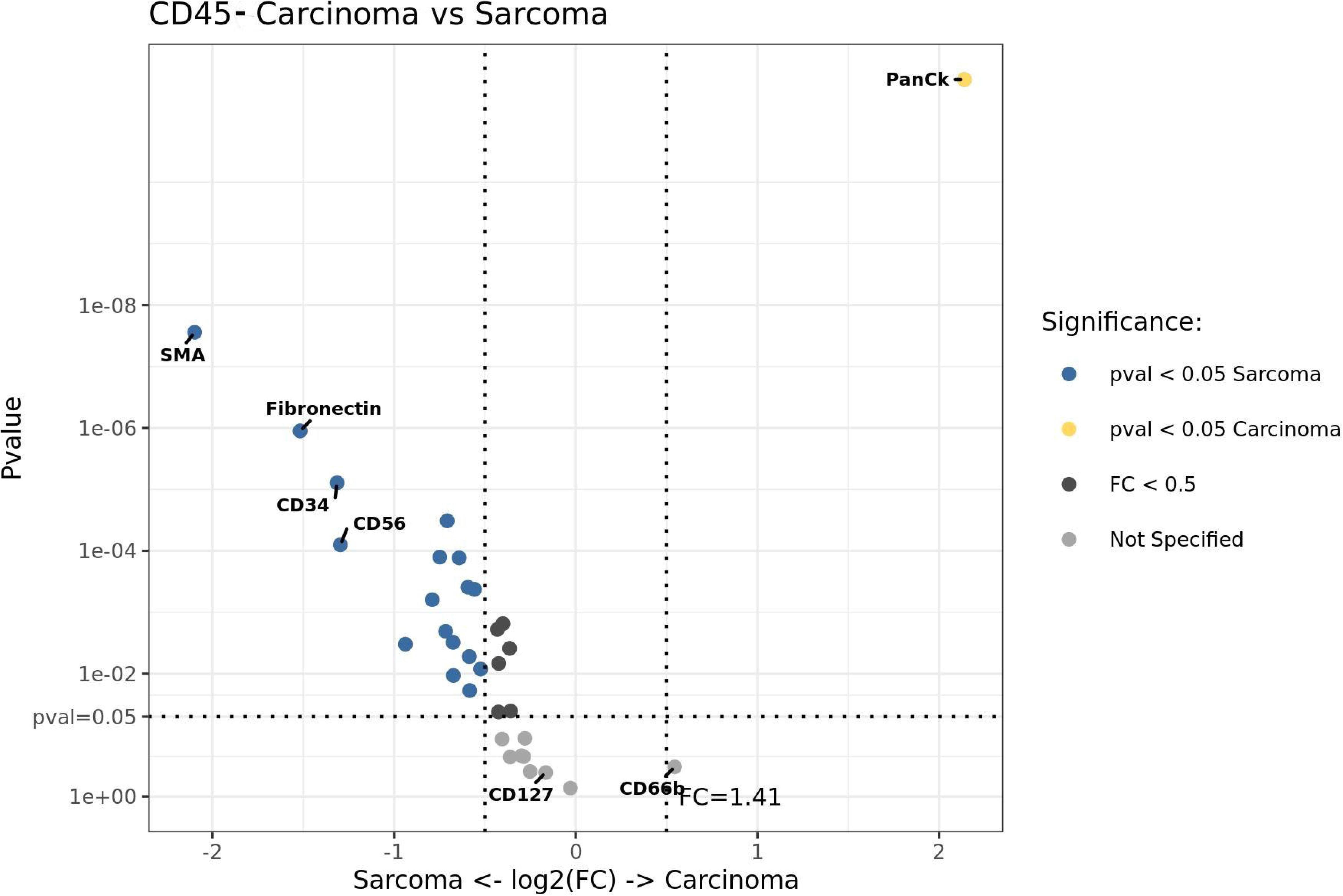

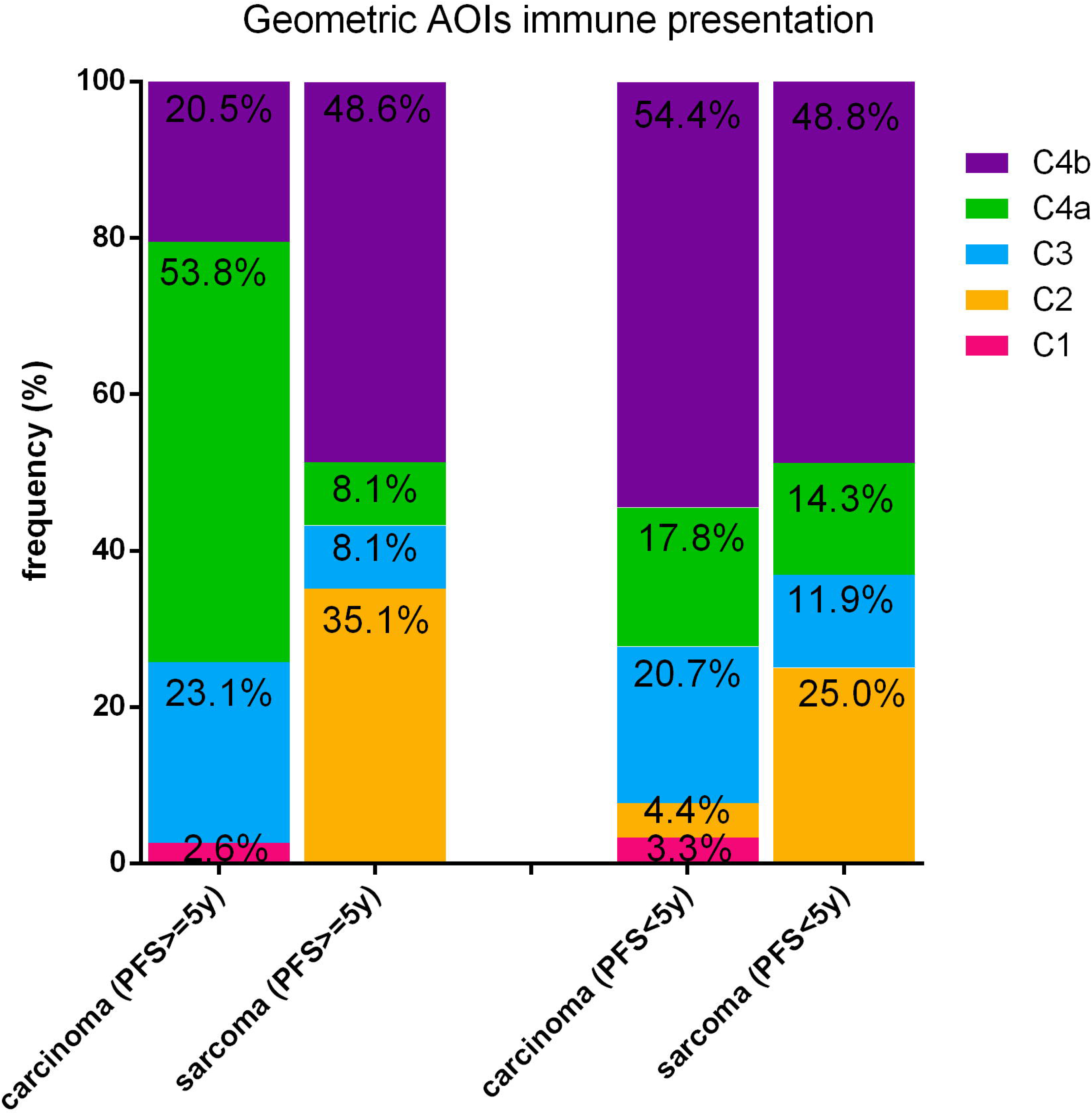

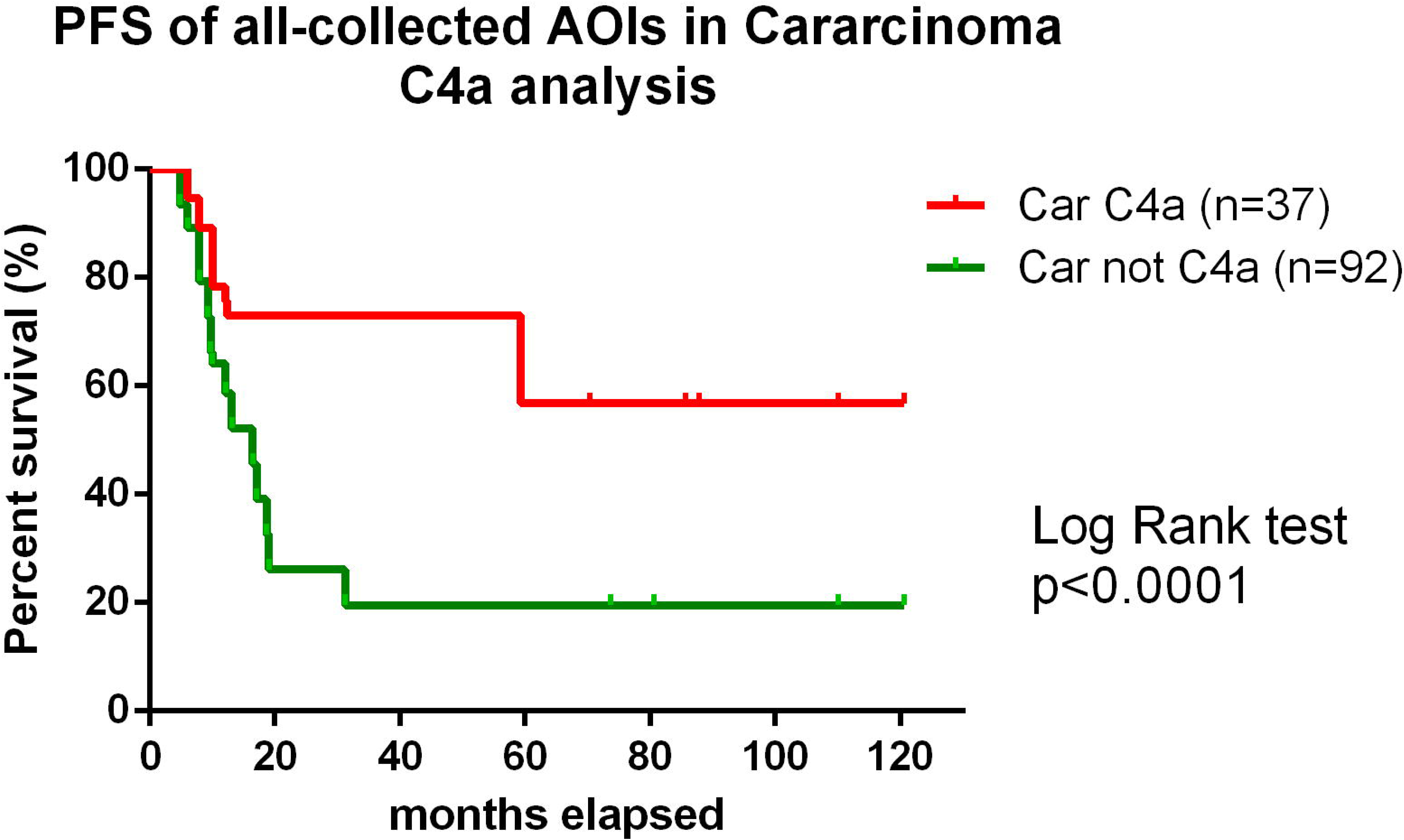

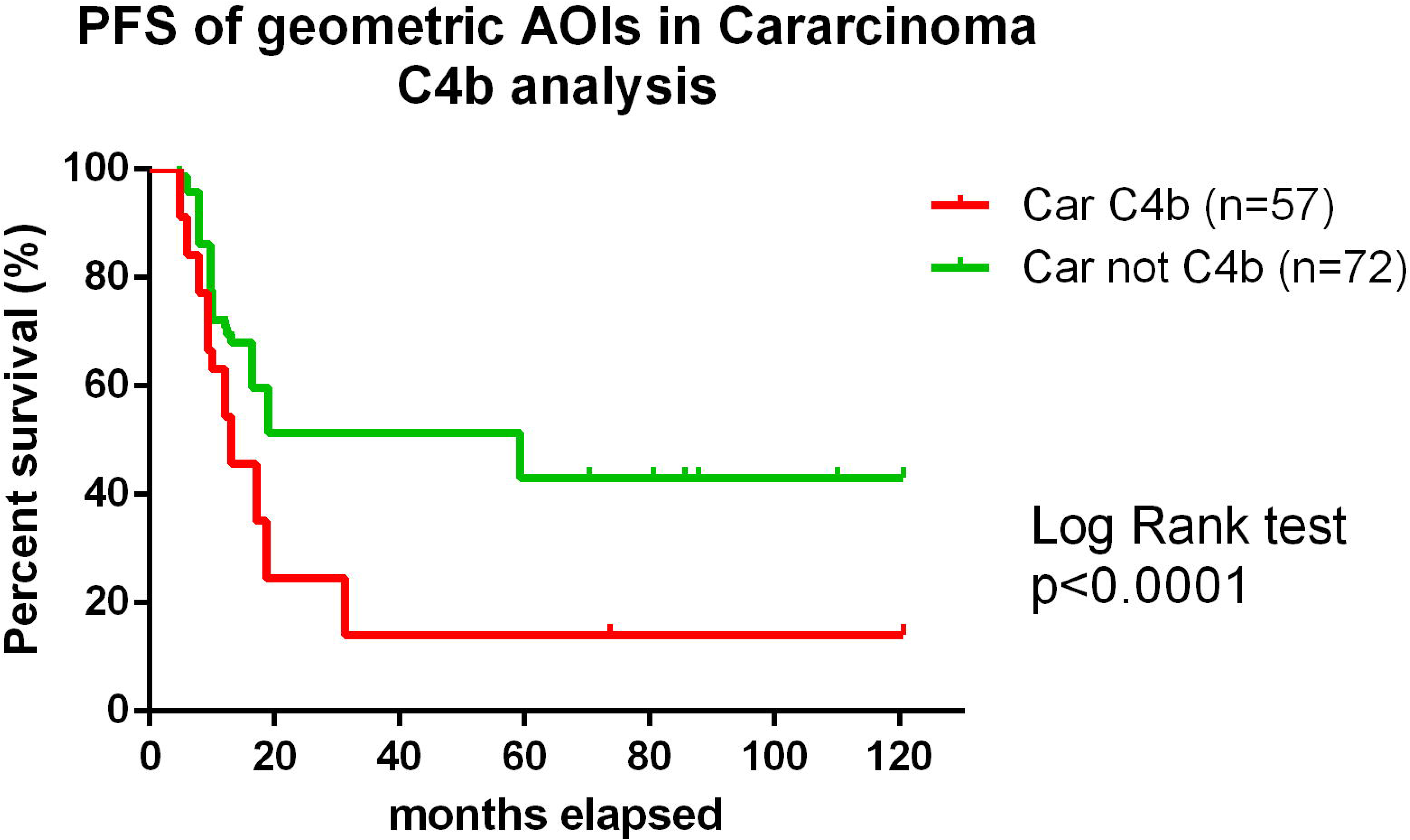

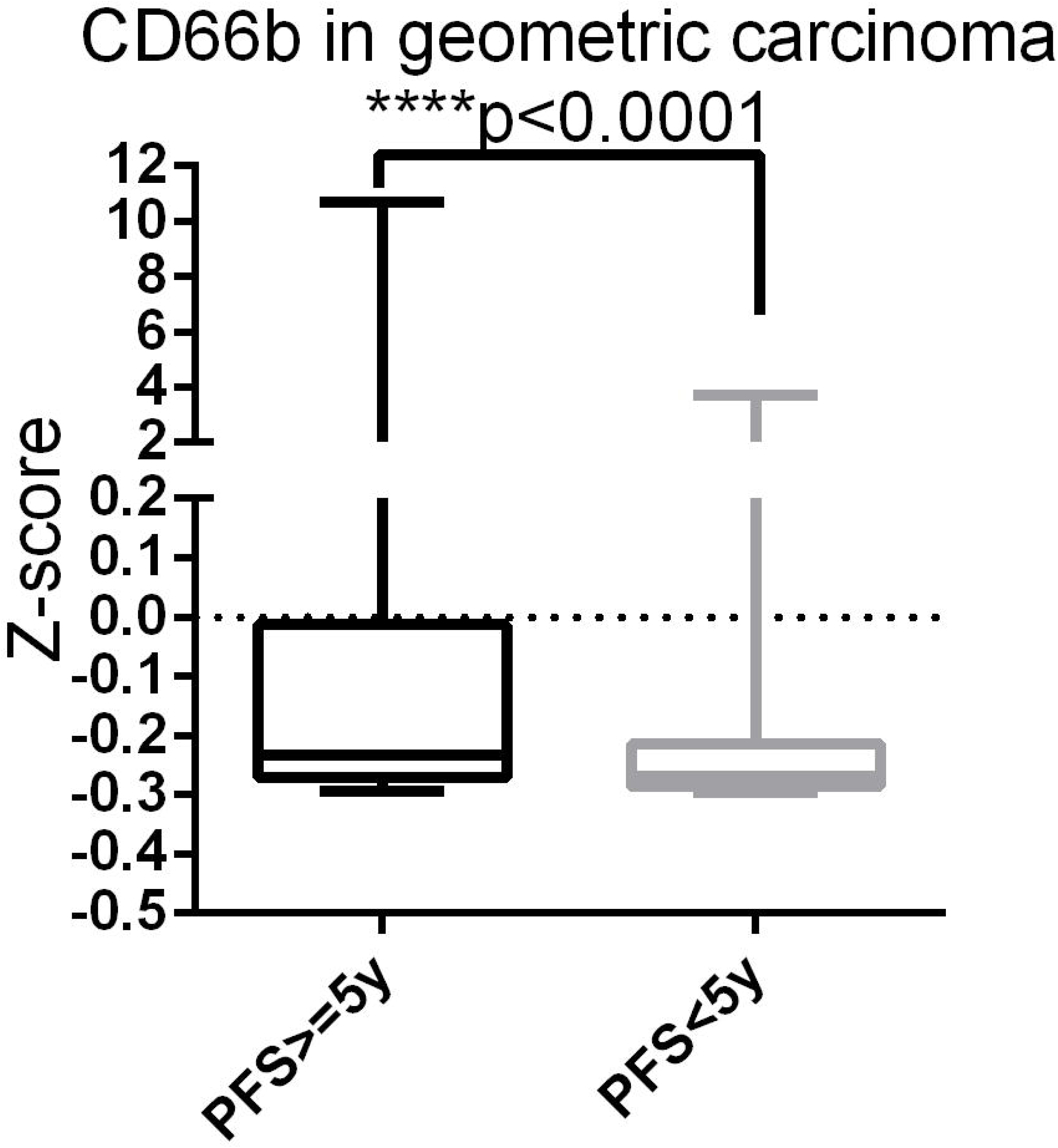

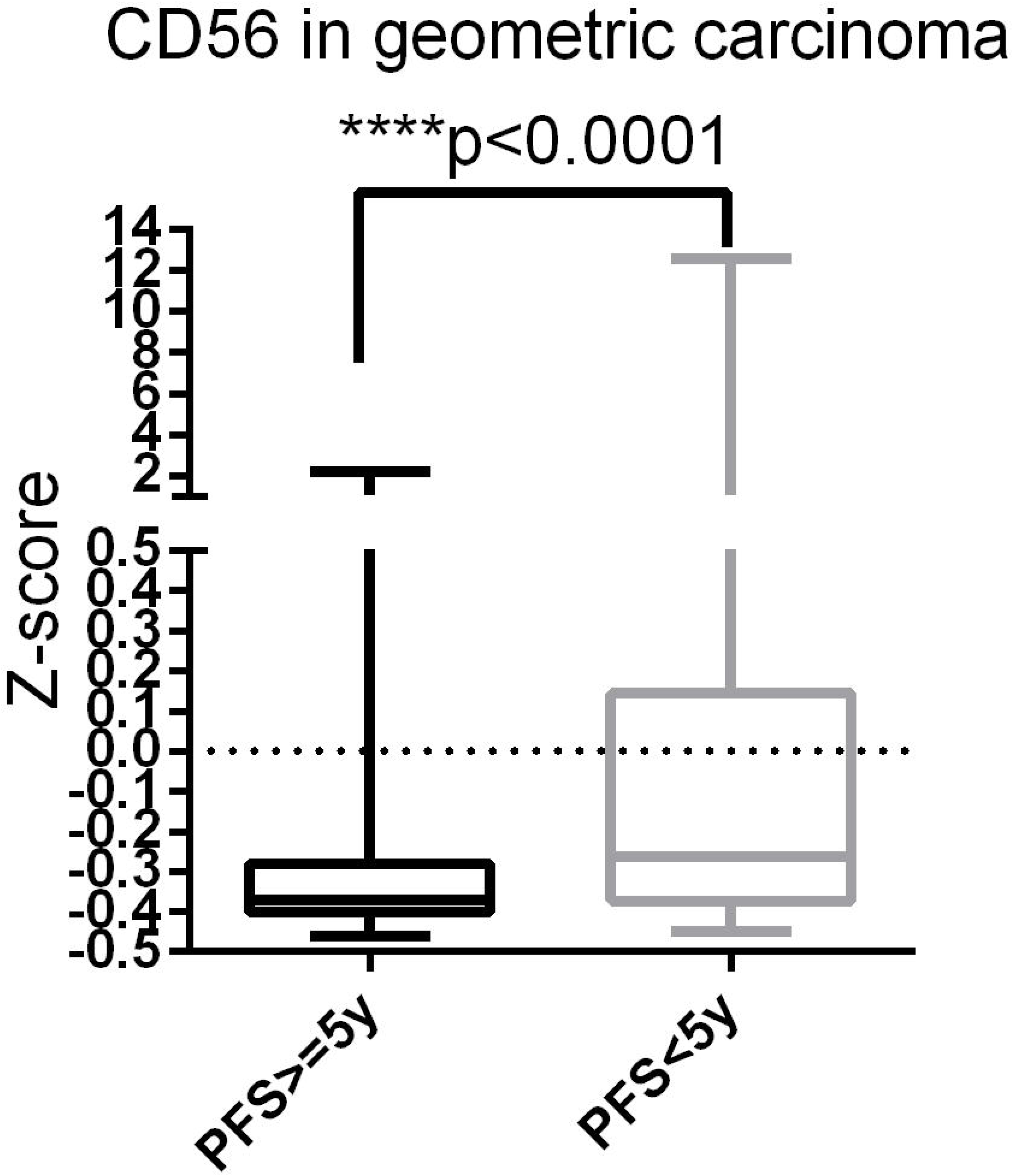

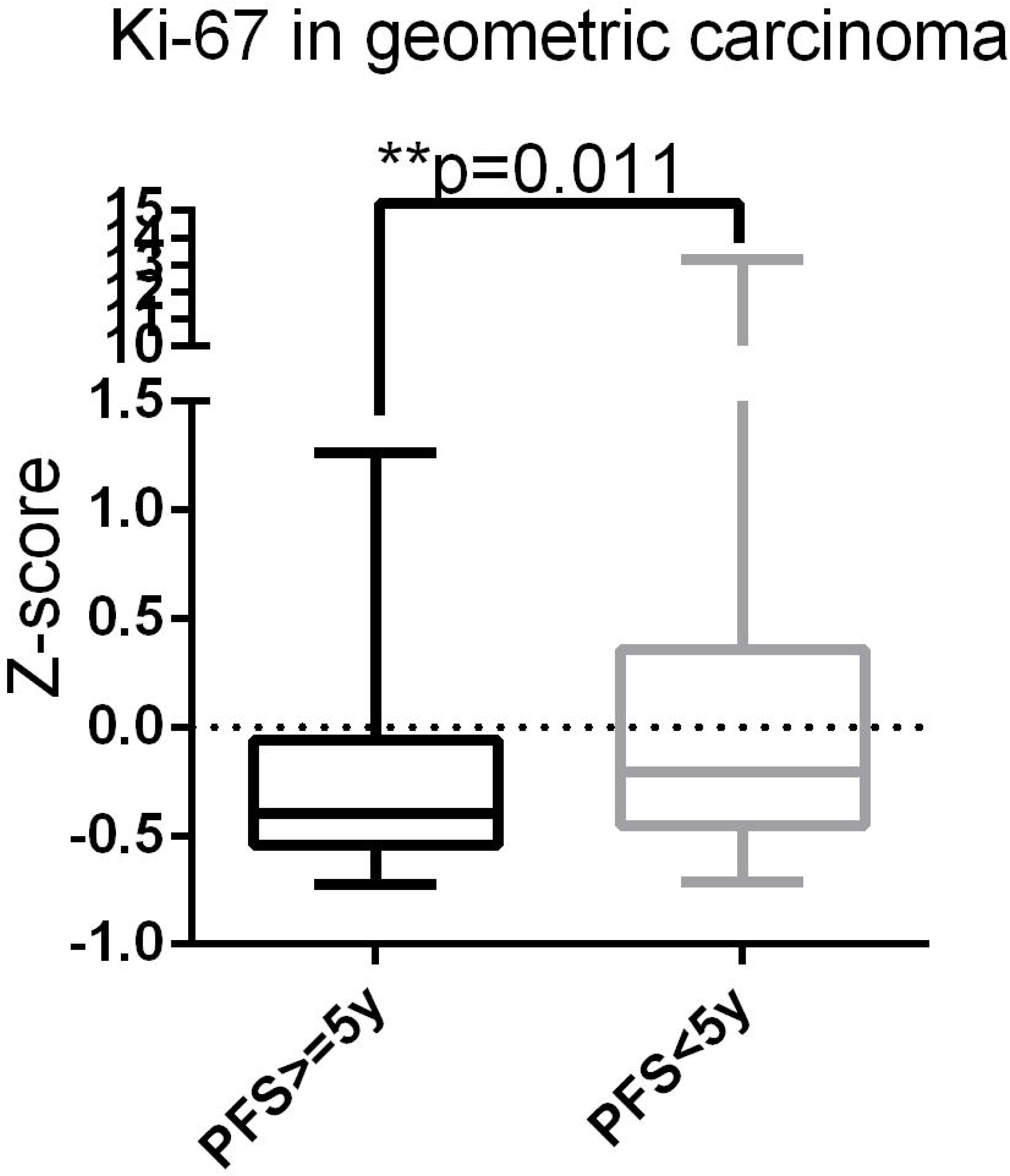

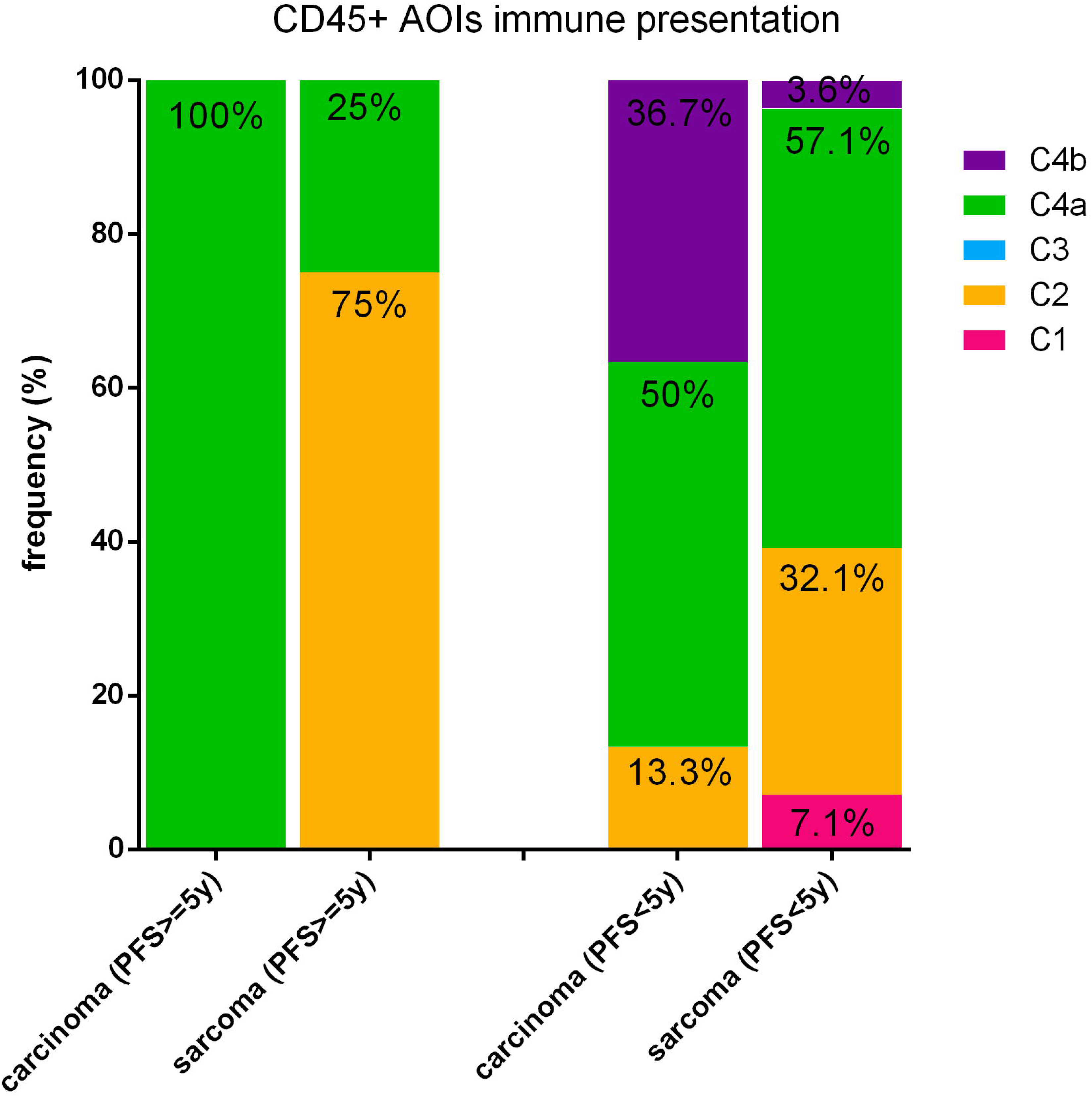

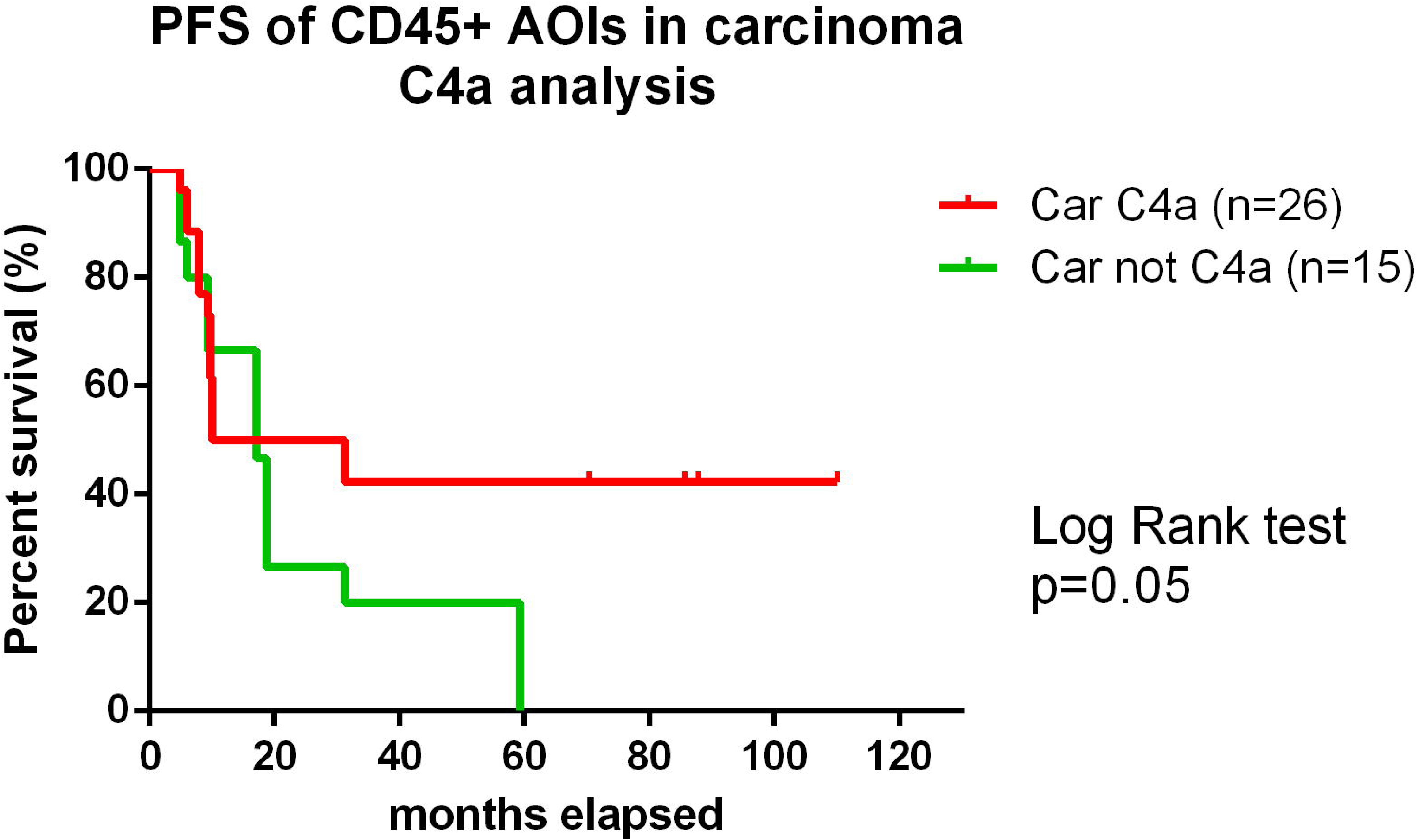

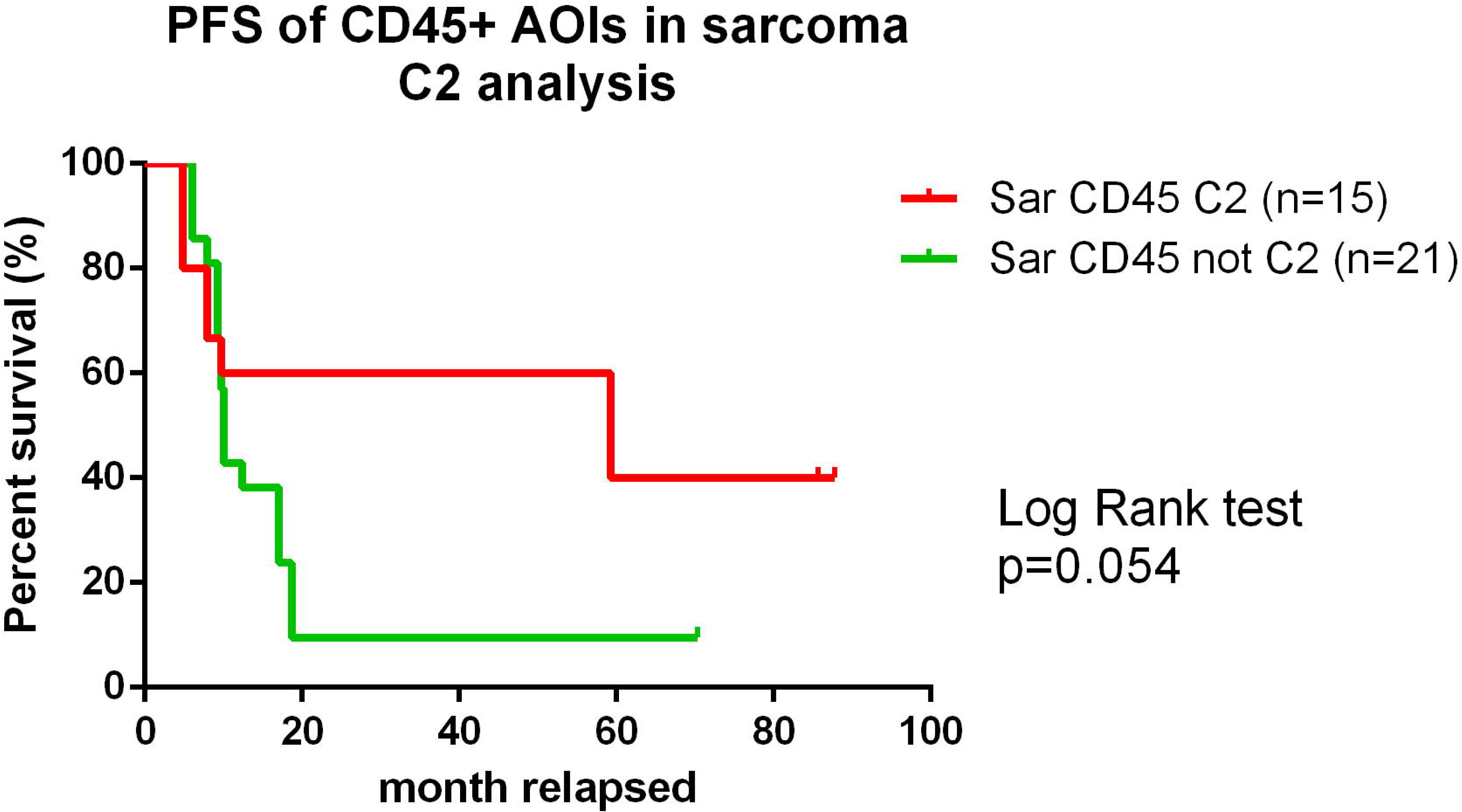

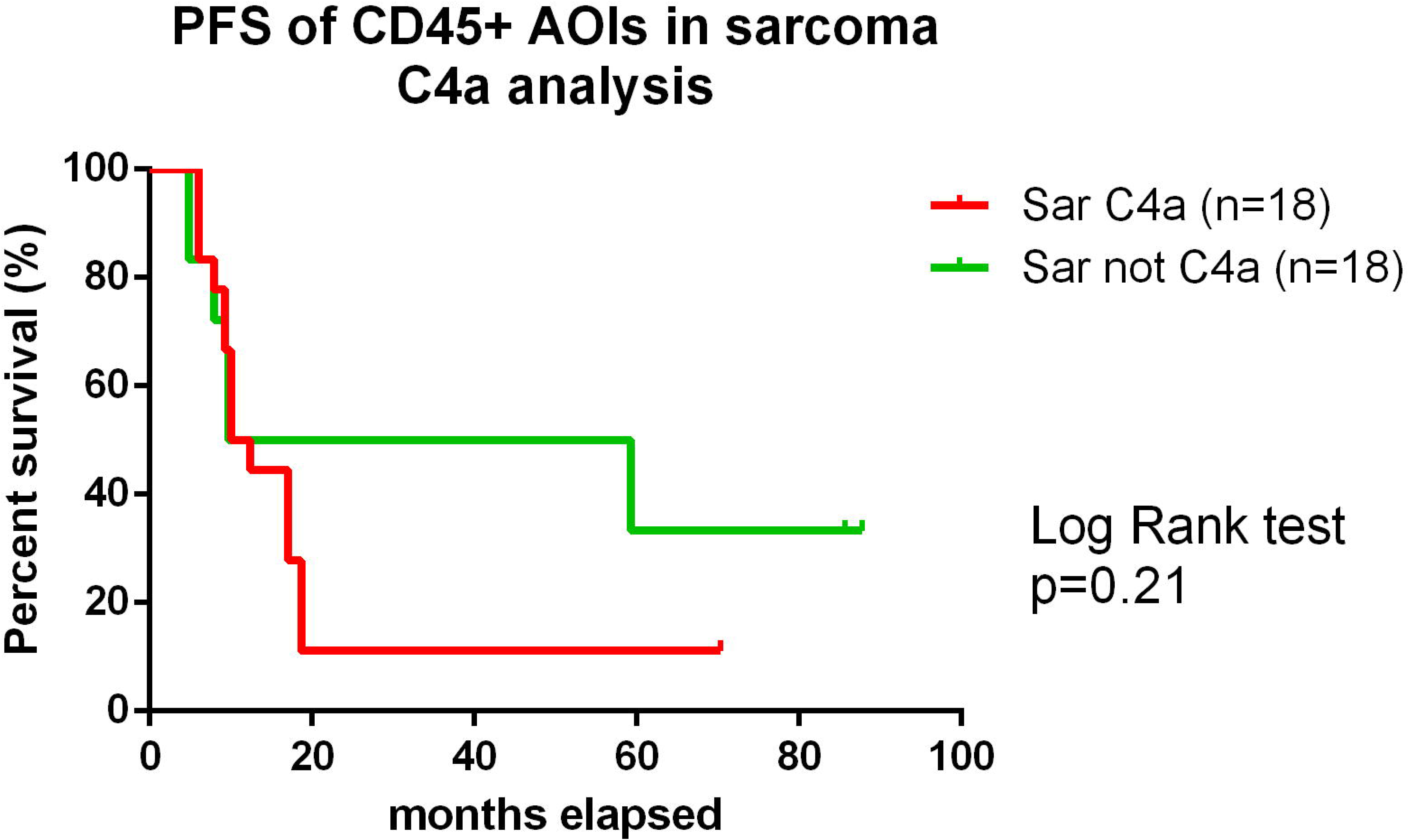

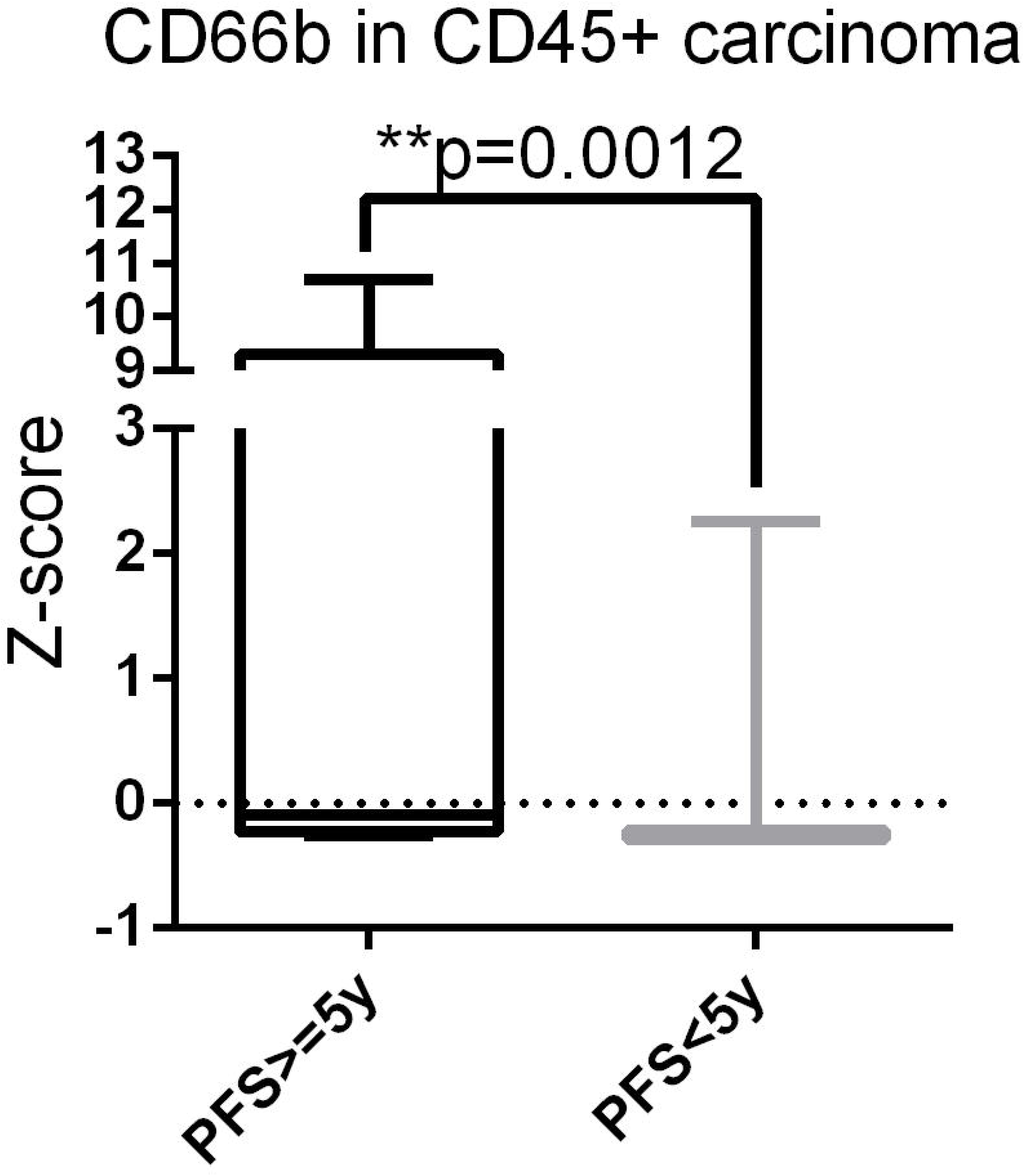

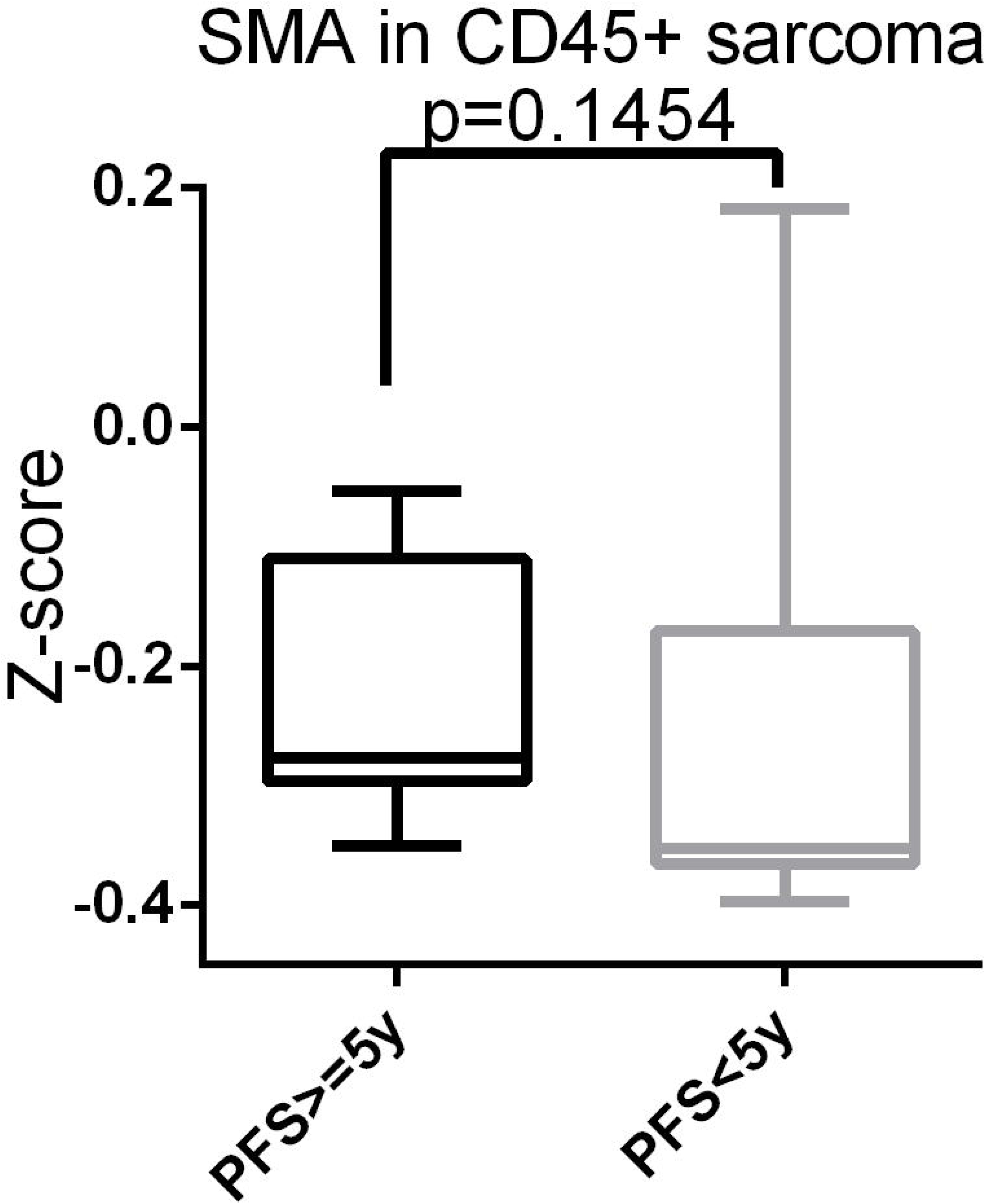

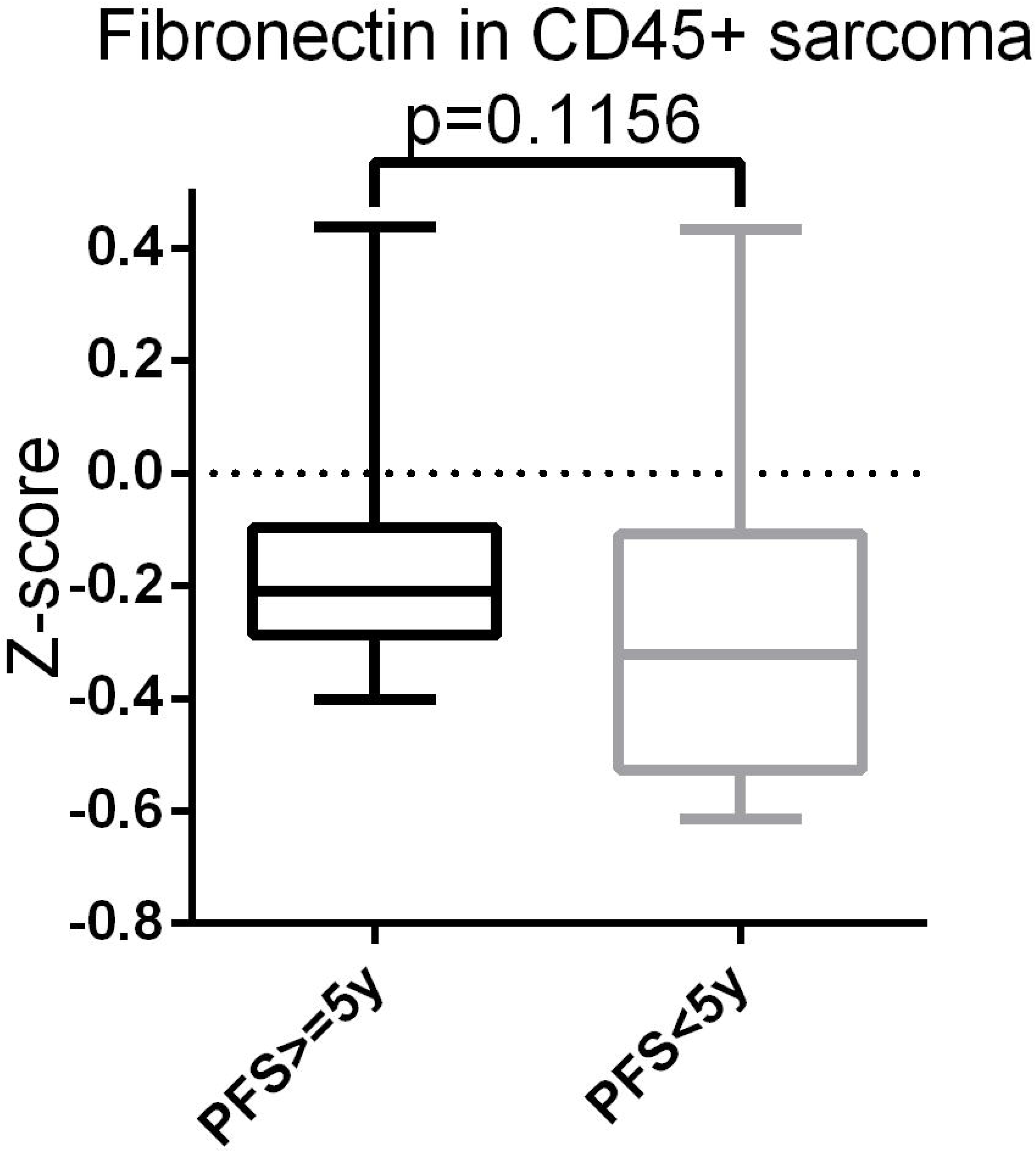

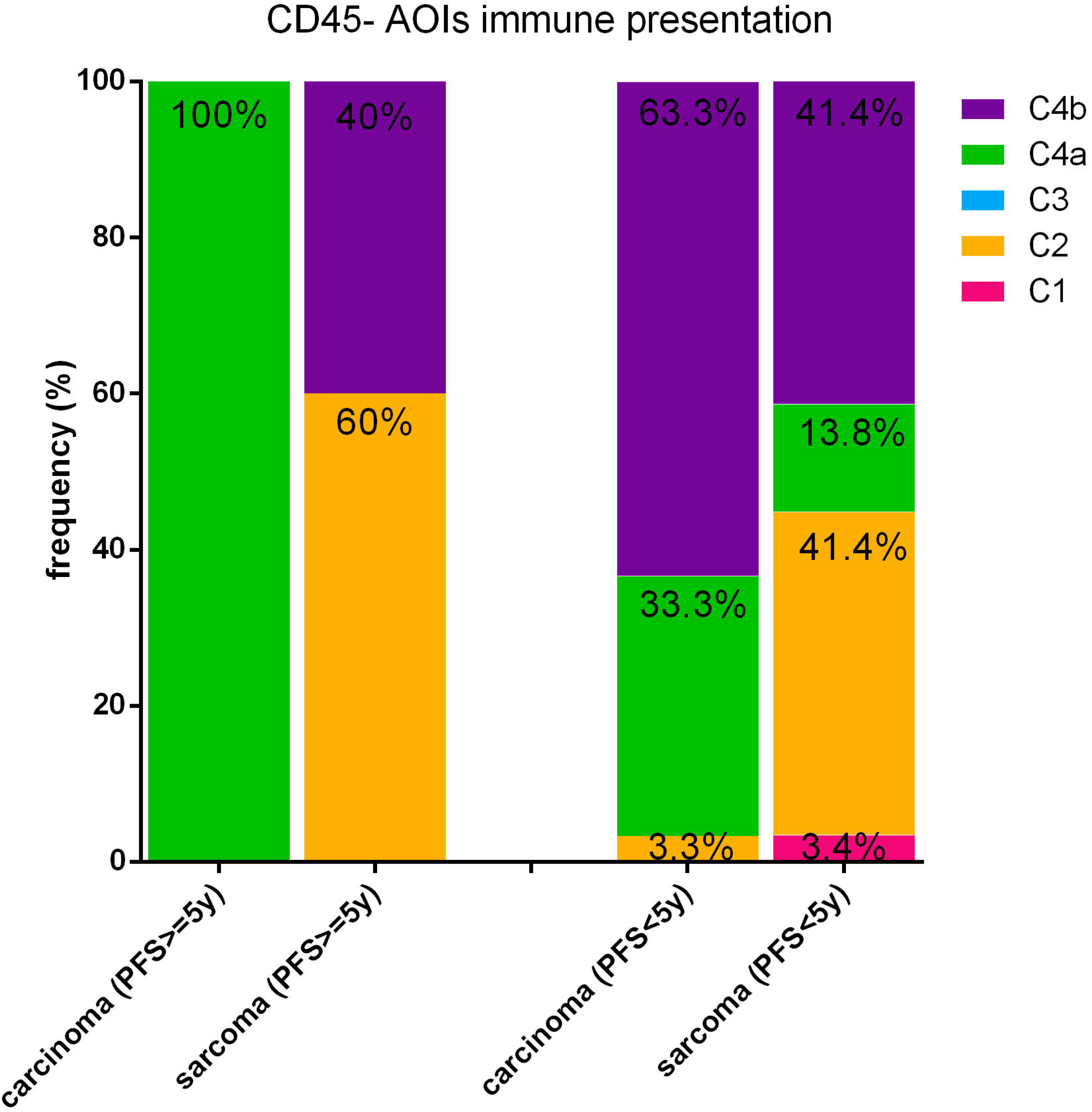

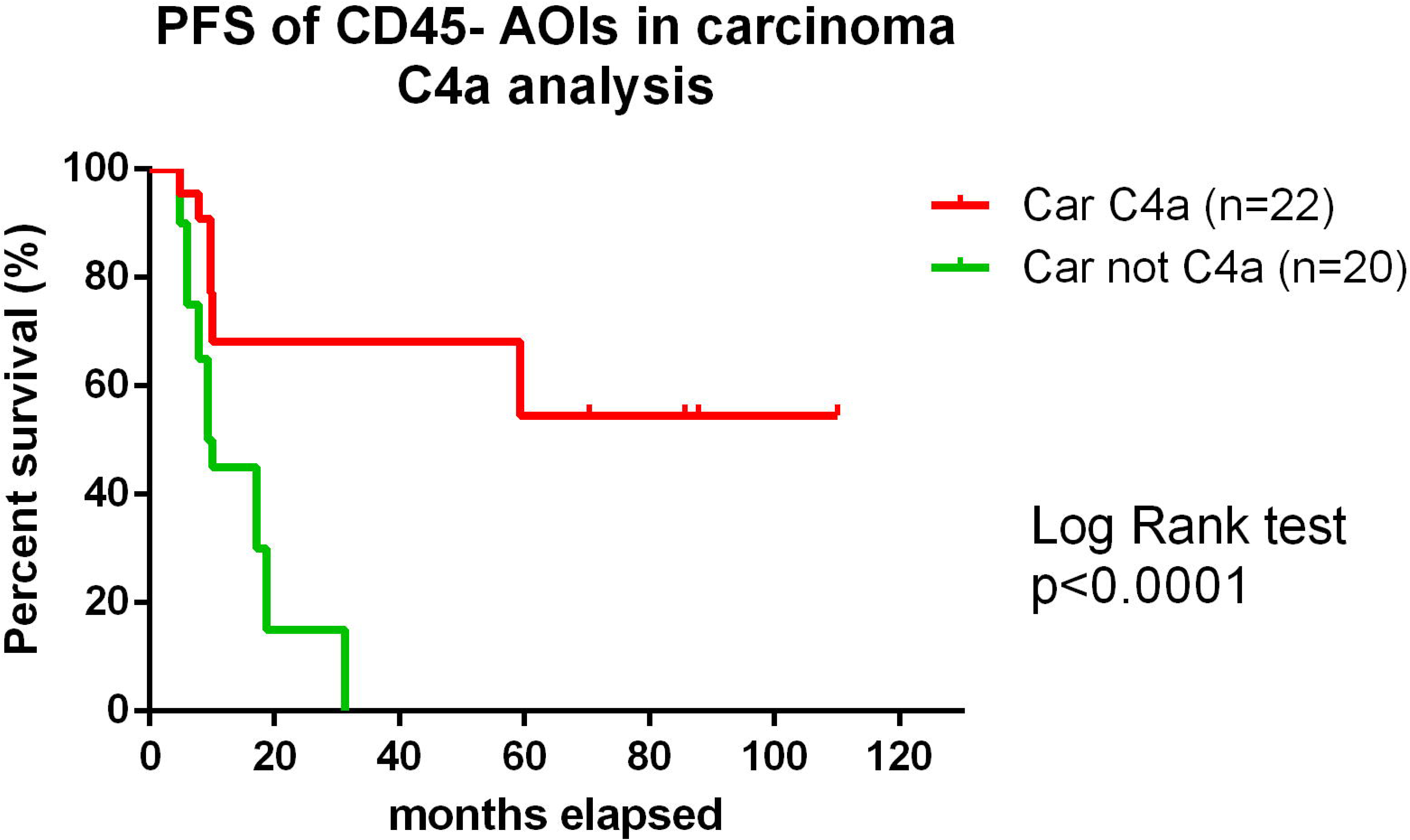

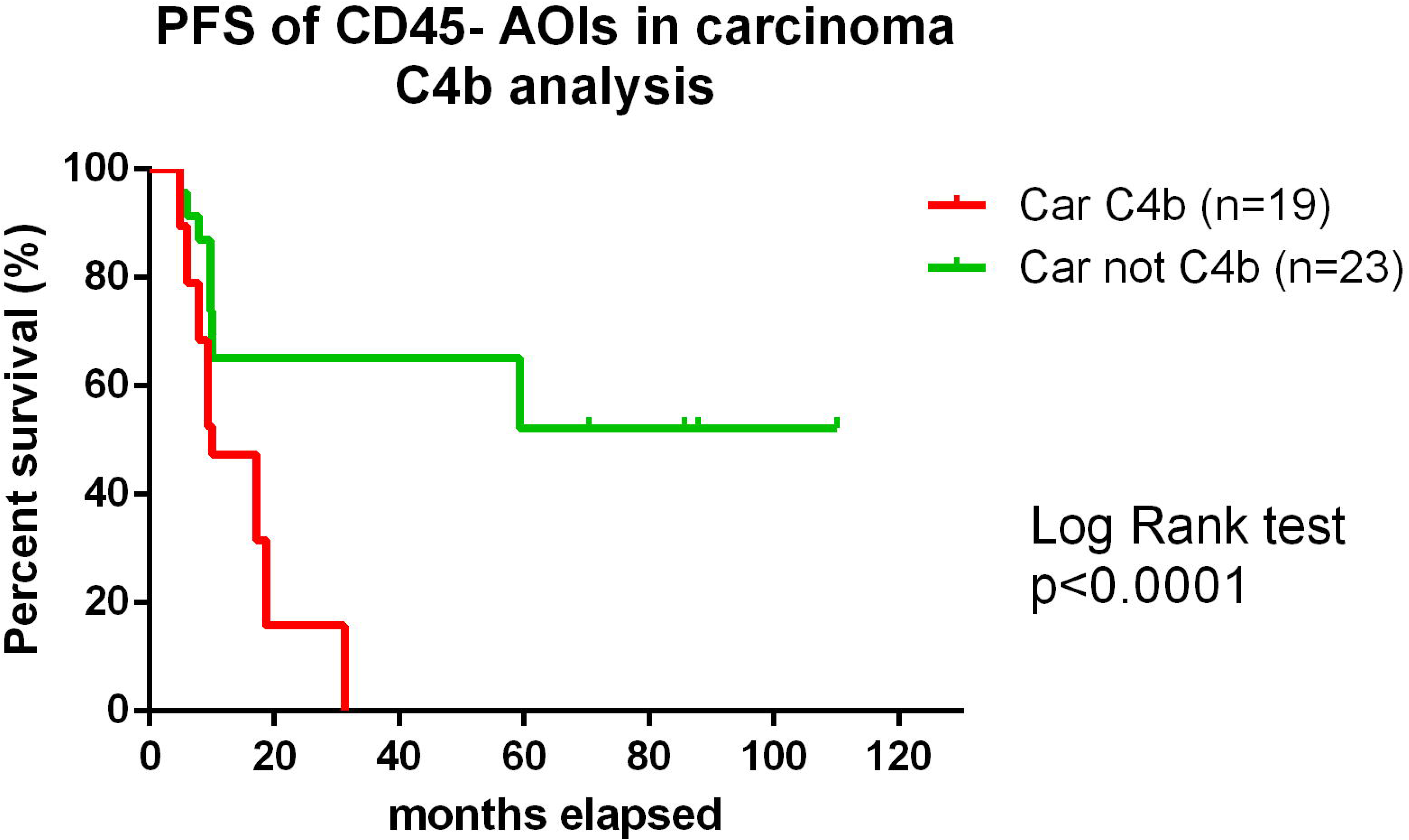

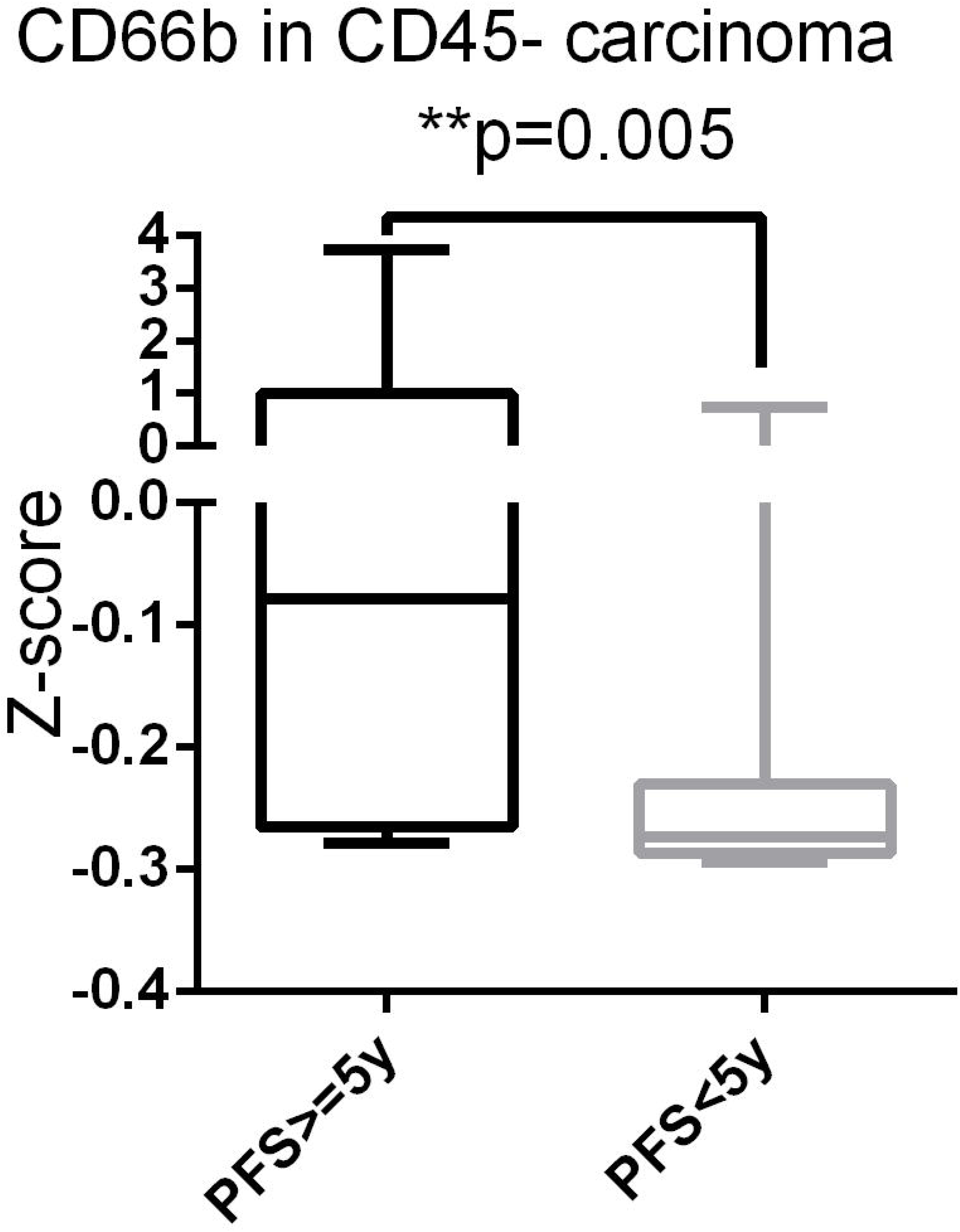

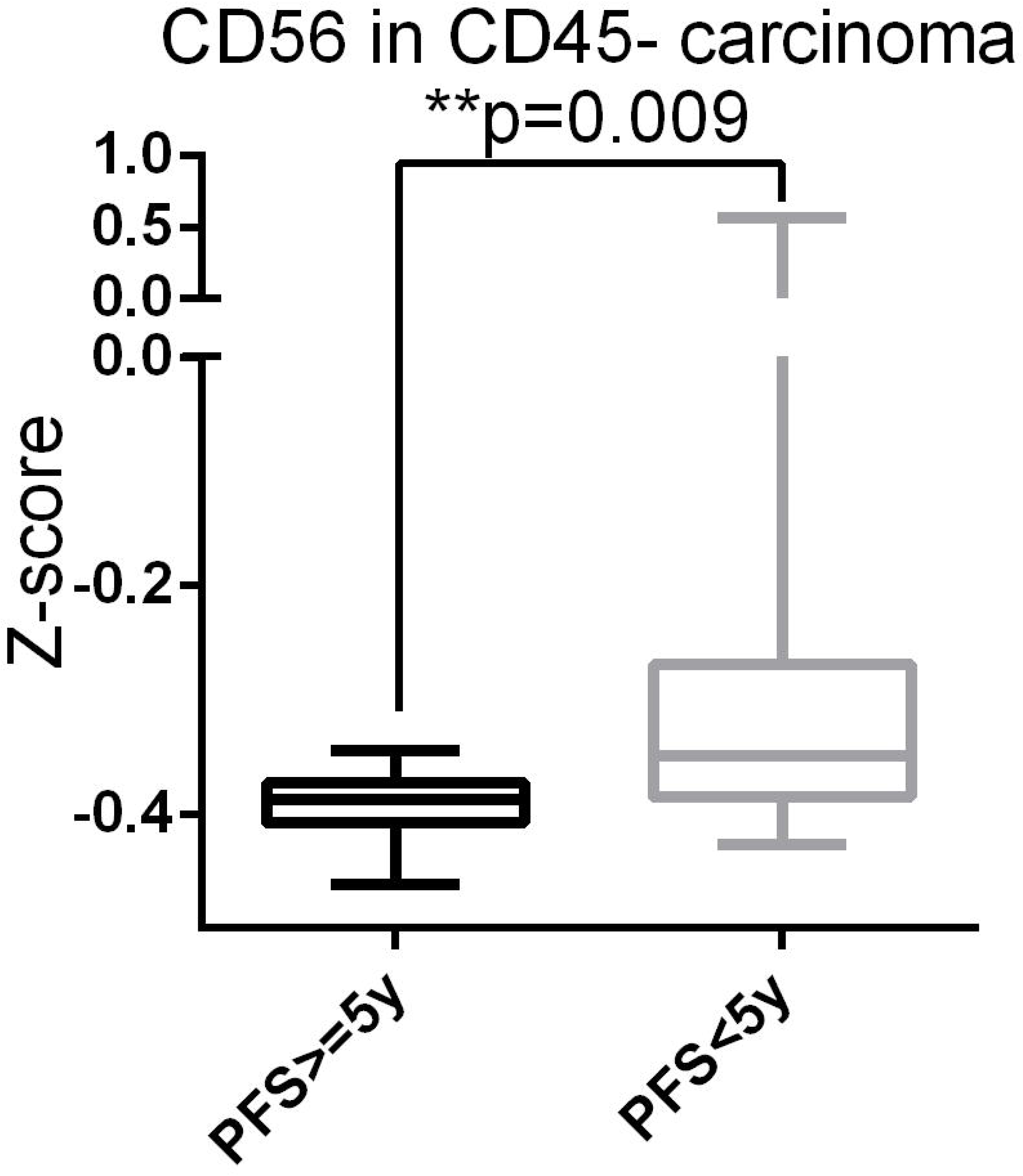

