## Supplementary figures and images for "Inter- and intra-tumor heterogeneous immune presentation in primary uterine carcinosarcoma determined by digital spatial profiling"

### Supplementary Figure 1

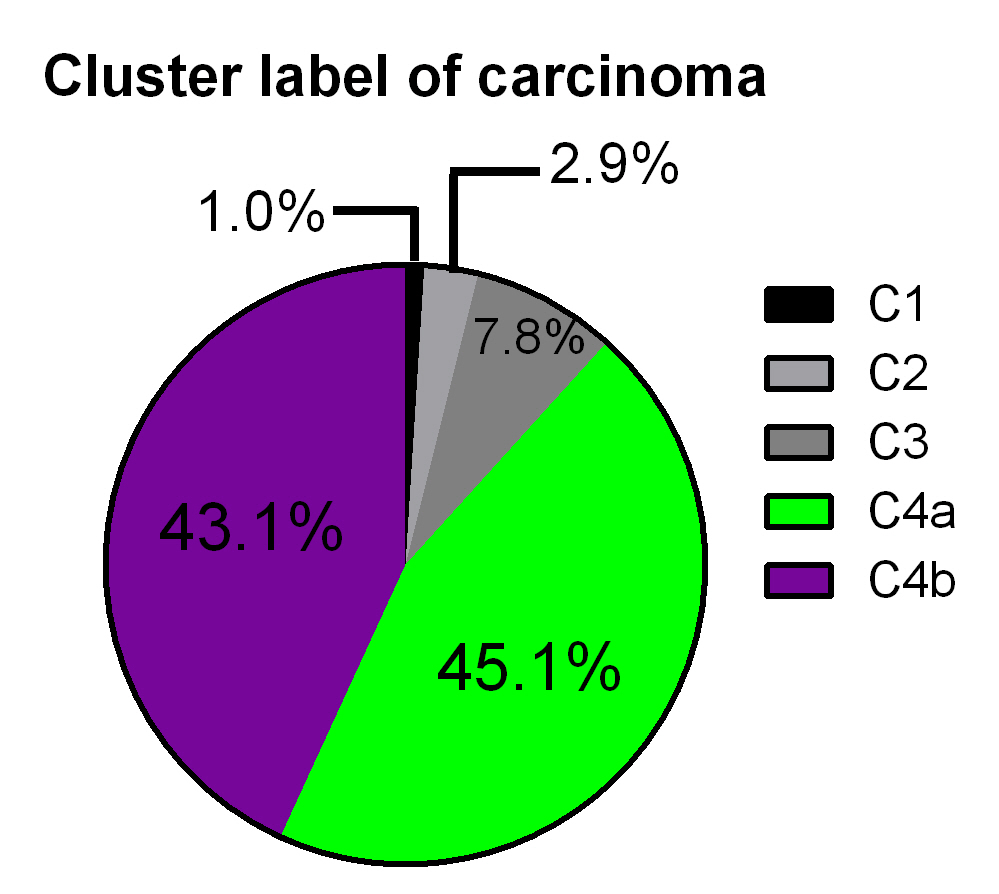

### Supplementary figure 2A

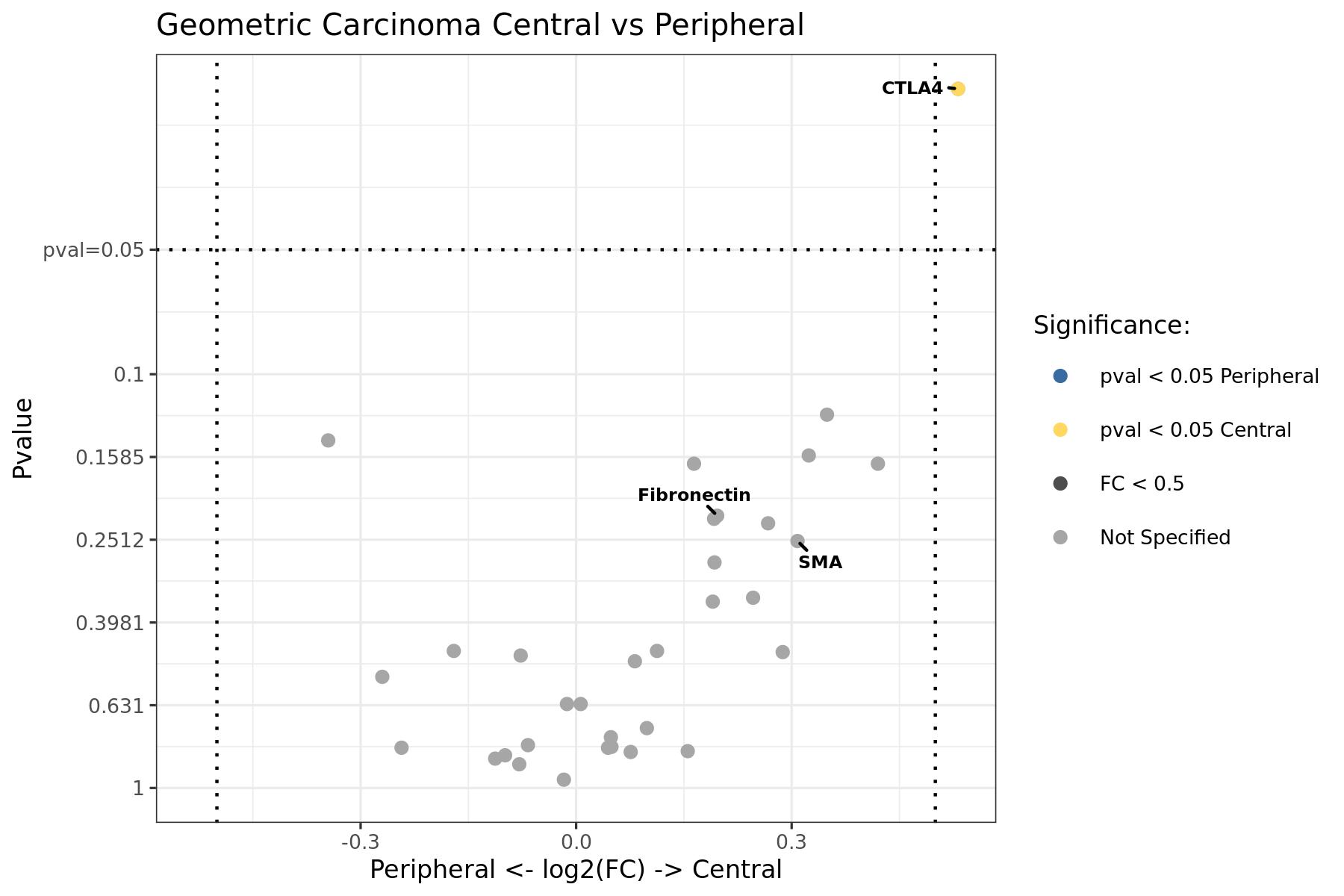

### Supplementary figure 2B

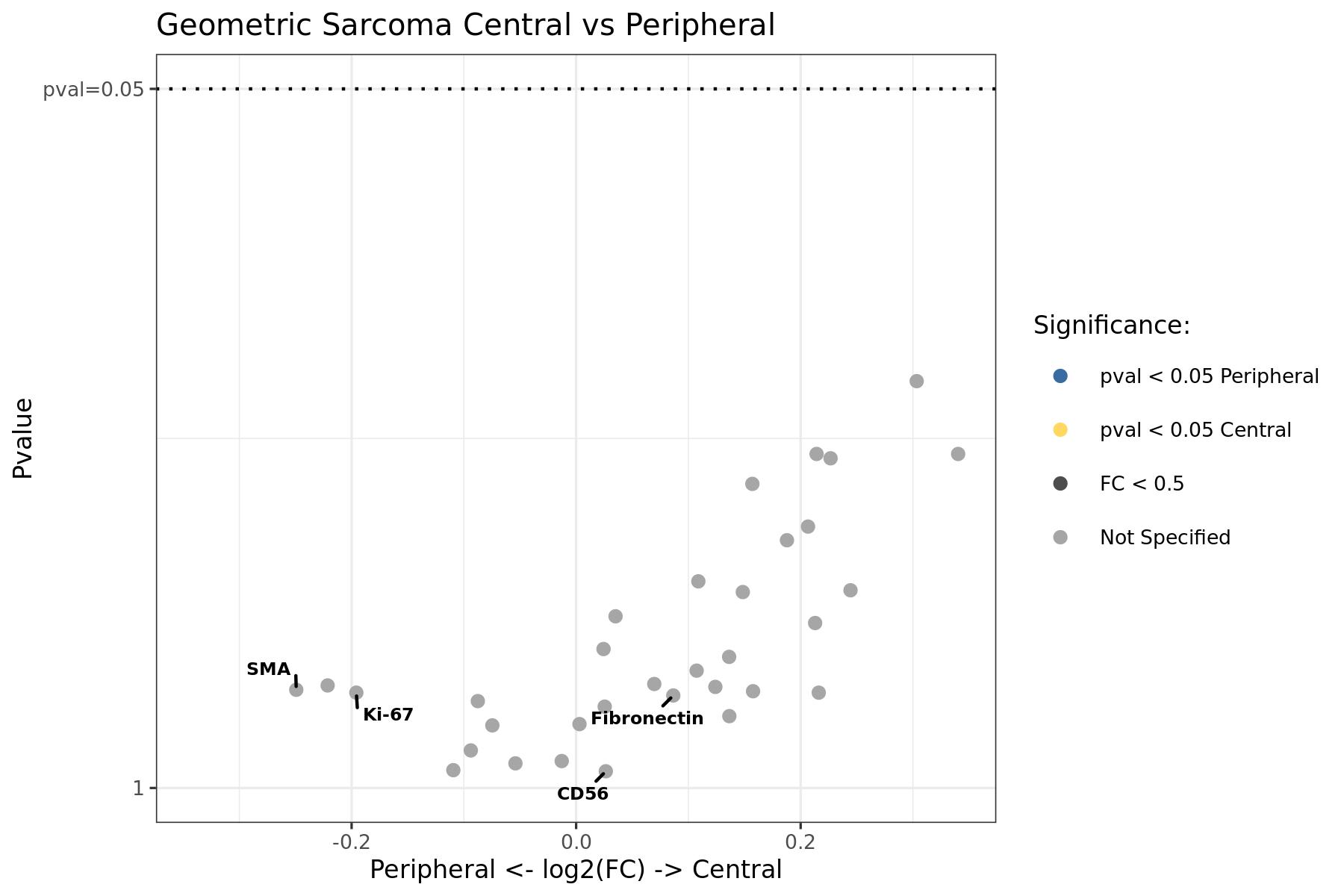

### Supplementary figure 2C

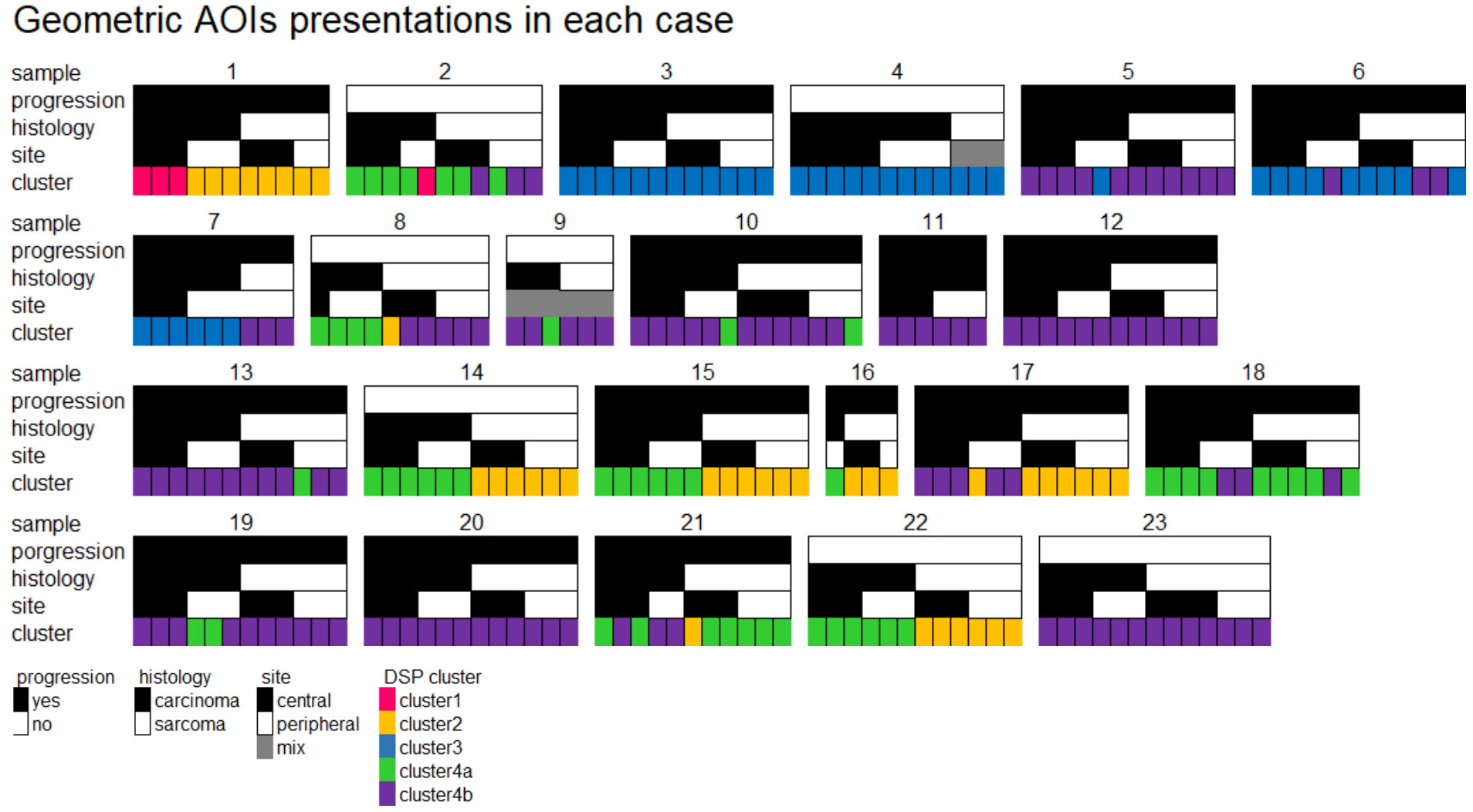

### Supplementary figure 2D

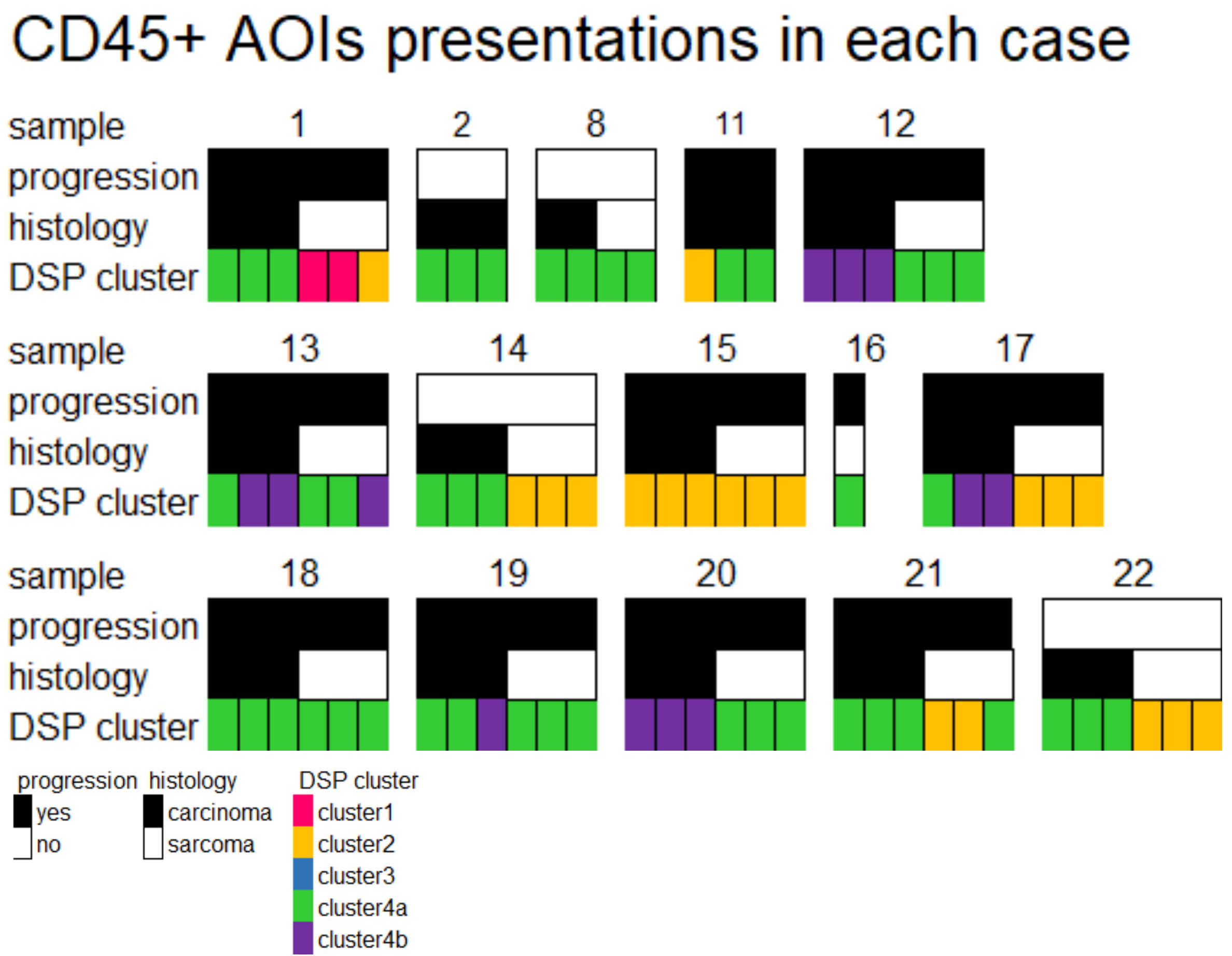

### Supplementary figure 2E

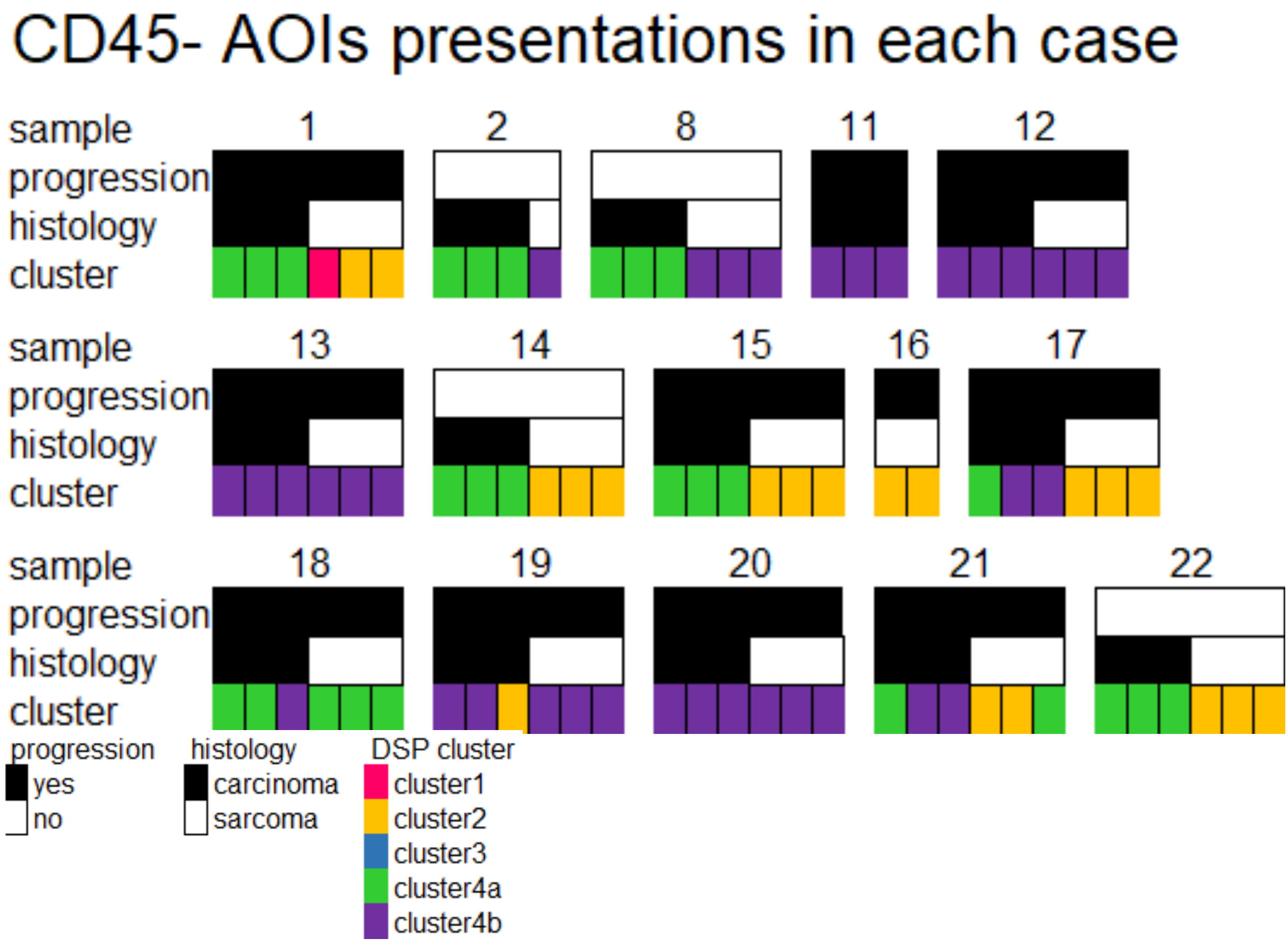
