## Supplemental figure legend for "Inter- and intra-tumor heterogeneous immune presentation in primary uterine carcinosarcoma determined by digital spatial profiling": Supplementary Figure legend.docx

Supplementary Figure 1 The proportion of cluster label within the carcinoma component.

Supplementary Figure 2 (A) A volcano plot showing the differences in immune-related marker levels between central and at peripheral sites of the carcinoma component, focusing on geometric segmentation. (B) A volcano plot showing the differences in immune-related marker levels at the central and peripheral site of the sarcoma component, focusing on geometric segmentation. (C) Geometric AOI immune presentation at different sites of various histology components and disease statuses. (D) The CD45+ AOI immune presentation in different histology components and disease statuses. (E) The CD45- AOI immune presentation in different histology components and disease statuses.
